# Global drivers of obligate mycorrhizal symbionts diversification

**DOI:** 10.1101/2020.07.28.224790

**Authors:** Benoît Perez-Lamarque, Maarja Öpik, Odile Maliet, Ana C. Afonso Silva, Marc-André Selosse, Florent Martos, Hélène Morlon

## Abstract

Analyzing diversification dynamics is key to understanding the past evolutionary history of clades that led to present-day biodiversity patterns. While such analyses are widespread in well-characterized groups of species, they are much more challenging in groups which diversity is mostly known through molecular techniques. Here, we use the largest global database on the small subunit (SSU) rRNA gene of Glomeromycotina, a subphylum of microscopic arbuscular mycorrhizal fungi that provide mineral nutrients to most land plants by forming one of the oldest terrestrial symbioses, to analyze the diversification dynamics of this clade in the past 500 million years (Myr). We perform a range of sensitivity analyses and simulations to control for potential biases linked to the nature of the data. We find that Glomeromycotina tend to have low speciation rates compared to other eukaryotes. After a peak of speciations between 200 and 100 Myr ago, they experienced an important decline in speciation rates toward the present. Such a decline could be at least partially related to a shrinking of their mycorrhizal niches and to their limited ability to colonize new niches. Our analyses identify patterns of diversification in a group of obligate symbionts of major ecological and evolutionary importance and illustrate that short molecular markers combined with intensive sensitivity analyses can be useful for studying diversification dynamics in microbial groups.

## Introduction

Understanding past dynamics of speciation and extinction, as well as the abiotic and biotic factors that modulate the frequency of speciation and extinction events (i.e. diversification rates) is key to understanding the historical processes that shaped present-day biodiversity patterns (Barnosky, 2001; Condamine, Rolland, Höhna, Sperling, & Sanmartín, 2018; Morlon, 2014; Varga et al., 2019) (Barnosky, 2001; Benton, 2009; Chomicki, Kiers, & Renner, 2020; Clarke & Gaston, 2006). While phylogenetic analyses of diversification are widespread in well-characterized groups of species, such as animals and plants (Givnish et al., 2015; Magallón & Sanderson, 2001; Rolland, Condamine, Jiguet, & Morlon, 2014; Upham, Esselstyn, & Jetz, 2019), they are much more challenging in groups which diversity is mostly known through environmental DNA sequences and molecular techniques. In particular, the characterization of poorly cultivable microbial groups such as most bacteria and fungi is often limited to metabarcoding techniques, which consist in the specific amplification and sequencing of a short DNA region (Taberlet, Bonin, Zinger, & Coissac, 2018). One the one hand, these data often render species delineation, phylogenetic reconstruction, and the estimation of global scale diversity highly uncertain, which all affect the phylogenetic inference of diversification dynamics (Lekberg et al., 2018; Moen & Morlon, 2014). On the other hand, it is possible to assess the robustness of phylogenetic diversification analyses to data uncertainty. Given the current limitations of sequencing technologies and the nature of the molecular data available for most microbial groups, using metabarcoding data and performing thorough robustness analyses is one of the only (if not the only) possible approach to analyze their diversification dynamics (Davison et al., 2015; Lewitus, Bittner, Malviya, Bowler, & Morlon, 2018; Louca et al., 2018).

Here we analyze the diversification dynamics of arbuscular mycorrhizal fungi from the subphylum Glomeromycotina. These fungi are obligate symbionts that have been referred to as an “evolutionary cul-de-sac, albeit an enormously successful one” (Malloch, 1987; Morton, 1990). This alludes to their ecological success despite limited morphological and species diversities: they associate with the roots of >80% of land plants, where they provide mineral resources in exchange for photosynthates (Smith & Read, 2008). Present in most terrestrial ecosystems, Glomeromycotina play key roles in plant protection, nutrient cycling, and ecosystem processes (van der Heijden, Martin, Selosse, & Sanders, 2015). Fossil evidence and molecular phylogenies suggest that Glomeromycotina contributed to the emergence of land plants (Feijen, Vos, Nuytinck, & Merckx, 2018; Field, Pressel, Duckett, Rimington, & Bidartondo, 2015; Selosse & Le Tacon, 1998; Strullu-Derrien, Selosse, Kenrick, & Martin, 2018) and coevolved with them for more than 400 million years (Myr)(Lutzoni et al., 2018; Simon, Bousquet, Lévesque, & Lalonde, 1993; Strullu-Derrien et al., 2018).

Glomeromycotina are microscopic soil- and root-dwelling fungi that are hard to differentiate based on morphology and difficult to cultivate without host plant. Although their classical taxonomy is mostly based on the characters of spores and root colonization (Smith & Read, 2008; Stürmer, 2012), Glomeromycotina species delineation has greatly benefited from DNA sequencing (Krüger, Krüger, Walker, Stockinger, & Schüßler, 2012). Experts have defined “virtual taxa” (VT) based on a minimal 97% similarity of a region of the 18S small subunit (SSU) rRNA gene and monophyly criteria (Öpik, Davison, Moora, & Zobel, 2014; Öpik et al., 2010). As for many other pragmatic species delineation criteria, VT have rarely been tested for their biological relevance (Powell, Monaghan, Öpik, & Rillig, 2011), and a consensual system of Glomeromycotina classification is still lacking (Bruns, Corradi, Redecker, Taylor, & Öpik, 2018). Besides the rDNA region, Glomeromycotina remain poorly known genetically: other gene sequences are available for only a few species (James et al., 2006; Lutzoni et al., 2018) and less than 30 complete genomes are currently available (Venice et al., 2020).

Hence, despite the ecological ubiquity and evolutionary importance of Glomeromycotina, large-scale patterns of their diversification dynamics, as well as the factors that correlate with these dynamics, remain poorly known. A previous dated phylogenetic tree of VT found that many speciation events occurred after the last major continental reconfiguration around 100 Myr ago (Davison et al., 2015), suggesting the radiation of Glomeromycotina is not linked to vicariant speciation during this geological event. Indeed, vicariant speciation might only play a minor role in Glomeromycotina diversification, as these organisms have spores that disperse efficiently, promoting gene flow (Bueno & Moora, 2019; Correia, Heleno, da Silva, Costa, & Rodríguez-Echeverría, 2019; Egan, Li, & Klironomos, 2014). Based on the diversity and abundance of Glomeromycotina in tropical grasslands (Read, 1991), it has been suggested (but never tested) that these habitats are diversification hotspots for Glomeromycotina (Pärtel et al., 2017). In this case, the pace of Glomeromycotina diversification through time could be tightly linked to changes in the total area of tropical grasslands. Finally, Glomeromycotina are currently obligate symbionts and their evolutionary history could thus have been largely influenced by their interactions with their host plants (Lutzoni et al., 2018; Sauquet & Magallón, 2018; Zanne et al., 2014). Over the last 400 Myr, land plants have experienced massive extinctions and radiations (Cleal & Cascales-Miñana, 2014; Zanne et al., 2014), adaptations to various ecosystems (Bredenkamp, Spada, & Kazmierczak, 2002; Brundrett & Tedersoo, 2018), and associations with different soil microorganisms (Werner et al., 2018; Werner, Cornwell, Sprent, Kattge, & Kiers, 2014). All these events could have influenced the diversification dynamics of Glomeromycotina, although their relative generalism (Perez-Lamarque, Selosse, Öpik, Morlon, & Martos, 2020; Sanders, 2003; van der Heijden et al., 2015) could buffer this influence.

We aim to characterize the pace of Glomeromycotina diversification in the last 500 Myr and to test the association between diversification rates and a variety of biotic and abiotic factors. We begin by reconstructing several thoroughly sampled phylogenetic trees of Glomeromycotina, considering several criteria of species delineations and uncertainty in phylogenetic reconstructions. We combine this phylogenetic data with paleoenvironmental data and data of current Glomeromycotina geographic distributions, ecological traits, interaction with host plants, and genetic diversity. Finally, we apply a series of birth-death models of cladogenesis to answer specific questions and test hypotheses related to Glomeromycotina diversification: (i) how often do speciation events occur? (ii) were speciation rates relatively constant, or were they higher during specific periods of evolutionary history? and do speciation rates decline through time, as observed for many macroorganisms (Moen & Morlon, 2014)? (iii) are speciation rates positively correlated with past temperature, CO^2^ concentration, and/or land plant diversity? (iv) are present-day speciation rates correlated with geographic distribution, spore size (itself often inversely related to dispersal capacity, Nathan et al., 2008), degree of specialization toward plant species, and genetic diversity? For each of these questions, we thoroughly assess the robustness of our results to uncertainty in the data.

## Material & methods

### Virtual taxa phylogenetic reconstruction

We downloaded the Glomeromycotina SSU rRNA gene sequences from MaarjAM, the largest global database of Glomeromycotina gene sequences updated in June 2019 (Öpik et al., 2010). We reconstructed several Bayesian phylogenetic trees of the 384 *virtual taxa* (VT) from the corresponding representative sequences available in the MaarjAM database (Supplementary Methods 1). We used the full length (1,700 base pairs) SSU rRNA gene sequences from (Rimington et al., 2018) to better align the VT sequences using MAFFT (Katoh & Standley, 2013). We selected the 520 base pair central variable region of the VT aligned sequences and performed a Bayesian phylogenetic reconstruction using BEAST2 (Bouckaert et al., 2014). We set the crown root age at 505 Myr (Davison et al., 2015), which is coherent with fossil data and previous dated molecular phylogenies (Lutzoni et al., 2018; Strullu-Derrien et al., 2018). We also used the youngest (437 Myr) and oldest (530 Myr) crown age estimates from (Lutzoni et al., 2018) in diversification analyses that may be particularly sensitive to absolute dates.

### Delineation into Evolutionary Units (EUs)

We considered several ways to delineate Glomeromycotina species based on the SSU rRNA gene. In addition to the VT species proxy, we delineated Glomeromycotina *de novo* into evolutionary units (EUs) using a monophyly criterion and 5 different thresholds of sequence similarity ranging from 97 to 99%. We gathered Glomeromycotina sequences of the SSU rRNA gene from MaarjAM, mainly amplified by the primer pair NS31–AML2 (variable region) (Lee, Lee, & Young, 2008; Simon, Lalonde, & Bruns, 1992) (dataset 1, Supplementary Table 1). There were 36,411 sequences corresponding to 27,728 haplotypes. We first built a phylogenetic tree of these haplotypes and then applied to this tree our own algorithm of EU delineation (R-package RPANDA (Morlon et al., 2016; R Core Team, 2020)) that traverses the tree from the root to the tips, at every node computes the average similarity of all sequences descending from the node, and collapses the sequences into a single EU if their sequence dissimilarity is lower than a given threshold (Supplementary Methods 2). In other words, Glomeromycotina sequences are merged into the same EU if they form a monophyletic clade and if they are on average more similar than the sequence similarity threshold. Finally, we performed Bayesian phylogenetic reconstructions of the EUs using BEAST2, using the same crown ages as above (Supplementary Methods 1).

### Coalescent-based species delineation analyses

Finally, we considered the Generalized Mixed Yule Coalescent method (GMYC) (Fujisawa & Barraclough, 2013; Pons et al., 2006), a species delineation approach that does not require specifying an arbitrary similarity threshold. GMYC estimates the time *t* in a reconstructed calibrated tree that separates species diversification (Yule process – before *t*) and intraspecific differentiation (coalescent process – after *t*). GMYC is too computationally intensive to be applied to the 36,411 SSU sequences; we used it here on three clades of manageable size (the family Claroideoglomeraceae; the order Diversisporales; and an early-diverging clade composed of the orders Archaeosporales and Paraglomerales) to (i) investigate whether the SSU gene evolves fast enough to accumulate substitutions between Glomeromycotina speciation events (Bruns et al., 2018) and (ii) evaluate the biological relevance of the VT and various EUs delineations. For each clade, we reconstructed Bayesian phylogenetic trees of haplotypes (Supplementary Methods 1). We then ran GMYC analyses (splits R-package (Ezard, Fujisawa, & Barraclough, 2009)) on each of these trees and evaluated the support of the GMYC model compared to a null model in which all tips are assumed to be different species, using a likelihood ratio test (LRT). If the LRT supports the GMYC model, different SSU haplotypes belong to the same Glomeromycotina species, *i.e.* the SSU rRNA gene has time to accumulate substitutions between Glomeromycotina speciation events.

### Total diversity estimates

We evaluated how thoroughly sampled our species-level Glomeromycotina phylogenetic trees are by estimating the total number of VT and EUs using rarefaction curves and the Bayesian Diversity Estimation Software (BDES (Quince, Curtis, & Sloan, 2008)) (Supplementary Methods 3). The BDES estimates the total number of species by extrapolating a sampled taxa abundance distribution at global scale (Quince et al., 2008).

### Additional molecular markers

We explored the possibility to carry some of our analyses using two other molecular markers: the large subunit (LSU) rRNA gene and the ITS2 region. We downloaded the Glomeromycotina LSU database of Delavaux et al. (2020) as well as the LSU sequences available in MaarjAM. We obtained a total 2,044 sequences that we aligned using MAFFT and TrimAl. We retained the 1,760 unique haplotypes, reconstructed the phylogenetic tree of the LSU sequences using BEAST2 and used the resulting calibrated tree to delineate Glomeromycotina LSU units with the GMYC model (same pipeline as above). We similarly downloaded the Glomeromycotina ITS dataset of Lekberg et al. (2018). We tried to align them but confirmed that the ITS sequences of Glomeromycotina are very difficult to align, making them unsuitable for phylogenetic reconstruction and subsequent diversification analyses (Supplementary Fig. 1).

### Diversification analyses

Unless specified differently, our diversification analyses were performed using the SSU rRNA gene. In order to account for various sources of uncertainties in the SSU rRNA data, we replicated all our diversification analyses across different species delineations, phylogenetic reconstructions and dating, and total diversity estimates. For each species delineation criterion, we obtained a consensus tree and selected 12 trees equally spaced in 4 independent Bayesian chains, hereafter referred to as the replicate trees. When the 12 trees were not sufficient to conclude, we used 100 replicate trees.

We estimated lineage-specific speciation rates using ClaDS, a Bayesian diversification model that accounts for speciation rate heterogeneity by modeling small rate shifts at speciation events (Maliet, Hartig, & Morlon, 2019). At each speciation event, the descending lineages inherit new speciation rates sampled from a log-normal distribution with an expected value log[*α*×λ] (where λ represents the parental speciation rate and *α* is a trend parameter) and a standard deviation *σ*. We considered the model with constant turnover ε (*i.e.* constant ratio between extinction and speciation rates; *ClaDS2*) and ran a newly-developed ClaDS algorithm based on data augmentation techniques which enables us to estimate mean rates through time (Maliet & Morlon, 2022). We ran ClaDS with 3 independent chains, checked their convergence using a Gelman-Rubin diagnostic criterion (Gelman & Rubin, 1992), and recorded lineage-specific speciation rates. We also recorded the estimated hyperparameters (*α*, σ, ε) and the value m=α×exp(σ^2^/2), which indicates the general trend of the rate through time (Maliet et al., 2019). We replicated these analyses using the LSU gene.

In addition, we applied CoMET (TESS R-package (Höhna, May, & Moore, 2016; May, Höhna, & Moore, 2016)), another diversification approach that does not consider rate variation across lineages, but models temporal shifts in speciation and extinction rates affecting all lineages simultaneously. CoMET is a piecewise-constant model in a Bayesian framework. We chose the Bayesian priors according to maximum likelihood estimates from TreePar (Stadler, 2011), disallowed or not mass extinction events, and ran the MCMC chains until convergence (minimum effective sample sizes of 500).

We also fitted a series of time-dependent and environment-dependent birth-death diversification models using RPANDA (Condamine, Rolland, & Morlon, 2013; Morlon et al., 2016) to confirm the observed temporal trends and test the influence of temperature, pCO^2^, and land plant fossil diversity on rates of Glomeromycotina speciation. For the time-dependent models, we considered models with constant or exponential variation of speciation rates through time and null or constant extinction rates (*fit_bd* function). As extinction is notoriously hard to estimate from reconstructed phylogenies (Rabosky, 2016), we tested the robustness of the inferred temporal trend in speciation when fixing arbitrarily high levels of extinction (Supplementary Methods 4). For the environment-dependent models, we considered an exponential dependency of the speciation rates with the environmental variable (env), *i.e.* speciation rate=b*exp(a*env), where a and b are two parameters estimated by maximum likelihood (*fit_env* function). Best-fit models were selected based on the corrected Akaike information criterion (AICc), considering that a difference of 2 in AICc indicates that the model with the lowest AICc is better. We replicated these analyses using the LSU gene.

The influence of temperature was tested on the complete Glomeromycotina phylogenetic trees, using estimates of past global temperature (Royer, Berner, Montañez, Tabor, & Beerling, 2004). We also carried a series of simulation analyses to test the robustness of our temperature-dependent results (Supplementary Methods 5). The influence of pCO^2^ (Foster, Royer, & Lunt, 2017) and of land plant fossil diversity was tested starting from 400 Myr ago, as these environmental data are not available for more ancient times. For these analyses we sliced the phylogenies at 400 and 200 Myr ago, and applied the diversification models to the sliced sub-trees larger than 50 tips. Estimates of land plant diversity were obtained using all available Embryophyta fossils from the Paleobiology database (https://paleobiodb.org) and using the shareholder quorum subsampling method (Supplementary Methods 6; (Alroy, 2010)).

We considered missing species in all our diversification analyses by imputing sampling fractions, computed as the number of observed VT or EUs divided by the corresponding BDES estimates of global Glomeromycotina diversity (Table 1). We used a global sampling fraction for all Glomeromycotina, as the main Glomeromycotina clades had a similar sampling fraction (Supplementary Table 2). To assess the robustness of our results to global diversity estimates, we replicated all diversification analyses using a range of lower sampling fractions (from 90% to 50%, *i.e.* assuming that only that percentage of the global Glomeromycotina species diversity is in fact represented in our dataset).

**Table 1:** Estimation of the total diversity of Glomeromycotina: Estimated sampling fraction using the Bayesian Diversity Estimation Software (BDES; Quince et al., 2008) for the different species delineations (VT or EU) assuming a Sichel species abundance distribution. The estimated number of units corresponds to the median value and we indicated the 95% confidence interval. We indicated the sampling fractions for each delineation, computed as the number of observed VT or EUs divided by the corresponding BDES estimates of global Glomeromycotina diversity. The Sichel distribution was selected compared to other distributions (log-normal, log-Student, and inverse gaussian) based on lowest deviance information criterion (DIC).

| Species delineation | Observed number of units | Estimated number of units | Sampling fraction | 95% confidence interval |  |
| --- | --- | --- | --- | --- | --- |
|  |  |  |  | Lower boundary | Upper boundary |
| VT | 384 | 403 | 95% | 391 (98%) | 422 (91%) |
| EU97 | 182 | 187 | 97% | 183 (99%) | 194 (94%) |
| EU97.5 | 340 | 357 | 95% | 348 (98%) | 370 (92%) |
| EU98 | 641 | 677 | 95% | 663 (97%) | 695 (92%) |
| EU98.5 | 1,190 | 1,268 | 94% | 1,247 (95%) | 1,293 (92%) |
| EU99 | 2,647 | 2,852 | 93% | 2,817 (94%) | 2,890 (92%) |

### Testing for correlates of present-day Glomeromycotina speciation rates

To further investigate the potential factors correlating with Glomeromycotina speciation rates, we assessed the relationship between lineage-specific estimates of present-day speciation rates (obtained with the ClaDS analyses) and characteristics of each Glomeromycotina taxonomic unit, *i.e.* VT or EUs.

First, to assess the effect of specialization on speciation rates, we characterized Glomeromycotina relative niche width using a set of 10 abiotic and biotic variables recorded in MaarjAM database for each Glomeromycotina unit. In short, among a curated dataset containing Glomeromycotina sequences occurring only in natural ecosystems (dataset 2; Supplementary Table 2; Perez-Lamarque et al., 2020), for each Glomeromycotina unit, we reported the number of continents, ecosystems, climatic zones, biogeographic realms, habitats, and biomes where it was sampled, as well as its number of plant partners, their phylogenetic diversity, and its centrality in the plant-fungus bipartite network, and performed a principal component analysis (PCA; Supplementary Methods 7). For Glomeromycotina units represented by at least 10 sequences, we tested whether these PCA coordinates reflecting Glomeromycotina niche widths were correlated with the present-day speciation rates using both linear mixed-models (not accounting for phylogeny) or MCMCglmm models (Hadfield, 2010). For MCMCglmm, we assumed a Gaussian residual distribution, included the fungal phylogenetic tree as a random effect, and ran the MCMC chains for 1,300,000 iterations with a burn-in of 300,000 and a thinning interval of 500.

Next, we tested the relationship between speciation rates and geographic characteristics of Glomeromycotina units. To evaluate the effect of latitude on speciation rates, we associated each Glomeromycotina unit with its set of latitudes and used similar MCMCglmm with an additional random effect corresponding to the Glomeromycotina unit. To account for inhomogeneous sampling along the latitudinal gradient, we re-ran the model on jackknifed datasets (we re-sampled 1,000 interactions per slice of latitude of twenty degrees). Similarly, we tested the effect of climatic zone and habitat on speciation rates.

Then, to assess the effect of dispersal capacity on speciation rates, we evaluated the relationship between spore size and speciation rate for the few (*n*=32) VT that contain sequences of morphologically characterized Glomeromycotina isolates (Davison et al., 2018). We gathered measures of their average spore length (Davison et al., 2018) and tested their relationship with speciation rate by using a phylogenetic generalized least square regression (PGLS).

Finally, as a first attempt at connecting Glomeromycotina macroevolutionary diversification to microevolutionary processes, we measured intraspecific genetic diversities across Glomeromycotina units. For each Glomeromycotina unit containing at least 10 sequences, we computed genetic diversity using Tajima’s estimator (Tajima, 1983)(θπ; Supplementary Methods 8). Using similar statistical tests as above, we investigated the correlation of Glomeromycotina genetic diversity with speciation rate, niche width, geographic characteristics, and spore size. We tested the robustness of the results to the minimal number of sequences per Glomeromycotina unit (10, 15, or 20) used to compute genetic diversity and to perform the PCA.

These statistical models were replicated on the different phylogenetic trees (consensus or replicates) for each delineation and we reported p-values (*P*) corresponding to two-sided tests.

### Simulation analyses

The use of a short and slowly evolving gene such as the central region of the SSU rRNA gene to delineate species may lead to an artificial lumping of species into the same unit that would reduce the number of phylogenetic branching events toward the present and result in a biased inference of temporal diversification dynamics, including an artifactual detection of a diversification slowdown (Moen & Morlon, 2014). We used simulations mimicking the evolution of the SSU rRNA gene as Glomeromycotina diversified to quantify this potential bias.

We simulated the diversification of a clade of species in the last 505 Myr, according to two scenarios: (i) constant speciation rate and no extinction and (ii) constant speciation and extinction rates (Supplementary Figure 2a). To model intraspecific differentiation, we added intraspecific splits on these simulated species trees by grafting coalescent events at each tip: for each species, we uniformly sampled between 2 and 15 individuals and we considered that all these individuals had to coalesce before the last speciation event; the age of the coalescent tree within each species was uniformly sampled between 0 and the age of the last speciation event (with a maximum of 30 Myr). We used the functions *pbtree* and *rcoal* from the R-packages phytools and ape (Paradis, Claude, & Strimmer, 2004; Revell, 2012) to simulate the species phylogenies and the intraspecific coalescences respectively. We used two net diversification rates (r=0.010 and r=0.015) for simulating the species phylogenies, in order to reach a total number of species similar to that obtained with our empirical data when using the VT and EU99 delineations, respectively. Next, we simulated the evolution of short 520 bp DNA sequences on the obtained trees, using the function *simulate_alignment* (R-package HOME; Perez-Lamarque & Morlon, 2019). We used a substitution rate of 0.001 event per Myr and only 25% of variable sites, which resulted in an alignment that mimicked the Glomeromycotina SSU rDNA alignment. We performed 10 simulations per scenario. For each of these simulations we kept the unique haplotypes at present and applied the same pipelines as above, using the EU99 species delineation criteria: after delineating the EU99 units, we reconstructed the EU99 phylogenetic trees, ran the ClaDS analyses on these trees, and recorded mean estimated speciation rates at present and 50, 100, and 150 Myr ago.

## Results

### Glomeromycotina species delineations & phylogenetic reconstructions

We automatically delineated Glomeromycotina into evolutionary units (EU) using a monophyly criterion and several thresholds of SSU rRNA sequence similarity (from 97% to 99%). The EU97.5 and EU98 delineations (obtained using a threshold of 97.5% and 98% respectively) provided a number of Glomeromycotina units (340 and 641) relatively comparable to the 384 currently recognized *virtual taxa* (VT), while the EU97 delineation had much less units (182). Conversely, the EU98.5 and EU99 delineations yielded a much larger number of Glomeromycotina units (1,190 and 2,647). These numbers obtained with the EU98.5 and EU99 delineations were consistent with the numbers obtained using GMYC analyses, which delineate species-like units based on detecting when splitting events in the haplotype tree start to follow branching patterns consistent with intra-specific differentiations (*i.e.* coalescent patterns) instead of speciation events (i.e. birth-death patterns; Supplementary Tables 3, 4, & 5). The GMYC results therefore support the idea that some VT might lump together several cryptic species (Bruns et al., 2018)(Supplementary Note 1), and that a 98.5 or 99% similarity threshold is more relevant for Glomeromycotina species delineation. In addition, the GMYC model is significantly supported over the model where all SSU rRNA haplotypes correspond to a different species (GMYC LRT: *P<*0.05; Supplementary Fig. 3), with on average 10 SSU haplotypes per species-like unit, and a mean intraspecific sequence similarity of 99% (Supplementary Table 5 & Supplementary Fig. 3). This indicates that the region of the SSU marker used to characterized Glomeromycotina evolves fast enough to accumulate substitutions between Glomeromycotina speciation events, meaning that it is an informative (although not perfect) marker for delineating Glomeromycotina species-like units. In comparison, the same pipeline carried on the LSU database delineated only 181 GMYC units, suggesting that it was much less complete than the SSU database. We replicated the subsequent diversification analyses using the LSU region, even though we put more trust in our results using the SSU database given the incompleteness of the LSU database.

Rarefaction curves as well as BDES (Bayesian Diversity Estimation Software) and Chao2 estimates of diversity suggested that more than 90% of the total Glomeromycotina diversity is represented in our SSU dataset regardless of the delineation threshold (Fig. 1, Table 1, Supplementary Tables 5 & 6), which is consistent with the proportion of new Glomeromycotina units detected in recent studies (Sepp et al., 2019).

**Figure 1:**
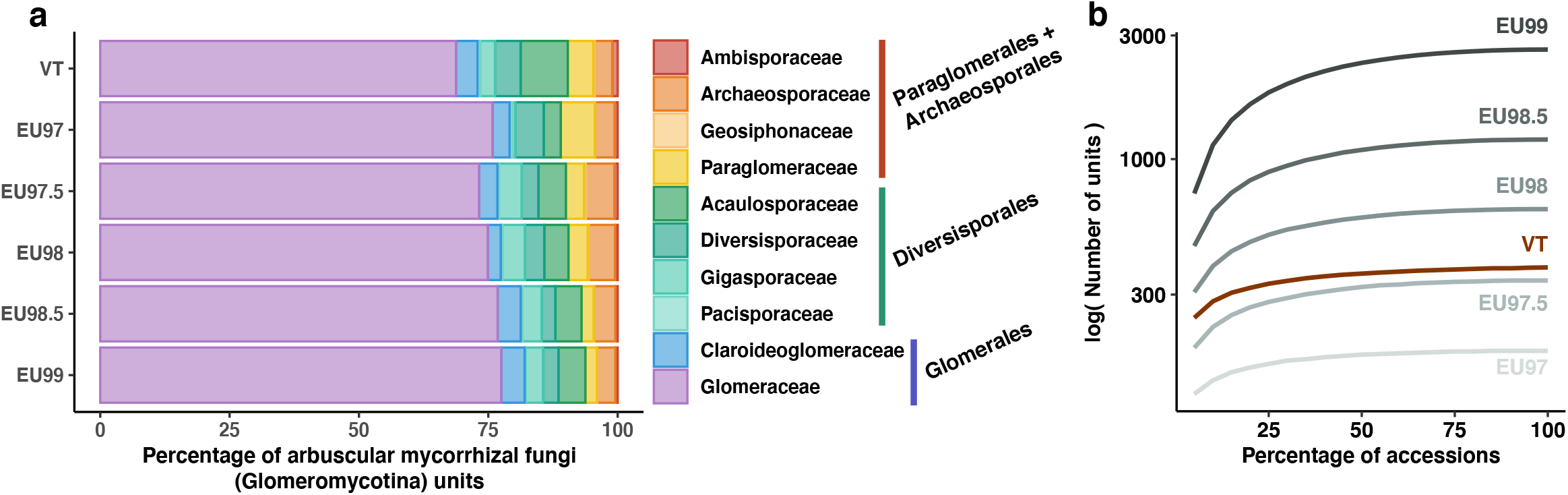
Molecular-based species delineations of Glomeromycotina (arbuscular mycorrhizal fungi) give consistent results and indicate a nearly complete sampling. We compared the *virtual taxa* (VT) delineation from (Öpik et al., 2010) with newly-developed automatic delineations into *evolutionary units* (EUs) based on an average threshold of similarity and a criterion of monophyly. **(a)** The proportion of Glomeromycotina units (VT or EUs) in each Glomeromycotina family reveals constant proportions across delineations, although Glomeraceae tend to be relatively less abundant compared with the other Glomeromycotina family in the VT delineation. The main Glomeromycotina orders are indicated on the right of the charts: Paraglomerales + Archaeosporales, Diversisporales, and Glomerales (Glomeraceae + Claroideoglomeraceae). **(b)** Rarefaction curves indicating the number of Glomeromycotina units as a function of the percentage of sampled Glomeromycotina accession revealed that the Glomeromycotina sampling in MaarjAM is close to saturation for all delineations (VT or EUs). Rarefactions were performed 100 times every 5 percent and the median of the 100 replicates is represented here.

The reconstructed Bayesian phylogenetic trees based on VT and EU delineations did not yield high support for the nodes separating the main Glomeromycotina orders; yet, the trees had no significantly-supported conflicts either, and similar branching times of the internal nodes overall (Fig. 2, Supplementary Fig. 4). As expected, finer delineations resulted in an increase in the number of nodes close to the present (Supplementary Fig. 5). However, we observed a slowdown in the accumulation of new lineages close to the present in all lineage through time plots (LTTs), including those with the finest delineations (EU98.5 and EU99; Supplementary Fig. 6).

**Figure 2:**
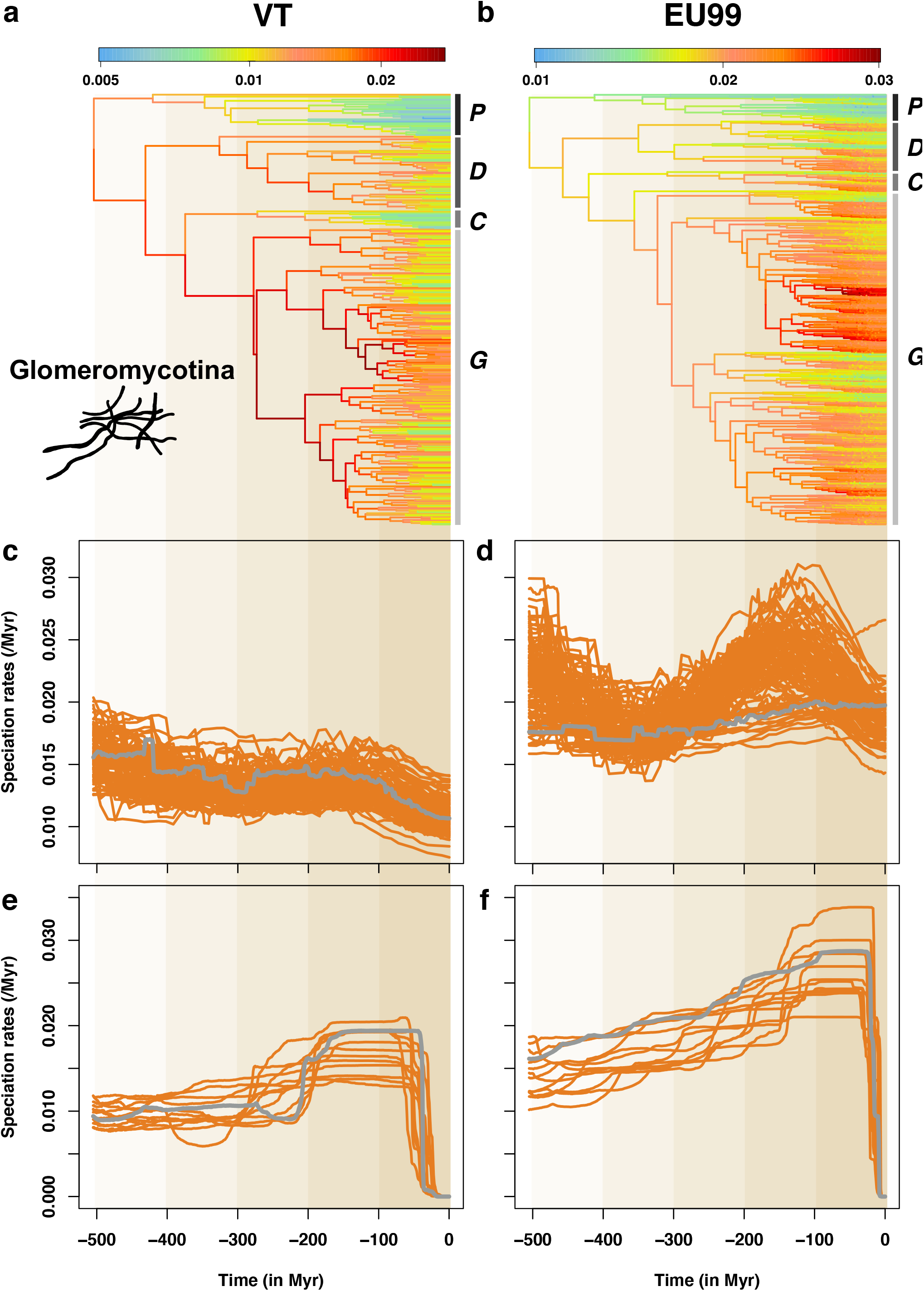
The speciation dynamic of Glomeromycotina (arbuscular mycorrhizal fungi) varies significantly through time and between lineages. **(a-b):** Glomeromycotina consensus phylogenetic trees corresponding to the VT (a) and EU99 (b) species delineations. Branches are colored according to the lineage-specific speciation rates estimated by ClaDS using the BDES estimated sampling fraction: lineages with low and high speciation rates are represented in blue and red, respectively. The main Glomeromycotina clades are indicated with the following letters: *P* = Paraglomerales + Archaeosporales, *D* = Diversisporales, *C* = Claroideoglomeraceae, and *G* = Glomeraceae. **(c-d):** Mean speciation rates through time estimated by ClaDS, for the VT (c) and EU99 (d) delineations and using the BDES estimated sampling fraction. The mean speciation rate corresponds to the maximum *a posteriori* (MAP) of the mean speciation rate across all fungal lineages back in time (including extinct and unsampled lineages). Orange and grey lines represent the independent replicate trees and the consensus tree, respectively: because some of the 12 replicate trees showed different trends, we replicated ClaDS inferences using 100 replicate trees. Unlike most replicate trees, the EU99 consensus tree tends to present a limited decline in speciation rates, which reinforces the idea that consensus trees can be misleading (Janzen & Etienne, 2017). **(e-f):** Mean speciation rates through time estimated by CoMET, for the VT (c) and EU99 (d) delineations and using the BDES estimated sampling fraction. Orange and grey lines represent the 12 independent replicate trees and the consensus tree, respectively.

### Temporal diversification dynamics

We found that speciation rates for Glomeromycotina ranged from 0.005 to 0.03 events per lineage per Myr, using both the VT and EU SSU rRNA delineations (Fig. 2; Supplementary Fig. 7). Speciation rates varied both within and among Glomeromycotina orders, with Glomerales and Diversisporales having the highest present-day speciation rates (Supplementary Fig. 8). As expected we observed higher present-day speciation rates for finer delineations, but at the haplotype level (i.e. at the level of the individual SSU rRNA sequences within each unit) we found a significant correlation of present-day speciation rates computed with ClaDS using different delineations (Supplementary Fig. 9). Whatever the delineations, Glomeromycotina experienced their highest speciation rates between 200 and 100 Myr ago according to estimates obtained with ClaDS (Fig. 2; Supplementary Fig. 10) and between 150 and 50 Myr ago according to CoMET (Fig. 2; Supplementary Fig. 11). ClaDS estimates of speciation rates at 150 Myr ago were 26% (± s.d. 17) higher than those at 300 Myr with the EU99 delineation. With the VT delineation, the increase was of 3% (±8). The peak was even stronger using CoMET: 30% ± 20 higher at 150 Myr in comparison to 300 Myr with the EU99 delineation (71% ± 40 with the VT delineation; Fig. 2).

The peak of speciation rates was followed by a decline in the recent past (Fig. 2; Supplementary Fig. 10), as suggested by the plateauing of the LTTs. A global decline of the speciation rates through time was independently supported by ClaDS and CoMET analyses, as well as time-dependent models in RPANDA (Morlon, Parsons, & Plotkin, 2011)(Supplementary Figs. 11, 12, & 13). This speciation rate decline was robust to all species delineations, the branching process prior (Supplementary Table 7), phylogenetic uncertainty, and assumed sampling fractions as low as 50%, except in ClaDS analyses where the trend disappeared in some EU99 trees and for sampling fractions lower than 70% (Supplementary Figs. 14 & 15). We also found a period of high speciation rates between 200 and 100 Myr ago followed by a decline in our analyses with the LSU region, for assumed sampling fractions as low as 60% (Supplementary Figure 16).

We did not find a strong signal of extinction in our analyses: the turnover rate estimated from ClaDS was generally close to zero (Supplementary Fig. 12b), and models including extinctions were never selected in RPANDA (Supplementary Fig. 13). Similarly, the extinction rates estimated in piecewise-constant models (CoMET) were not significantly different from 0 and we did not find significant support for mass extinction events (Supplementary Fig. 17). Yet, forcing the extinction rate to high positive values did not modify the general trend of speciation rate slowdown (Supplementary Figs. 18 & 19).

### Correlates of Glomeromycotina diversification

When fitting environment-dependent models of diversification, we found that temperature-dependent models better fit Glomeromycotina diversification than time-dependent models, with higher speciation rates during warm climatic periods (Fig. 3; Supplementary Fig. 20). This was true for all Glomeromycotina delineations, sampling fractions, and crown ages (Supplementary Figs. 21, 22, 23, & 24), with the exception of some EU99 trees with a 50% sampling fraction (Supplementary Fig. 24). It was also true in our analyses using the LSU region, for sampling fractions down to 50% (Supplementary Figure 29). This signal of temperature dependency was not due to a temporal trend (Supplementary Figs. 25 & 26) nor to an artefact caused by rate heterogeneities (Supplementary Fig. 27). Evidence for temperature dependency, however, decreased in some clades closer to the present, as small trees tend to be best fit by constant or time-depend models (Supplementary Fig. 28). We detected a significant positive dependency of the speciation rates on CO^2^ concentrations in some sub-trees, but rarely found a significant effect of plant fossil diversity (Supplementary Fig. 28).

**Figure 3:**
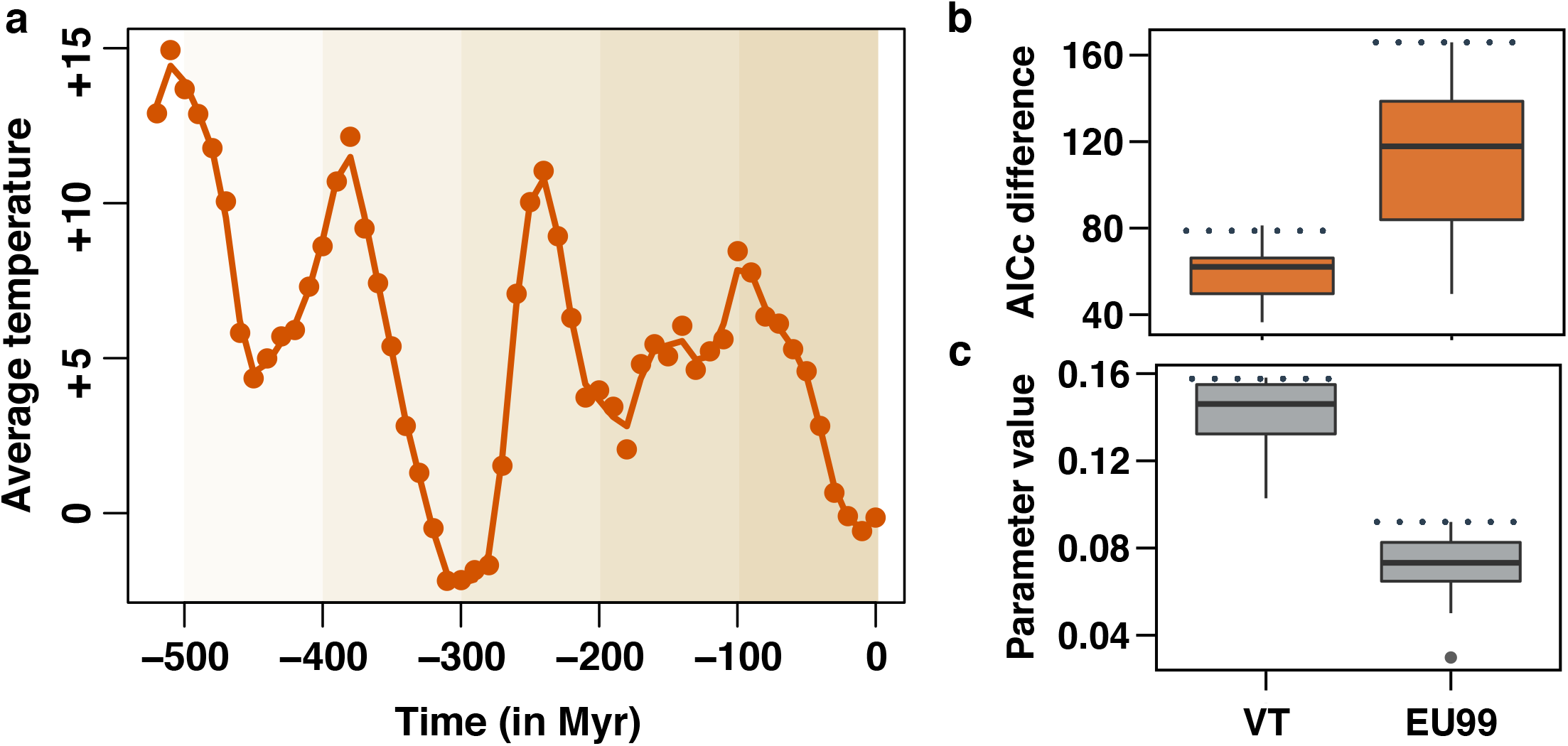
Temperature-dependent diversification models reveal that global temperature positively associates with the speciation rates of Glomeromycotina (arbuscular mycorrhizal fungi) in the last 500 million years. **(a):** Average global temperature in the last 500 million years (Myr) relative to the average temperature of the period 1960-1990. The smoothed orange line represents cubic splines with 33 degrees of freedom used to fit temperature-dependent models of Glomeromycotina diversification with RPANDA. This default smoothing was estimated using the R function *smooth.spline*. **(b):** AICc difference between the best-supported time-dependent model and the temperature-dependent model in RPANDA, for the VT (left) and EU99 (right) delineations, using the BDES estimated sampling fraction. An AICc difference greater than 2 indicates that there is significant support for the temperature-dependent model. **(c):** Parameter estimations of the temperature-dependent models (speciation rate ∼ exp(parameter * temperature)). A positive parameter value indicates a positive effect of temperature on speciation rates. For both delineations, the boxplots represent the results obtained for the consensus tree and the 12 independent replicate trees. Boxplots indicate the median surrounded by the first and third quartiles, and whiskers extend to the extreme values but no further than 1.5 of the inter-quartile range. The horizontal dotted lines highlighted the values estimated for the consensus trees. Compared to the replicate trees, the consensus trees tends to present extreme values (stronger support for temperature-dependent model), which reinforces the idea that consensus trees can be a misleading representation (Janzen & Etienne, 2017).

The PCA of Glomeromycotina relative niche width characteristics had a first principal component (PC1) that indicated the propensity of each Glomeromycotina unit (VT or EUs) to be vastly distributed among continents, ecosystems and/or associated with many plant species and lineages (*i.e.* high generalism), whereas the second principal component (PC2) indicated the propensity of a given Glomeromycotina unit to associate with few plant species on many continents (*i.e.* high specialism toward plants; Supplementary Figs. 30, 31, & 32). Hence, PC1 reflects Glomeromycotina niche width, whereas PC2 discriminates the width of the abiotic relatively to the biotic niche (Fig. 4a-b). We found a positive correlation between lineage-specific speciation rates and PC1 in the majority of the VT and EU99 trees, but no significant correlation with PC2 (Fig. 4c-d; Supplementary Fig. 33a). However, these results were no longer significant when controlling for phylogenetic non-independence between Glomeromycotina units (Supplementary Fig. 33b), likely because a single *Glomeraceae* clade, including the abundant and widespread morphospecies *Rhizophagus irregularis* and *R. clarus* (high PC1 values), had both the highest speciation rates and the largest niche widths among Glomeromycotina (Supplementary Fig. 34).

**Figure 4:**
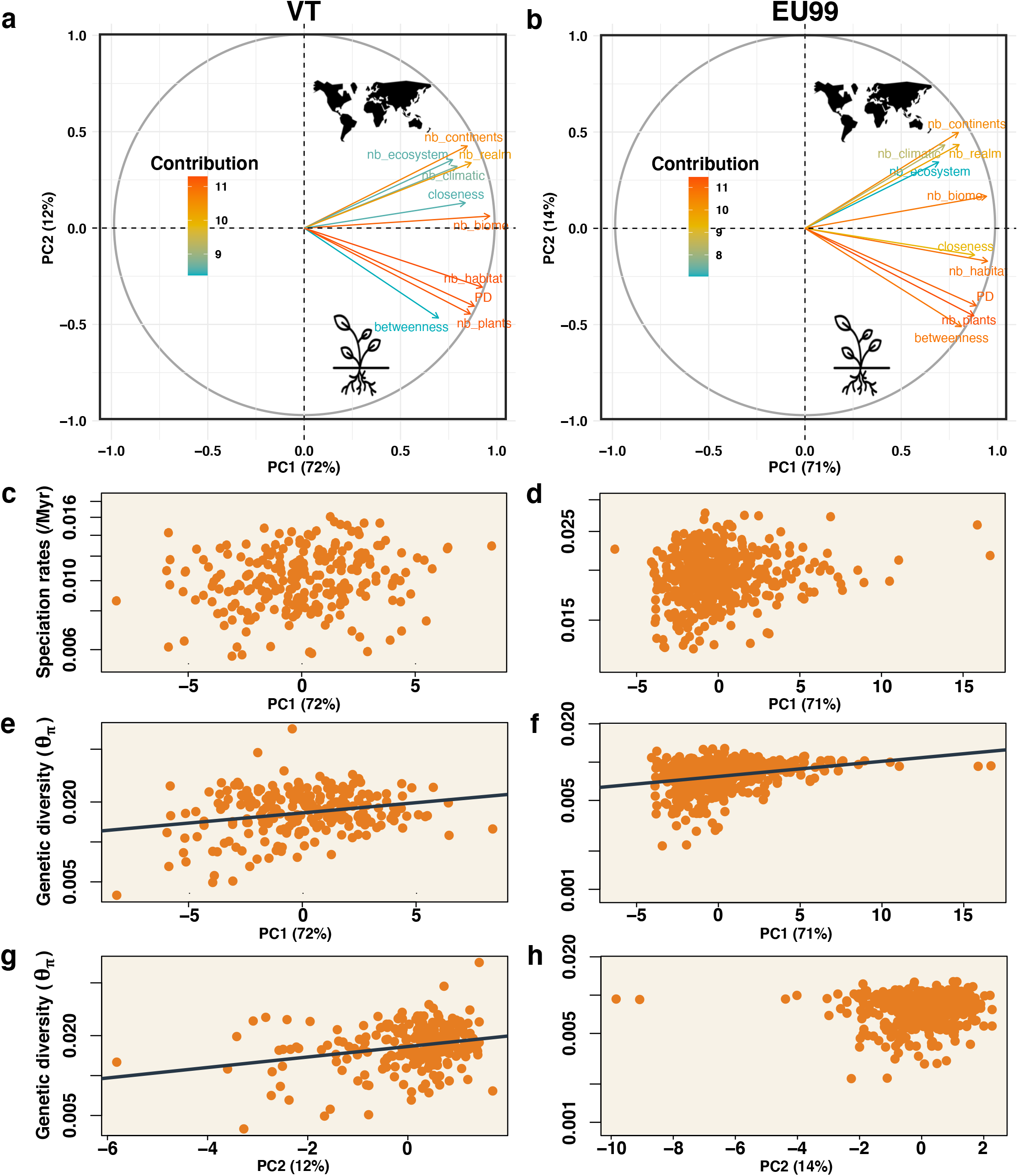
Abiotic and biotic correlates of speciation rates and genetic differentiation in Glomeromycotina (arbuscular mycorrhizal fungi) **(a-b):** Projection of 10 abiotic and biotic variables on the two principal coordinates according to the VT (a) or EU99 (b) delineations. Principal coordinate analysis (PCA) was performed for the Glomeromycotina units represented by at least 10 sequences. Colors represent the contribution of the variable to the principal coordinates. The percentage for each principal coordinate (PC) indicates its amount of explained variance. Tested variables were: the numbers of continents on which the Glomeromycotina unit occurs (nb_continent), of realms (nb_realm), of ecosystems (nb_ecosystems), of habitats (nb_habitats), of biomes (nb_biomes), and climatic zones (nb_climatic) (Öpik et al., 2010), as well as information about the associated plant species of each unit, such as the number of plant partners (nb_plants), the phylogenetic diversity of these plants (PD), and the betweenness and closeness measurement of each fungal unit in the plant-fungus interaction network (see Methods). **(c-d):** Speciation rates as a function of the PC1 coordinates for each VT (c) or EU99 (d) unit. Only the Glomeromycotina consensus tree is represented here (other replicate trees are presented in Supplementary Fig. 33). **(e-h):** Genetic diversity (Tajima’s θπ estimator) as a function of the PC1 (e-f) or PC2 (g-h) coordinates for each VT (e-g) or EU99 (f-h) unit. Only the Glomeromycotina consensus tree is represented here (other replicate trees are presented in Supplementary Fig. 33). The grey lines indicate the statistically significant linear regression between the two variables inferred using MCMCglmm.

Although Glomeromycotina diversity is currently higher in the (sub)tropics (Supplementary Fig. 35), we found no effect of latitude on speciation rates, regardless of the Glomeromycotina delineation or the minimum number of sequences per Glomeromycotina unit (MCMCglmm: *P>*0.05). In addition, we actually did not detect a higher total number of Glomeromycotina species in grasslands compared to forests (Supplementary Figure 36; confirming the results of Davison et al. 2015), and it is thus not surprising that we reported no effect of habitat or climatic zone on speciation rates (Supplementary Fig. 37), suggesting that tropical grasslands are not particular diversification hotspots for Glomeromycotina. Similarly, we recovered no significant correlation between spore size and speciation rate (Supplementary Fig. 38), nor between spore size and level of endemism (Supplementary Fig. 39).

Finally, Tajima’s estimator of Glomeromycotina genetic diversity was significantly and positively correlated with niche width (PC1) for all Glomeromycotina delineations and minimal number of sequences per Glomeromycotina unit considered, and in particular with abiotic aspects of the niche (PC2) in many cases (Fig. 4e-h; Supplementary Fig. 33). Genetic diversity was not correlated with speciation rate (Supplementary Fig. 33), latitude, habitat, climatic zone (MCMCglmm: *P*>0.05), or spore size (PGLS: *P*>0.05).

### Simulation results

When we simulated the evolution of a short DNA gene mimicking the SSU rRNA marker and used it to delineate species, we found that the number of EU99 delineated units was generally lower than the number of simulated species (∼10% to 20% lower; Supplementary Figure 2b). Hence, even the EU99 delineation tends to lump together some closely related species. As expected, this lumping resulted in an artefactual inference of a decline of speciation rates toward the present, but this artifactual decline was significantly smaller in magnitude than that observed in Glomeromycotina (Figure 5). Hence, these analyses suggest that the lumping of species resulting from the use of a small, slowly evolving marker is unlikely to fully explain the strong temporal decline in speciation rate we found in Glomeromycotina.

**Figure 5:**
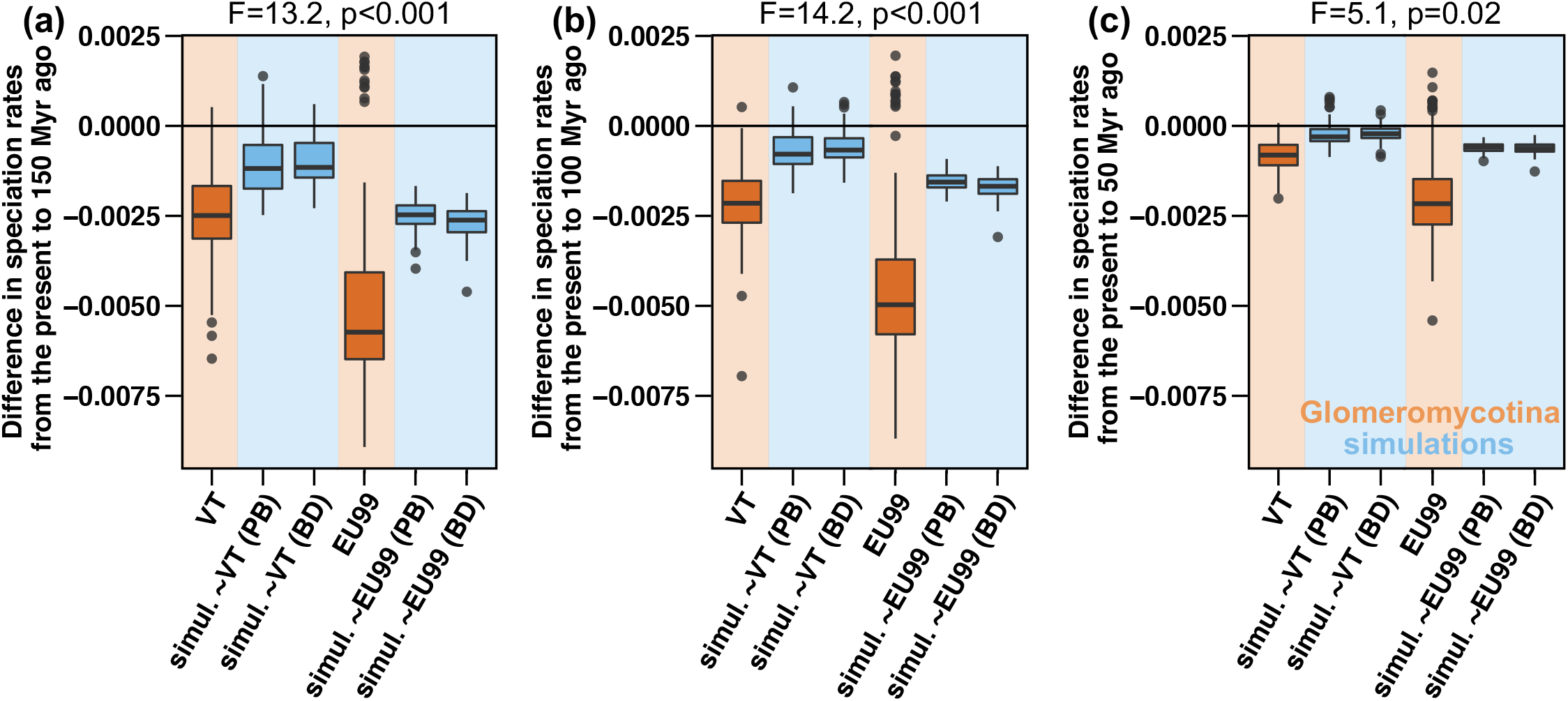
Artefactual species lumping and lack of phylogenetic resolution in the SSU rRNA region are not enough to explain the temporal decline in speciation rates detected in Glomeromycotina. Comparison of the magnitude of the decline in speciation rates observed in Glomeromycotina (in orange) and on simulated data (in blue). The intensity of the slowdown is measured as the difference between the mean speciation rate estimated at present and the mean speciation rate estimated 150 Myr ago (a), 100 Myr ago (b), or 50 Myr ago (c). Negative differences indicate a speciation rate decline. Sequence alignments were simulated on phylogenetic trees obtained under a scenario of constant speciation rate and no extinction (*i.e.* pure birth “PB”) and constant speciation and extinction rates (*i.e.* birth death “BD”), with characteristics mimicking the slow evolution of the SSU rRNA marker. We simulated phylogenies with two different net diversification rates, such that we obtained simulations with total numbers of species similar to the total numbers of VT or EU99 units (Supplementary Fig. 2). Boxplots indicate the median surrounded by the first and third quartiles, and whiskers extend to the extreme values but no further than 1.5 of the inter-quartile range. Each boxplot represents results for the consensus trees and the 12 independent replicate trees for each of the 10 simulations, and for the consensus trees and the 100 independent replicate trees for the Glomeromycotina. Differences between the magnitude of the decline measured in Glomeromycotina (VT or EU99) and in the corresponding simulations were tested using linear models (reported at the top of the plots).

## Discussion

### Glomeromycotina species delineations, diversity, and phylogeny

It is difficult to delineate species in Glomeromycotina, which are poorly differentiated morphologically and mainly characterized by environmental sequences (Bruns et al., 2018). Our GMYC analyses suggest that Glomeromycotina species-like units correspond to SSU rRNA haplotypes with a sequence similarity between 98.5 and 99%. With this criterion of species delineation, we estimate that there are between 1,300 and 2,900 Glomeromycotina ‘species’. These estimates are largely above the number of currently described morphospecies or VT (Supplementary Note 1) but remain low in comparison with other fungal groups, like the Agaricomycetes that include taxa forming ectomycorrhiza (Varga et al., 2019).

Our phylogenies based on the SSU rRNA gene did not resolve the branching of the Glomeromycotina orders, with node supports similar to those of previous studies (Davison et al., 2015; Krüger et al., 2012; Rimington et al., 2018)(Supplementary Note 2). These findings confirm that additional genomic evidence is required to reach consensus. We considered this uncertainty in species delineation and phylogenetic reconstruction by repeating our diversification analyses across species delineation criteria and on a set of trees spanning the likely tree space. We found effects of species delineation consistent with *a priori* expectations: criteria that lump together more dissimilar sequences (e.g. those that use a lower percentage of similarity cut-off) result in lower diversity estimates, lower estimates of speciation rates, and patterns of diversification through time that reflect longer terminal branch-lengths, such as peaks of diversification that occur earlier. Despite this variability, we found that general patterns, such as the observed temporal decline in speciation rates and the significant association between temperature and speciation rates, were consistent across species delineations and trees. Therefore, our study based on a short SSU (or LSU) rRNA region should encourage both efforts to obtain more genetic data, including longer reads (Krehenwinkel et al., 2019; Tedersoo, Albertsen, Anslan, & Callahan, 2021) and additional genomic information, with the aim of reconstructing better supported, comprehensive phylogenies and efforts to conduct diversification analyses despite uncertainty in the data for groups where better data is not yet available.

### Glomeromycotina diversify slowly

We found speciation rates for Glomeromycotina an order of magnitude lower than rates typically found for macro-eukaryotes (Maliet et al., 2019; Upham et al., 2019), like plants (Zanne et al., 2014), or Agaricomycetes (Varga et al., 2019). Low speciation rates in Glomeromycotina may be linked to their multinucleate hyphal state (Yildirir, Malar, Kokkoris, & Corradi, 2020), to their occasional long-distance dispersal that homogenizes populations globally over evolutionary timescales (Savary et al., 2018), and/or to the fact that they are generalist obligate symbionts (Morlon, Kemps, Plotkin, & Brisson, 2012). Regardless of the proximal cause, and contrary to Agaricomycetes for example, which present a large diversity of species, morphologies, and ecologies, Glomeromycotina have poorly diversified in the last 500 Myr despite their ubiquity; their niche space is restricted to plant roots and the surrounding soil because of their obligate dependence on plants for more than 400 Myr (Rich, Nouri, Courty, & Reinhardt, 2017; Tisserant et al., 2013).

Our estimates of speciation rates were highly variable across lineages. We reported the highest speciation rates in Glomeraceae and Diversisporaceae. Speciation rates in Paraglomeraceae and Archaeosporaceae, which are thought to be less beneficial for the plants than the fast diversifying Glomeraceae and Diversisporaceae (Säle et al., 2021), were an order of magnitude lower. We can therefore speculate that good symbiotic abilities may favor diversification, although this remains to be tested in further investigations.

We found little evidence for species extinction in Glomeromycotina, including at mass extinction events. Because Glomeromycotina are relatively widespread and have a an ancient tendency toward generalism, they might therefore be quite resilient to land plant mass extinctions and low extinction rates have been predicted before based on their ecology (Morton, 1990). Yet, these low extinction rate estimates could also come from the difficulty of estimating extinction from molecular phylogenies (Rabosky, 2016), one of the limitations of phylogeny-based diversification analyses (Supplementary Note 3). Fossils of Glomeromycotina that can be ascribed to species or genera are too scarce to support or conflict with this finding.

### Glomeromycotina diversification through time

The observed peak of Glomeromycotina speciations detected between 200 and 100 Myr (or 150-50 Myr depending on the models) was mainly linked to the frequent speciations in the largest family Glomeraceae. This peak was concomitant with the radiation of flowering plants (Sauquet & Magallón, 2018), but also with a major continental reconfiguration, including the breakdown of Pangea and the formation of climatically contrasted landmasses (Davison et al., 2015). This period was also characterized by a warm climate potentially directly or indirectly favorable to Glomeromycotina diversification, such that disentangling the impact of these various factors on Glomeromycotina diversification rates is not straightforward. Interestingly, a peak of speciations at this period was also found in the Agaricomycetes, a clade of fungi including lineages forming ectomycorrhizae (Varga et al., 2019).

This peak in the occurrence of speciation events was followed by a decline in speciation rates. The detection of temporal declines in speciation rates in phylogenetic diversification analyses can sometimes be artifactual, for example if some species are incorrectly lumped together during species delineation or if the proportion of species not represented in the phylogeny is under-estimated (Moen & Morlon, 2014). We considered these potential biases, conducted sensitivity analyses, and found that the observed slowdown was robust, and even amplified under scenarios of high extinction. Some Glomeromycotina species are likely lumped together into the same SSU haplotypes (Krüger et al., 2012), but both our use of an overly small assumed sampling intensity (50%) and our simulation analyses demonstrated that this lumping is not sufficient to explain the observed slowdown. In addition, we also detected a temporal decline in speciation rates when using another marker (the LSU rRNA gene).

Temporal declines in speciation rates have been observed in many clades, including microorganisms (Condamine, Rolland, & Morlon, 2019; Morlon et al., 2012; Rabosky & Lovette, 2008). They have often been interpreted as a progressive reduction of the number of available niches as species diversify and accumulate (Moen & Morlon, 2014; Rabosky, 2009). In Glomeromycotina, this potential effect of niche saturation could be exacerbated by a reduction of their niches linked to both repetitive breakdowns of their symbiosis with plants and climatic changes. Indeed, since the Cretaceous, many plant lineages evolved alternative root symbioses or became non-symbiotic (Brundrett & Tedersoo, 2018; Maherali, Oberle, Stevens, Cornwell, & McGlinn, 2016; Selosse & Le Tacon, 1998; Werner et al., 2018): approximately 20% of extant plants do not interact with Glomeromycotina anymore (van der Heijden et al., 2015). Additionally, the cooling of the Earth during the Cenozoic reduced the surface of tropical regions (Meseguer & Condamine, 2020; Ziegler et al., 2003), which tend to be a reservoir of ecological niches for Glomeromycotina (Brundrett & Tedersoo, 2018; Davison et al., 2015; Read, 1991).

The difficulty of reconstructing past symbiotic associations prevents direct testing the hypothesis that the emergence of new root symbioses in plants led to a decline in speciation rates in Glomeromycotina. However, we tested the hypothesis that global temperature changes affected speciation rates and found a strong relationship. Such associations between temperature and speciation rates have been observed before in eukaryotes and have several potential causes (Condamine et al., 2019). In particular, the productivity hypothesis states that resources and associated ecological niches are more numerous in warm and productive environments, especially when the tropics are large, which entail higher speciation rates (Clarke & Gaston, 2006). This hypothesis is particularly relevant for Glomeromycotina, which have many host plant niches in the tropics, as shown by their latitudinal diversity gradient, and potentially relatively less in temperate and polar regions (Toussaint et al., 2020), where a higher proportion of plants are non-mycorrhizal (Bueno et al., 2017) or ectomycorrhizal (Brundrett & Tedersoo, 2018; Varga et al., 2019). Hence, the observed effect of past global temperatures could reflect the shrinkage of tropical areas and the associated decrease of the relative proportion of arbuscular mycorrhizal plants. Future developments of diversification models incorporating interspecific interactions would allow us to better test these hypotheses.

A few Glomeromycotina clades displayed a significant support for diversification models with a positive dependency of speciation rates on CO^2^ concentrations, which reinforces the idea that for the corresponding Glomeromycotina, benefits retrieved from plants could have been amplified by high CO^2^ concentrations and fostered diversification (Field et al., 2016; Humphreys et al., 2010). Conversely, we found a limited effect of land plant fossil diversity, which indicates that variations in the tempo of Glomeromycotina diversification did not systematically follow those of land plants. Still, the possible concordance of the peak of Glomeromycotina speciations with the radiation of the Angiosperms is noteworthy, in particular in Glomeraceae that frequently interact with present-day Angiosperms (Rimington et al., 2018). Plant diversification might have fostered the diversification of Glomeromycotina from the emergence of land plants until the Mesozoic (Lutzoni et al., 2018; Morton, 1990), but less so thereafter, when Glomeromycotina diversification declined while some flowering plants radiated, including Glomeromycotina-associated groups, like the Poaceae, but also Glomeromycotina-free groups such as the extraordinary radiation of Orchidaceae (Givnish et al., 2015), blurring co-diversification patterns (Supplementary Fig. 40)(Cleal & Cascales-Miñana, 2014; Ramírez-Barahona, Sauquet, & Magallón, 2020).

### Correlates of Glomeromycotina recent speciation rates

Looking at the correlates of Glomeromycotina present-day speciation rates, we found no effect of habitat or climatic zone, even though Glomeromycotina are more frequent and diverse in the tropics (Davison et al., 2015; Pärtel et al., 2017; Toussaint et al., 2020) and a positive correlation with global temperature. Further work, including a more thorough sampling of the distribution of Glomeromycotina species across latitudes and habitats, would be required to confirm these patterns and to distinguish whether speciation events are indeed no more frequent in the tropics or, if they are, whether long-distance dispersal redistributes the new lineages at different latitudes over long time scales. Contrary to previous predictions (Pärtel et al., 2017), we did not find that tropical grasslands are diversification hotspots for Glomeromycotina; we actually did not even find higher Glomeromycotina species richness in (tropical) grasslands *versus* forests at global scale, in agreement with Davison *et al*. (2015).

Similarly, although the temporal changes in the availability of Glomeromycotina niches likely influenced the diversification of the group, we found little support for Glomeromycotina species with larger niche width having higher lineage-specific speciation rates. We also note that there are important aspects of the niche that we do not (and yet cannot) account for in our characterization of Glomeromycotina niche width: it is thought that some Glomeromycotina species may mainly provide mineral nutrients extracted from the soil, whereas others may be more specialized in protecting plants from biotic or abiotic stresses (Chagnon, Bradley, Maherali, & Klironomos, 2013) and such (inter- or intra-specific) functional variations may have evolutionary significance. Finally, although spore size is often inversely related to dispersal capacity (Nathan et al., 2008), which can limit speciation by increasing gene flow, we found no significant correlation between spore size and speciation rates, which may be explained either by a weak or absent effect or by the low number of species for which this data is available. In addition, the absence of correlation between spore size and level of endemism suggests that even Glomeromycotina with large spores experience long-distance dispersal (Davison et al., 2018; Kivlin, 2020). Thus, if large spores might limit dispersal at smaller (*e.g.* intra-continental) scales in Glomeromycotina (Bueno & Moora, 2019; Chaudhary, Nolimal, Sosa-Hernández, Egan, & Kastens, 2020), this does not seem to affect speciation rates.

In Glomeromycotina, intraspecific variability is an important source of functional diversity (Munkvold, Kjøller, Vestberg, Rosendahl, & Jakobsen, 2004; Savary et al., 2018) and their genetic diversity may indicate the intraspecific variability on which selection can act, potentially leading to speciation. Here, geographically widespread Glomeromycotina species appear to be more genetically diverse, as previously suggested by population genomics (Savary et al., 2018), but do not necessarily speciate more frequently. Along with a decoupling between genetic diversity and lineage-specific speciation rate, this suggests that the accumulation of genetic diversity in the SSU region among distant subpopulations is not enough to spur Glomeromycotina speciation.

### Analyzing diversification dynamics using a short marker gene

Short DNA regions, like those used in metabarcoding surveys, typically do not allow to robustly delineate species, estimate global-scale diversity, and reconstruct phylogenetic trees. As these three aspects can all affect results of diversification analyses (Moen & Morlon, 2014), such analyses are rarely performed with these types of data. Yet, for many species-rich groups of organisms, in particular microorganisms, no other data currently provide a thorough representation of diversity at the “species” level. Hence, these data, although far from ideal, are the only one that can be used to study the past diversification of such groups (see Lewitus et al., 2018; Louca et al., 2018 for antecedents). The approach we took here is to recognize all these potential sources of uncertainty and biases and to test the robustness of our results. We demonstrated the usefulness of this approach: while some results inevitably depend on the choices made for species delineation, phylogenetic reconstruction, and the estimation of global scale diversity, others are sufficiently strong to hold despite uncertainty in the data. Our results therefore illustrate that using a short DNA marker (e.g. a metabarcode) combined with intensive sensitivity analyses can be useful for studying the diversification dynamics of poorly-known groups.

## Conclusion

Our findings that Glomeromycotina have low speciation rates, likely constrained by the availability of suitable niches, reinforce the vision of Glomeromycotina as an “evolutionary cul-de-sac” (Malloch, 1987). We interpret the significant decline in speciation rates toward the present as the conjunction of the emergence of plant lineages not associated with Glomeromycotina and the reduction of tropical areas induced by climate cooling, in the context of obligate dependence of Glomeromycotina on plants. Temporal declines in speciation rates have often been interpreted as the signal of adaptive radiations (Harmon, Schulte, Larson, & Losos, 2003; Moen & Morlon, 2014), that is clades that experienced a rapid accumulation of morphological, ecological, and species diversity (Simpson, 1953). Conversely, Glomeromycotina provide here a striking example of a clade with slow morphological, ecological, and species diversification that features a pattern of temporal decline in speciation rates, that might reflect the reduction of the global availability of their mycorrhizal niches.

## Acknowledgment

The authors acknowledge C. Strullu-Derrien, M. Elias, D. de Vienne, A. Vogler, J.-Y. Dubuisson, C. Quince, S.-K. Sepp, and M. Chase for helpful discussions. They also thank L. Aristide, S. Lambert, J. Clavel, I. Quintero, I. Overcast, and G. Sommeria for comments on the manuscript, D. Marsh for English editing, and the Editor and three reviewers for improvements of an earlier version of this manuscript. BPL acknowledges B. Robira, F. Foutel-Rodier, F. Duchenne, E. Faure, E. Kerdoncuff, R. Petrolli, and G. Collobert for useful discussions and C. Fruciano and E. Lewitus for providing codes. This work was supported by a doctoral fellowship from the École Normale Supérieure de Paris attributed to BPL and the École Doctorale FIRE – Programme Bettencourt. MÖ was supported by the European Regional Development Fund (Centre of Excellence EcolChange) and University of Tartu (PLTOM20903). Funding of the research of FM was from the Agence Nationale de la Recherche (ANR-19-CE02-0002). HM acknowledges support from the European Research Council (grant CoG-PANDA).

## Data Accessibility

All of the data used in this study are available in the open-access MaarjAM database (https://maarjam.botany.ut.ee). Spore lengths of Glomeromycotina were collected in the supplementary data of Davison et al. (2018). New scripts for delineating evolutionary units (EU) are available in the R-package RPANDA (through this GiHub branch for the moment: https://github.com/hmorlon/PANDA/tree/Benoit_phylosignal). The alignment of all the SSU rRNA sequences and their associated metadata publicly accessible through the Open Science Framework (osf) portal: osf.io/y2ts5.

## Author contributions

All the authors designed the study. MÖ gathered the data and BPL performed the analyses. OM and ACAS provided some codes. BPL and HM wrote the first version of the manuscript and all authors contributed substantially to the revisions.

## References

Alroy, J. (2010). Geographical, environmental and intrinsic biotic controls on Phanerozoic marine diversification. Palaeontology, 53(6), 1211–1235. doi:10.1111/j.1475-4983.2010.01011.x

Barnosky, A. D. (2001). Distinguishing the effects of the red queen and court jester on miocene mammal evolution in the northern rocky mountains. Journal of Vertebrate Paleontology, 21(1), 172–185. doi:10.1671/0272-4634(2001)021[0172:DTEOTR]2.0.CO;2

Benton, M. J. (2009). The Red Queen and the Court Jester: Species diversity and the role of biotic and abiotic factors through time. Science, 323(5915), 728–732. doi:10.1126/science.1157719

Bouckaert, R., Heled, J., Kühnert, D., Vaughan, T., Wu, C.-H., Xie, D., … Drummond, A. J. (2014). BEAST 2: A software platform for Bayesian evolutionary analysis. PLoS Computational Biology, 10(4), e1003537. doi:10.1371/journal.pcbi.1003537

Bredenkamp, G. J., Spada, F., & Kazmierczak, E. (2002). On the origin of northern and southern hemisphere grasslands. Plant Ecology, 163(2), 209–229. doi:10.1023/A:1020957807971

Brundrett, M. C., & Tedersoo, L. (2018). Evolutionary history of mycorrhizal symbioses and global host plant diversity. New Phytologist, 220(4), 1108–1115. doi:10.1111/nph.14976

Bruns, T. D., Corradi, N., Redecker, D., Taylor, J. W., & Öpik, M. (2018). Glomeromycotina: what is a species and why should we care? New Phytologist, 220(4), 963–967. doi:10.1111/nph.14913

Bueno, C. G., & Moora, M. (2019). How do arbuscular mycorrhizal fungi travel? New Phytologist, 222(2), 645–647. doi:10.1111/nph.15722

Bueno, C. G., Moora, M., Gerz, M., Davison, J., Öpik, M., Pärtel, M., … Zobel, M. (2017). Plant mycorrhizal status, but not type, shifts with latitude and elevation in Europe. Global Ecology and Biogeography, 26(6), 690–699. doi:10.1111/geb.12582

Chagnon, P.-L., Bradley, R. L., Maherali, H., & Klironomos, J. N. (2013). A trait-based framework to understand life history of mycorrhizal fungi. Trends in Plant Science, 18(9), 484–491. doi:10.1016/j.tplants.2013.05.001

Chaudhary, V. B., Nolimal, S., Sosa-Hernández, M. A., Egan, C., & Kastens, J. (2020). Trait-based aerial dispersal of arbuscular mycorrhizal fungi. New Phytologist, 228(1), 238–252. doi:10.1111/nph.16667

Chomicki, G., Kiers, E. T., & Renner, S. S. (2020). The evolution of mutualistic dependence. Annual Review of Ecology, Evolution, and Systematics, 51(1), 409–432. doi:10.1146/annurev-ecolsys-110218-024629

Clarke, A., & Gaston, K. J. (2006). Climate, energy and diversity. Proceedings of the Royal Society B: Biological Sciences, 273(1599), 2257–2266. doi:10.1098/rspb.2006.3545

Cleal, C. J., & Cascales-Miñana, B. (2014). Composition and dynamics of the great Phanerozoic Evolutionary Floras. Lethaia, 47(4), 469–484. doi:10.1111/let.12070

Condamine, F. L., Rolland, J., Höhna, S., Sperling, F. A. H., & Sanmartín, I. (2018). Testing the role of the Red Queen and Court Jester as drivers of the macroevolution of Apollo butterflies. Systematic Biology, 67(6), 940–964. doi:10.1093/sysbio/syy009

Condamine, F. L., Rolland, J., & Morlon, H. (2013). Macroevolutionary perspectives to environmental change. Ecology Letters, 16(SUPPL.1), 72–85. doi:10.1111/ele.12062

Condamine, F. L., Rolland, J., & Morlon, H. (2019). Assessing the causes of diversification slowdowns: temperature-dependent and diversity-dependent models receive equivalent support. Ecology Letters, 22(11), 1900–1912. doi:10.1111/ele.13382

Correia, M., Heleno, R., da Silva, L. P., Costa, J. M., & Rodríguez-Echeverría, S. (2019). First evidence for the joint dispersal of mycorrhizal fungi and plant diaspores by birds. New Phytologist, 222(2), 1054–1060. doi:10.1111/nph.15571

Davison, J., Moora, M., Öpik, M., Adholeya, A., Ainsaar, L., Bâ, A., … Zobel, M. (2015). Global assessment of arbuscular mycorrhizal fungus diversity reveals very low endemism. Science, 349(6251), 970–973. doi:10.1126/science.aab1161

Davison, J., Moora, M., Öpik, M., Ainsaar, L., Ducousso, M., Hiiesalu, I., … Zobel, M. (2018). Microbial island biogeography: isolation shapes the life history characteristics but not diversity of root-symbiotic fungal communities. The ISME Journal, 12(9), 2211–2224. doi:10.1038/s41396-018-0196-8

Delavaux, C. S., Sturmer, S. L., Wagner, M. R., Schütte, U., Morton, J. B., & Bever, J. D. (2020). Utility of large subunit for environmental sequencing of arbuscular mycorrhizal fungi: a new reference database and pipeline. New Phytologist, 1–5. doi:10.1111/nph.17080

Egan, C., Li, D.-W., & Klironomos, J. (2014). Detection of arbuscular mycorrhizal fungal spores in the air across different biomes and ecoregions. Fungal Ecology, 12, 26–31. doi:10.1016/j.funeco.2014.06.004

Ezard, T., Fujisawa, T., & Barraclough, T. G. (2009). SPLITS: SPecies’ LImits by Threshold Statistics. R-package.

Feijen, F. A., Vos, R. A., Nuytinck, J., & Merckx, V. S. F. T. (2018). Evolutionary dynamics of mycorrhizal symbiosis in land plant diversification. Scientific Reports, 8(1), 10698. doi:10.1038/s41598-018-28920-x

Field, K. J., Pressel, S., Duckett, J. G., Rimington, W. R., & Bidartondo, M. I. (2015). Symbiotic options for the conquest of land. Trends in Ecology & Evolution, 30(8), 477– 486. doi:10.1016/j.tree.2015.05.007

Field, K. J., Rimington, W. R., Bidartondo, M. I., Allinson, K. E., Beerling, D. J., Cameron, D. D., … Pressel, S. (2016). Functional analysis of liverworts in dual symbiosis with Glomeromycota and Mucoromycotina fungi under a simulated Palaeozoic CO2 decline. ISME Journal, 10(6), 1514–1526. doi:10.1038/ismej.2015.204

Foster, G. L., Royer, D. L., & Lunt, D. J. (2017). Future climate forcing potentially without precedent in the last 420 million years. Nature Communications, 8(1), 14845. doi:10.1038/ncomms14845

Fujisawa, T., & Barraclough, T. G. (2013). Delimiting species using single-locus data and the generalized mixed yule coalescent approach: A revised method and evaluation on simulated data sets. Systematic Biology, 62(5), 707–724. doi:10.1093/sysbio/syt033

Gelman, A., & Rubin, D. B. (1992). Inference from iterative simulation using multiple sequences. Statistical Science, 7(4), 457–472. doi:10.1214/ss/1177011136

Givnish, T. J., Spalink, D., Ames, M., Lyon, S. P., Hunter, S. J., Zuluaga, A., … Cameron, K. M. (2015). Orchid phylogenomics and multiple drivers of their extraordinary diversification. Proceedings of the Royal Society B: Biological Sciences, 282(1814), 20151553. doi:10.1098/rspb.2015.1553

Hadfield, J. D. (2010). MCMC methods for multi-response generalized linear mixed models: The MCMCglmm R package. Journal of Statistical Software, 33(2), 1–22. doi:10.18637/jss.v033.i02

Harmon, L. J., Schulte, J. A., Larson, A., & Losos, J. B. (2003). Tempo and mode of evolutionary radiation in iguanian lizards. Science, 301(5635), 961–964. doi:10.1126/science.1084786

Höhna, S., May, M. R., & Moore, B. R. (2016). TESS: An R package for efficiently simulating phylogenetic trees and performing Bayesian inference of lineage diversification rates. Bioinformatics, 32(5), 789–791. doi:10.1093/bioinformatics/btv651

Humphreys, C. P., Franks, P. J., Rees, M., Bidartondo, M. I., Leake, J. R., & Beerling, D. J. (2010). Mutualistic mycorrhiza-like symbiosis in the most ancient group of land plants. Nature Communications, 1(8), 103. doi:10.1038/ncomms1105

James, T. Y., Kauff, F., Schoch, C. L., Matheny, P. B., Hofstetter, V., Cox, C. J., … Vilgalys, R. (2006). Reconstructing the early evolution of Fungi using a six-gene phylogeny. Nature, 443(7113), 818–822. doi:10.1038/nature05110

Janzen, T., & Etienne, R. S. (2017). Inferring the role of habitat dynamics in driving diversification: evidence for a species pump in Lake Tanganyika cichlids. BioRxiv, 11(2), 1–18. doi:https://doi.org/10.1101/085431

Katoh, K., & Standley, D. M. (2013). MAFFT Multiple sequence alignment software version 7: Improvements in performance and usability. Molecular Biology and Evolution, 30(4), 772–780. doi:10.1093/molbev/mst010

Kivlin, S. N. (2020). Global mycorrhizal fungal range sizes vary within and among mycorrhizal guilds but are not correlated with dispersal traits. Journal of Biogeography, 47(9), 1994–2001. doi:10.1111/jbi.13866

Krehenwinkel, H., Pomerantz, A., Henderson, J. B., Kennedy, S. R., Lim, J. Y., Swamy, V., … Prost, S. (2019). Nanopore sequencing of long ribosomal DNA amplicons enables portable and simple biodiversity assessments with high phylogenetic resolution across broad taxonomic scale. GigaScience, 8(5). doi:10.1093/gigascience/giz006

Krüger, M., Krüger, C., Walker, C., Stockinger, H., & Schüßler, A. (2012). Phylogenetic reference data for systematics and phylotaxonomy of arbuscular mycorrhizal fungi from phylum to species level. New Phytologist, 193(4), 970–984. doi:10.1111/j.1469-8137.2011.03962.x

Lee, J., Lee, S., & Young, J. P. W. (2008). Improved PCR primers for the detection and identification of arbuscular mycorrhizal fungi. FEMS Microbiology Ecology, 65(2), 339– 349. doi:10.1111/j.1574-6941.2008.00531.x

Lekberg, Y., Vasar, M., Bullington, L. S., Sepp, S.-K. K., Antunes, P. M., Bunn, R., … Öpik, M. (2018). More bang for the buck? Can arbuscular mycorrhizal fungal communities be characterized adequately alongside other fungi using general fungal primers? New Phytologist, 220(4), 971–976. doi:10.1111/nph.15035

Lewitus, E., Bittner, L., Malviya, S., Bowler, C., & Morlon, H. (2018). Clade-specific diversification dynamics of marine diatoms since the Jurassic. Nature Ecology and Evolution, 2(11), 1715–1723. doi:10.1038/s41559-018-0691-3

Louca, S., Shih, P. M., Pennell, M. W., Fischer, W. W., Parfrey, L. W., & Doebeli, M. (2018). Bacterial diversification through geological time. Nature Ecology and Evolution, 2(9), 1458–1467. doi:10.1038/s41559-018-0625-0

Lutzoni, F., Nowak, M. D., Alfaro, M. E., Reeb, V., Miadlikowska, J., Krug, M., … Magallón, S. (2018). Contemporaneous radiations of fungi and plants linked to symbiosis. Nature Communications, 9(1), 1–11. doi:10.1038/s41467-018-07849-9

Magallón, S., & Sanderson, M. J. (2001). Absolute diversification rates in angiosperm clades. Evolution, 55(9), 1762–1780. doi:10.1111/j.0014-3820.2001.tb00826.x

Maherali, H., Oberle, B., Stevens, P. F., Cornwell, W. K., & McGlinn, D. J. (2016). Mutualism persistence and abandonment during the evolution of the mycorrhizal symbiosis. American Naturalist, 188(5), E113–E125. doi:10.1086/688675

Maliet, O., Hartig, F., & Morlon, H. (2019). A model with many small shifts for estimating species-specific diversification rates. Nature Ecology & Evolution, 3(7), 1086–1092. doi:10.1038/s41559-019-0908-0

Maliet, O., & Morlon, H. (2022). Fast and accurate estimation of species-specific diversification rates using data augmentation. Systematic Biology, 71(2), 353–366. doi:10.1093/sysbio/syab055

Malloch, D. M. (1987). The evolution of mycorrhizae. Can. J. Plant. Path., 9, 398–402.

May, M. R., Höhna, S., & Moore, B. R. (2016). A Bayesian approach for detecting the impact of mass-extinction events on molecular phylogenies when rates of lineage diversification may vary. Methods in Ecology and Evolution, 7(8), 947–959. doi:10.1111/2041-210X.12563

Meseguer, A. S., & Condamine, F. L. (2020). Ancient tropical extinctions at high latitudes contributed to the latitudinal diversity gradient. Evolution, 74(9), 1966–1987. doi:10.1111/evo.13967

Moen, D., & Morlon, H. (2014). Why does diversification slow down? Trends in Ecology and Evolution, 29(4), 190–197. doi:10.1016/j.tree.2014.01.010

Morlon, H. (2014). Phylogenetic approaches for studying diversification. Ecology Letters, 17(4), 508–525. doi:10.1111/ele.12251

Morlon, H., Kemps, B. D., Plotkin, J. B., & Brisson, D. (2012). Explosive radiation of a bacterial species group. Evolution, 66(8), 2577–2586. doi:10.1111/j.1558-5646.2012.01598.x

Morlon, H., Lewitus, E., Condamine, F. L., Manceau, M., Clavel, J., & Drury, J. (2016). RPANDA: An R package for macroevolutionary analyses on phylogenetic trees. Methods in Ecology and Evolution, 7(5), 589–597. doi:10.1111/2041-210X.12526

Morlon, H., Parsons, T. L., & Plotkin, J. B. (2011). Reconciling molecular phylogenies with the fossil record. Proceedings of the National Academy of Sciences, 108(39), 16327–16332. doi:10.1073/pnas.1102543108

Morton, J. B. (1990). Species and clones of arbuscular mycorrhizal fungi (Glomales, Zygomycetes): their role in macro- and microevolutionary processes. Mycotaxon (USA), 37, 493–515.

Munkvold, L., Kjøller, R., Vestberg, M., Rosendahl, S., & Jakobsen, I. (2004). High functional diversity within species of arbuscular mycorrhizal fungi. New Phytologist, 164(2), 357–364. doi:10.1111/j.1469-8137.2004.01169.x

Nathan, R., Schurr, F. M., Spiegel, O., Steinitz, O., Trakhtenbrot, A., & Tsoar, A. (2008). Mechanisms of long-distance seed dispersal. Trends in Ecology & Evolution, 23(11), 638–647. doi:10.1016/j.tree.2008.08.003

Öpik, M., Davison, J., Moora, M., & Zobel, M. (2014). DNA-based detection and identification of Glomeromycota: the virtual taxonomy of environmental sequences. Botany, 92(2), 135–147. doi:10.1139/cjb-2013-0110

Öpik, M., Vanatoa, A., Vanatoa, E., Moora, M., Davison, J., Kalwij, J. M., … Zobel, M. (2010). The online database MaarjAM reveals global and ecosystemic distribution patterns in arbuscular mycorrhizal fungi (Glomeromycota). New Phytologist, 188(1), 223–241. doi:10.1111/j.1469-8137.2010.03334.x

Paradis, E., Claude, J., & Strimmer, K. (2004). APE: Analyses of phylogenetics and evolution in R language. Bioinformatics, 20(2), 289–290. doi:10.1093/bioinformatics/btg412

Pärtel, M., Öpik, M., Moora, M., Tedersoo, L., Szava-Kovats, R., Rosendahl, S., … Zobel, M. (2017). Historical biome distribution and recent human disturbance shape the diversity of arbuscular mycorrhizal fungi. New Phytologist, 216(1), 227–238. doi:10.1111/nph.14695

Perez-Lamarque, B., & Morlon, H. (2019). Characterizing symbiont inheritance during host–microbiota evolution: Application to the great apes gut microbiota. Molecular Ecology Resources, 19(6), 1659–1671. doi:10.1111/1755-0998.13063

Perez-Lamarque, B., Selosse, M. A., Öpik, M., Morlon, H., & Martos, F. (2020). Cheating in arbuscular mycorrhizal mutualism: a network and phylogenetic analysis of mycoheterotrophy. New Phytologist, 226(6), 1822–1835. doi:10.1111/nph.16474

Pons, J., Barraclough, T. G., Gomez-Zurita, J., Cardoso, A., Duran, D. P., Hazell, S., … Vogler, A. P. (2006). Sequence-based species delimitation for the DNA taxonomy of undescribed insects. Systematic Biology, 55(4), 595–609. doi:10.1080/10635150600852011

Powell, J. R., Monaghan, M. T., Öpik, M., & Rillig, M. C. (2011). Evolutionary criteria outperform operational approaches in producing ecologically relevant fungal species inventories. Molecular Ecology, 20(3), 655–666. doi:10.1111/j.1365-294X.2010.04964.x

Quince, C., Curtis, T. P., & Sloan, W. T. (2008). The rational exploration of microbial diversity. ISME Journal, 2(10), 997–1006. doi:10.1038/ismej.2008.69

R Core Team. (2020). R: A language and environment for statistical computing. Vienna, Austria: R Foundation for Statistical Computing.

Rabosky, D. L. (2009). Ecological limits and diversification rate: Alternative paradigms to explain the variation in species richness among clades and regions. Ecology Letters, 12(8), 735–743. doi:10.1111/j.1461-0248.2009.01333.x

Rabosky, D. L. (2016). Challenges in the estimation of extinction from molecular phylogenies: A response to Beaulieu and O’Meara. Evolution, 70(1), 218–228. doi:10.1111/evo.12820

Rabosky, D. L., & Lovette, I. J. (2008). Density-dependent diversification in North American wood warblers. Proceedings of the Royal Society B: Biological Sciences, 275(1649), 2363–2371. doi:10.1098/rspb.2008.0630

Ramírez-Barahona, S., Sauquet, H., & Magallón, S. (2020). The delayed and geographically heterogeneous diversification of flowering plant families. Nature Ecology and Evolution, 4(9), 1232–1238. doi:10.1038/s41559-020-1241-3

Read, D. J. (1991). Mycorrhizas in ecosystems. Experientia, 47(4), 376–391. doi:10.1007/BF01972080

Revell, L. J. (2012). phytools: An R package for phylogenetic comparative biology (and other things). Methods in Ecology and Evolution, 3(2), 217–223. doi:10.1111/j.2041-210X.2011.00169.x

Rich, M. K., Nouri, E., Courty, P.-E., & Reinhardt, D. (2017). Diet of arbuscular mycorrhizal fungi: Bread and butter? Trends in Plant Science, 22(8), 652–660. doi:10.1016/j.tplants.2017.05.008

Rimington, W. R., Pressel, S., Duckett, J. G., Field, K. J., Read, D. J., & Bidartondo, M. I. (2018). Ancient plants with ancient fungi: liverworts associate with early-diverging arbuscular mycorrhizal fungi. Proceedings of the Royal Society B: Biological Sciences, 285(1888), 20181600. doi:10.1098/rspb.2018.1600

Rolland, J., Condamine, F. L., Jiguet, F., & Morlon, H. (2014). Faster speciation and reduced extinction in the tropics contribute to the mammalian latitudinal diversity gradient. PLoS Biology, 12(1), e1001775. doi:10.1371/journal.pbio.1001775

Royer, D. L., Berner, R. A., Montañez, I. P., Tabor, N. J., & Beerling, D. J. (2004). CO2 as a primary driver of Phanerozoic climate. GSA Today, 14(3), 4. doi:10.1130/1052-5173(2004)014<4:CAAPDO>2.0.CO;2

Säle, V., Palenzuela, J., Azcón-Aguilar, C., Sánchez-Castro, I., da Silva, G. A., Seitz, B., … Oehl, F. (2021). Ancient lineages of arbuscular mycorrhizal fungi provide little plant benefit. Mycorrhiza, 1–18. doi:10.1007/s00572-021-01042-5

Sanders, I. R. (2003). Preference, specificity and cheating in the arbuscular mycorrhizal symbiosis. Trends in Plant Science, 8(4), 143–145. doi:10.1016/S1360-1385(03)00012-8

Sauquet, H., & Magallón, S. (2018). Key questions and challenges in angiosperm macroevolution. New Phytologist, 219(4), 1170–1187. doi:10.1111/nph.15104

Savary, R., Masclaux, F. G., Wyss, T., Droh, G., Cruz Corella, J., Machado, A. P., … Sanders, I. R. (2018). A population genomics approach shows widespread geographical distribution of cryptic genomic forms of the symbiotic fungus Rhizophagus irregularis. ISME Journal, 12(1), 17–30. doi:10.1038/ismej.2017.153

Selosse, M.-A., & Le Tacon, F. (1998). The land flora: a phototroph-fungus partnership? Trends in Ecology & Evolution, 13(1), 15–20. doi:10.1016/S0169-5347(97)01230-5

Sepp, S. K., Davison, J., Jairus, T., Vasar, M., Moora, M., Zobel, M., & Öpik, M. (2019). Non-random association patterns in a plant–mycorrhizal fungal network reveal host–symbiont specificity. Molecular Ecology, 28(2), 365–378. doi:10.1111/mec.14924

Simon, L., Bousquet, J., Lévesque, R. C., & Lalonde, M. (1993). Origin and diversification of endomycorrhizal fungi and coincidence with vascular land plants. Nature, 363(6424), 67–69. doi:10.1038/363067a0

Simon, L., Lalonde, M., & Bruns, T. D. (1992). Specific amplification of 18S fungal ribosomal genes from vesicular-arbuscular endomycorrhizal fungi colonizing roots. Applied and Environmental Microbiology, 58(1), 291–5.

Simpson, G. G. (1953). The major features of evolution. New York: Columbia University Press.

Smith, S. E., & Read, D. J. (2008). Mycorrhizal Symbiosis. Mycorrhizal Symbiosis. Elsevier. doi:10.1016/B978-0-12-370526-6.X5001-6

Stadler, T. (2011). Mammalian phylogeny reveals recent diversification rate shifts. Proceedings of the National Academy of Sciences, 108(15), 6187–6192. doi:10.1073/pnas.1016876108

Strullu-Derrien, C., Selosse, M.-A. A., Kenrick, P., & Martin, F. M. (2018). The origin and evolution of mycorrhizal symbioses: from palaeomycology to phylogenomics. New Phytologist, 220(4), 1012–1030. doi:10.1111/nph.15076

Stürmer, S. L. (2012). A history of the taxonomy and systematics of arbuscular mycorrhizal fungi belonging to the phylum Glomeromycota. Mycorrhiza, 22(4), 247– 258. doi:10.1007/s00572-012-0432-4

Taberlet, P., Bonin, A., Zinger, L., & Coissac, E. (2018). DNA amplification and multiplexing. In Environmental DNA (Oxford Uni, pp. 41–57).

Tajima, F. (1983). Evolutionary relationship of DNA sequences in finite populations. Genetics, 105(2), 437–460.

Tedersoo, L., Albertsen, M., Anslan, S., & Callahan, B. (2021). Perspectives and benefits of high-throughput long-read sequencing in microbial ecology. Applied and Environmental Microbiology, 87(17), 1–19. doi:10.1128/AEM.00626-21

Tisserant, E., Malbreil, M., Kuo, A., Kohler, A., Symeonidi, A., Balestrini, R., … Martin, F. (2013). Genome of an arbuscular mycorrhizal fungus provides insight into the oldest plant symbiosis. Proceedings of the National Academy of Sciences of the United States of America, 110(50), 20117–20122. doi:10.1073/pnas.1313452110

Toussaint, A., Bueno, G., Davison, J., Moora, M., Tedersoo, L., Zobel, M., … Pärtel, M. (2020). Asymmetric patterns of global diversity among plants and mycorrhizal fungi. Journal of Vegetation Science, 31(2), 355–366. doi:10.1111/jvs.12837

Upham, N. S., Esselstyn, J. A., & Jetz, W. (2019). Inferring the mammal tree: Species-level sets of phylogenies for questions in ecology, evolution, and conservation. PLoS Biology, 17(12), e3000494. doi:10.1371/journal.pbio.3000494

van der Heijden, M. G. A. A., Martin, F. M., Selosse, M.-A. A., & Sanders, I. R. (2015). Mycorrhizal ecology and evolution: the past, the present, and the future. New Phytologist, 205(4), 1406–1423. doi:10.1111/nph.13288

Varga, T., Krizsán, K., Földi, C., Dima, B., Sánchez-García, M., Sánchez-Ramírez, S., … Nagy, L. G. (2019). Megaphylogeny resolves global patterns of mushroom evolution. Nature Ecology & Evolution, 3(4), 668–678. doi:10.1038/s41559-019-0834-1

Venice, F., Ghignone, S., Salvioli di Fossalunga, A., Amselem, J., Novero, M., Xianan, X., … Bonfante, P. (2020). At the nexus of three kingdoms: the genome of the mycorrhizal fungus Gigaspora margarita provides insights into plant, endobacterial and fungal interactions. Environmental Microbiology, 22(1), 122–141. doi:10.1111/1462-2920.14827

Werner, G. D. A., Cornelissen, J. H. C., Cornwell, W. K., Soudzilovskaia, N. A., Kattge, J., West, S. A., & Kiers, E. T. (2018). Symbiont switching and alternative resource acquisition strategies drive mutualism breakdown. Proceedings of the National Academy of Sciences, 115(20), 5229–5234. doi:10.1073/pnas.1721629115

Werner, G. D. A., Cornwell, W. K., Sprent, J. I., Kattge, J., & Kiers, E. T. (2014). A single evolutionary innovation drives the deep evolution of symbiotic N2-fixation in angiosperms. Nature Communications, 5(1), 4087. doi:10.1038/ncomms5087

Yildirir, G., Malar, M., Kokkoris, V., & Corradi, N. (2020). Parasexual and sexual reproduction in arbuscular mycorrhizal fungi: Room for both. Trends in Microbiology, 28(7), 517–519. doi:10.1016/j.tim.2020.03.013

Zanne, A. E., Tank, D. C., Cornwell, W. K., Eastman, J. M., Smith, S. A., FitzJohn, R. G., … Beaulieu, J. M. (2014). Three keys to the radiation of angiosperms into freezing environments. Nature, 506(7486), 89–92. doi:10.1038/nature12872

Ziegler, A. M., Eshel, G., McAllister Rees, P., Rothfus, T. A., Rowley, D. B., & Sunderlin, D. (2003). Tracing the tropics across land and sea: Permian to present. Lethaia, 36(3), 227–254. doi:10.1080/00241160310004657

