## Supplementary for "Global drivers of obligate mycorrhizal symbionts diversification"

### Analyzing diversification dynamics using barcoding data: the case of obligate mycorrhizal symbionts

**Supplementary Methods: 1-8**
**Supplementary Notes: 1-3**
**Supplementary Tables: 1-7**
**Supplementary Figures: 1-40**
**Supplementary References**

#### Supplementary Methods:

#### Supplementary Methods 1: Bayesian phylogenetic reconstructions

The VT alignment contains 450 segregating sites within the 520 bp SSU marker, which indicates that this SSU marker is relatively variable across Glomeromycotina. Prior to the VT phylogenetic analysis, we used ModelFinder (Kalyaanamoorthy, Minh, Wong, von Haeseler, & Jermiin, 2017) to select the best substitution model and performed nested sampling analyses (NS (Russel, Brewer, Klaere, & Bouckaert, 2019) in BEAST2) to select the most accurate models among strict, relaxed log-normal, or exponential molecular clocks and coalescent, pure-birth, or birth-death branching process priors. As a result, we generated an input file using BEAUti with the following parameters: a GTR model with 4 classes of rates and invariant sites, a pure birth prior, and relaxed log-normal clock (Supplementary Table 7). The selection of a pure birth prior over a birth death prior by the NS analyses suggested that there is low support for extinction, and this differed from the phylogenetic analyses of Davison *et al.* (2015) who selected a coalescent prior (which tends to push the nodes close to the present compared to a pure birth prior).

In order to efficiently explore tree space, reduce computation time, and improve the robustness of our phylogenies, we constrained the monophyly of the orders Diversisporales, Glomerales, and Paraglomerales + Archaeosporales, based on preliminary phylogenetic reconstructions that supported their monophyly and previous phylogenetic reconstructions of the Glomeromycotina subphylum (Davison et al., 2015; Rimington et al., 2018). We ran BEAST2 to generate a posterior distribution of ultrametric trees sampled using Markov chain Monte Carlo (MCMC) with 4 independent chains each composed of 200,000,000 steps sampled every 20,000 generations. The convergences of the 4 chains were checked using Tracer (Rambaut, Drummond, Xie, Baele, & Suchard, 2018). We used LogCombiner to merge the results setting a 25% burn-in and TreeAnnotator to obtain a maximum clade credibility tree with median branch lengths, hereafter referred to as the VT consensus tree. We also selected 12 trees equally spaced in the 4 independent chains (3 trees per chain) to account for phylogenetic uncertainty in the subsequent diversification analyses, hereafter referred to as the VT replicate trees. Phylogenetic trees were visualized using FigTree (v1.5) and potential negative branch lengths in the consensus tree were set equal to 0 using R (R Core Team, 2020).

Finally, we converted relative branch lengths to absolute time by setting the crown root age at 505 Myr (Davison et al., 2015), which is coherent with fossil data and previous dated molecular phylogenies (Lutzoni et al., 2018; Strullu-Derrien, Selosse, Kenrick, & Martin, 2018).

We used a similar procedure for reconstructing the EU phylogenetic trees and the phylogenetic trees of the clades investigated using GMYC.

#### Supplementary Methods 2: Delineating Evolutionary units

To delineate Glomeromycotina sequences into Evolutionary units, we gathered 41,989 Glomeromycotina sequences from the MaarjAM database accessed in June 2019, from which we filtered 36,411 sequences corresponding to the 18S SSU rRNA gene. In MaarjAM, most of these fungal sequences are assigned to a Glomeromycotina order (Diversisporales, Glomerales, Paraglomerales or Archaeosporales), a Glomeromycotina family, and are mapped to one of the 384 VT. We first built the phylogenetic tree of all the sequences from dataset 1: we aligned the 36,411 sequences using MAFFT with the VT representative sequences as a backbone, filtered out gaps present in more than 50% of the sequences using TrimAl (Capella-Gutierrez, Silla-Martinez, & Gabaldon, 2009), removed sequences having more than 25% of gaps, and selected the 520 base pair barcode region. We obtained a total of 34,205 sequences reduced to 27,728 after removing identical sequences using the GetHaplo function from the R-package *sidier* (Muñoz-Pajares, 2013). We used FastTree (Price, Dehal, &

Arkin, 2009) to build an uncalibrated tree of all Glomeromycotina sequences constrained with the VT consensus tree (option *-constraints*), that we rooted using midpoint rooting in R.

Next, we wrote an R algorithm, *delineate_phylotypes* available in the R-package RPANDA (Morlon et al., 2016), which traverses a tree from the root to the tips, at every node computes the average similarity of all sequences descending from the node, and collapses the sequences into a single EU if their sequence dissimilarity is lower than a given threshold (Morlon et al., 2015). To compute average pairwise similarity among sequences, we used the formula for estimating nucleotide diversity of Ferretti *et al.* (2012) which has the advantage of accounting for gaps in the alignment (Supplementary Method 8). We used such a monophyly criterion, as incorporating phylogeny into Glomeromycotina molecular delineations seems to give more accurate units (Powell, Monaghan, Öpik, & Rillig, 2011). We ran this algorithm on the tree of Glomeromycotina sequences for 97, 97.5, 98, 98.5, and 99% average sequence similarity thresholds, resulting in the delineation of EUs denoted EU97, EU97.5, EU98, EU98.5, and EU99. We chose similarity thresholds higher than 97% as this threshold used in VT has been debated and some authors suggested it might merge several biological species in the same unit (Bruns, Corradi, Redecker, Taylor, & Öpik, 2018).

Finally, we performed Bayesian phylogenetic reconstructions of the EUs. We removed singletons (EUs represented by a unique sequence) as they mostly corresponded to sequences with many gaps and/or badly placed tips in the phylogenetic tree. We assigned to each EU a representative sequence corresponding to its longest available sequence. We performed the same BEAST2 Bayesian phylogenetic reconstructions as the one used to build the VT trees, resulting in a dated consensus tree and 12 replicate trees for each delineation threshold.

#### Supplementary Methods 3: Estimating global Glomeromycotina species diversity:

Before performing diversification analyses, we evaluated how thoroughly sampled our species-level Glomeromycotina phylogenetic trees are by estimating the total number of VT and EUs. First, we constructed rarefaction curves of the number of VT or EUs as a function of the number of sequences and estimated the total number of units using Chao2 index (specpool function, vegan R-package). Second, we used the Bayesian Diversity Estimation Software (BDES (Quince, Curtis, & Sloan, 2008)), which estimates the total number of species by extrapolating sampled taxa abundance distributions. For every delineation, we generated global-scale taxon abundance distributions and assumed that they followed either a log-normal, log-Student, inverse Gaussian, or Sichel distribution (Lewitus, Bittner, Malviya, Bowler, & Morlon, 2018). For each distribution, we ran three independent MCMC chains of 250,000 steps complemented by non-informative prior distributions from trial MCMC runs. The total number of units was estimated by the median value of the last 150,000 steps in the three chains and selected the most likely distribution according to the lowest deviance information criterion (DIC=-2*log(likelihood)+number of parameters).

When performing the EU and GMYC delineations, we discarded all singletons as they can represent a substantial number of units and mostly result from short, badly aligned sequences, and/or wrongly-placed sequences in the phylogenetic tree. As most of the singletons are likely artifacts rather than "biological units" discarding them is a classical approach in such molecular-based delineations from metabarcoding data (Edgar, 2013). However, their absence in the computation of the sampling fraction could bias our estimates. We evaluated this potential bias in the case of GMYC delineation, as this is more tractable computationally. We compared the sampling fractions estimated with or without singletons, and found that discarding singletons only marginally bias the estimations of the sampling fractions, which are always larger than 85% (Supplementary Table 5). Thus, to account for this uncertainly in the estimation of the sampling fraction, we also performed all the diversification analyses on a range of sampling fractions from 50% to 90%.

#### Supplementary Methods 4: Testing the robustness of temporal declines in speciation rates to the presence of high levels of extinction

We found no (or low) support for extinction in the Glomeromycotina phylogenies across all species delineations, sampling fractions, and phylogenetic replicates. However, extinction is notoriously difficult to estimate from reconstructed trees (Rabosky, 2016). We thus tested the robustness of the observed temporal decline in speciation rates under scenarios of high extinction rates. To do so, we considered the best-fit time-dependent model in RPANDA (*i.e.* a model with an exponential variation of the speciation rates through time; see Supplementary Fig. 13) and carried two types of analyses.

First, we fixed positive extinction rates (0.001, 0.005, 0.01, 0.02, or 0.03 extinction events/lineage/Myr) and investigated how the temporal trend in speciation varied with these increasing levels of extinction. Second, we explored models congruent (i.e. with the same likelihood, Louca & Pennell, 2020) to our best-fit model, but with arbitrarily high extinction rates (0.001, 0.005, 0.01, 0.02, and 0.03 extinction events/lineage/Myr), following Morlon, Robin, & Hartig (2022).

#### Supplementary Methods 5: Testing the support for temperature-dependency

In order to check that temperature-dependent models were not artifactually selected, we first simulated 100 phylogenetic trees, for each Glomeromycotina delineation, under the best-supported time-dependent model with associated parameters (estimated by maximum likelihood on the consensus tree) and compared the statistical support for time-dependent and temperature-dependent models fitted to these simulated trees. If temperature-dependency is not artifactually selected on these simulated trees, it should not be artifactually selected on our empirical trees either (Lewitus & Morlon, 2018).

Second, following Clavel & Morlon (2017) and Condamine *et al.* (2019), we tested whether a support for temperature-dependent models could be due to a global temporal trend rather than to details in the temperature curve by performing our model comparison while increasingly smoothing temperature curves using cubic splines with a range of degrees of freedom from 20 to 2. If support for temperature-dependent models disappears when the temperature curve is smoothed, this means that support for temperature dependency is not linked only to a global temporal trend.

Finally, ClaDS analyses revealed that the Glomeromycotina lineages have a relatively high heterogeneity of diversification rates across families. However, the models in RPANDA that assess the effect of environmental dependence are homogeneous models, i.e. they assume that rates are identical across lineages at any given time. In order to check that temperature-dependent models were not artifactually selected because of unaccounted for rate heterogeneity, we simulated 100 phylogenetic trees, for each delineation, under the ClaDS model with heterogeneity in rates according to the hyperparameters estimated on each consensus Glomeromycotina phylogeny (function sim_ClaDS in RPANDA). Then, we compared the statistical support in RPANDA for time-dependent and temperature-dependent models fitted to these simulated trees with heterogeneity in rates across lineages. If temperature-dependency is not artifactually selected on these simulated trees, it should not be artifactually selected on our empirical trees either.

#### Supplementary Methods 6: Estimating land plant fossil diversity

We estimated land plant diversity by using all available land plant fossils in the last 500 Myr by screening all the "Embryophyta" entries the Paleobiology Database (https://paleobiodb.org) accessed on May 4$^{th}$ 2020. Diversity curves were obtained using the shareholder quorum subsampling method (Alroy, 2010) implemented in the pipeline http://fossilworks.org. 17,697 fossil occurrences were used from 880 publications and 3,203 collections. Estimates were performed at the level of land plant genera, with a number of subsampling trials of 100. We tested the effect of different quorum values from 0.2 to 0.7 (Supplementary Fig. 20). As they all suggested similar trends, we kept the quorum value of 0.5 for the main analyses.

We acknowledge that the fossil record of land plants over such a large period of time is discontinuous and patchy. We see our estimates as reflecting the general relative tendency of the land plant diversity through time rather than precise estimates of this diversity.

From Supplementary Fig. 20d, some variations could be attributed to major events, such as mass extinctions and are coherent with previous findings (Silvestro, Cascales-Miñana, Bacon, & Antonelli, 2015). For instance, we can see a sharp decrease of the land plant fossil diversity following the K-Pg extinction (66 Myr ago) or the Permian-Triassic extinction event (250 Myr ago). The early Cretaceous (from -145 Myr) is also characterized by a large increase of the land plant diversity that likely corresponds to the radiation of Angiosperms, that progressively replaced some clades of ferns and Gymnosperms including the Corystospermales, Caytoniales, and Pentoxylales that are now extinct. These lineages were mainly associated with arbuscular mycorrhizal fungi, whereas the Angiosperms and the Gymnosperms that radiated in the last 100 Myr (including Pinaceae) developed a range of alternative nutritive strategies, including symbiont shifting by forming ectomycorrhiza.

Overall, these trends were similar to previous estimates of land plant diversity through time (Cleal & Cascales-Miñana, 2014; Niklas, 1988; Niklas, Tiffney, & Knoll, 1983), although they estimated an even sharper increase of land plant diversity close to the present due to the radiation of Angiosperms (see Fig. 1 in Cleal & Cascales-Miñana, 2014).

#### Supplementary Methods 7: Characterizing Glomeromycotina niche width

We characterized Glomeromycotina relative niche width using a set of 10 abiotic and biotic variables recorded for each Glomeromycotina unit (VT or EUs).

First of all, among dataset 1, given that some interactions were characterized in disturbed ecosystems (e.g. agricultural systems, urban areas, etc.) or sampled directly from soil, we only retained the sequences of Glomeromycotina that occurred in natural ecosystems and interacted with a plant identified at the species level (Öpik et al., 2010) (dataset 2, Supplementary Table 1). This filtering led to 351 fungal VT interacting with a total of 490 plant species (Perez-Lamarque, Selosse, Öpik, Morlon, & Martos, 2020).

Among dataset 2, for each VT or EU unit, we reported the number of continents where it has been sampled, as well as the number of ecosystems, climatic zones, realms, habitats, and biomes defined in MaarjAM (Öpik et al., 2010). We simultaneously reported the number of plant partners that have been recorded (degree of the Glomeromycotina unit) and computed, for each fungal unit, (i) the phylogenetic diversity of these interacting partners (indicating whether a fungus is interacting with closely related plant species or not; R-package *picante* (Kembel et al., 2010)) and (ii) its centrality in the plant-fungus bipartite network (we reported both the closeness and the betweenness centrality; R-package bipartite (Dormann, Gruber, & Fründ, 2008)). The measure of centrality in an interaction network indicates whether a given species is isolated and interacting with few specialist partners (reciprocal specialization – restricted niche) versus part of the interaction core (well-connected generalists interacting with generalists – large niche). For the plants, we used the Plant List (http://www.theplantlist.org) to update some taxonomic assignations following the Angiosperm Phylogeny Group (APG) III system. The land plant phylogenetic tree of the 490 species was obtained by pruning the synthesis phylogeny (Zanne et al., 2014) using Phylomatic (http://phylodiversity.net/phylomatic/) and manually grafting the 37 missing taxa onto the Phylomatic tree as polytomies based on the current literature (Perez-Lamarque et al., 2020). Given that most of these 10 scaled variables are highly correlated, we performed a principal component analysis (PCA), reported the contributions of the different variables to the first axes, and extracted the first two components for each fungal unit that are represented by at least 10 sequences (in dataset 2). We selected the first two axes based on parallel analyses (Buja & Eyuboglu, 1992) using the function *parallel_analysis* from the R-package GeometricMorphometricsMix (Fruciano, 2020). We reported the contributions of the different variables to the first axes and extracted the first two components for each fungal unit that are represented by at least 10 sequences (in dataset 2). PC1 and PC2 explained more than 75% of the variance of the 10 variables (Supplementary Fig. 30). Log-transforming the count variables such as the number of plant species associated with a Glomeromycotina unit did not significantly change the PCA and the downstream analyses.

Importantly, with a dataset containing only 490 plant species, our measure of "Glomeromycotina niche width" actually represents a relative proxy of the complete Glomeromycotina niche width. We considered that a generalist Glomeromycotina unit (*i.e.* a Glomeromycotina unit with a large niche width) is likely to be found associated with many plant species in MaarjAM, whereas a specialist Glomeromycotina unit (*i.e.* a Glomeromycotina unit with a narrow niche width) is more likely to be found associated with a low number of plant species. Given that previous analyses suggested that the relative generalism of Glomeromycotina units does not quantitively change when rarefying the MaarjAM database (Perez-Lamarque et al., 2020), our measure of relative niche width of each Glomeromycotina unit should be reliable despite the relatively low number of plant species in MaarjAM.

### Supplementary Methods 8: Correcting the estimator of genetic diversity for nucleotide alignments with gaps

(Ferretti et al., 2012; Sigwart, 2009; Tajima, 1983, 1989; Watterson, 1975)

Our goal is to correct the estimators of Glomeromycotina genetic diversity ($\theta$) in order to count for gaps in the alignment as technical missing data and not as evolutionary events (indels). We only consider here the single-nucleotide polymorphisms (SNP) as evolutionary events.

Given an alignment $A$ of $n$ haploid individuals with a total sequence length $L$ (i.e. $L$ sites), we are interested in the corrected Tajima's estimator of the nucleotide diversity ($\hat{\theta}_\pi$), such as:

$$\theta = E(\hat{\theta}_{\pi c})$$

Tajima's estimator (Tajima, 1983) corresponds to the average number of nucleotide differences per site between two individuals in all possible pairs. Let's denote $\pi_{i,j}$ the number of nucleotide differences between the $i^{th}$ and $j^{th}$ sequences:

$$\hat{\theta}_\pi = \frac{2}{n(n-1)L} \sum_{i=1}^{n-1} \sum_{j=i+1}^{n} \pi_{i,j}$$

Let $\eta_i$ be the number of sites having one base present in $i$ individuals and another base in $n-i$ individuals, with $1 \leq i \leq [n/2]$. The count of $\eta_i$ corresponds to the site-frequency spectrum, and thus it gives the number of segregated sites:

$$S = \sum_{i=1}^{[n/2]} \eta_i$$

For a given polymorphic site, if one base is present in $i$ individuals and another in $n-i$ individuals, it will be counted as a pairwise difference in $i(n-i)$ pairs of individuals. Thus, instead of calculating the number of pairwise differences by summing $\pi_{i,j}$, we can calculate for every site, the number of pairs of individuals presenting different bases:

$$\hat{\theta}_\pi = \frac{2}{n(n-1)L} \sum_{i=1}^{[n/2]} i\ (n-i)\ \eta_i$$

$$\hat{\theta}_\pi = \frac{2}{n(n-1)L} \sum_{i=1}^{[n/2]} i\ (n-i)\ \eta_i$$

NB: compared to Watterson's estimator ($\hat{\theta}_W$; Watterson, 1975), the sites presenting even nucleotidic frequencies (around $i = [n/2]$) have more weight in the estimator $\hat{\theta}_\pi$. Thus, $\hat{\theta}_\pi$ might be less sensitive to sequencing errors present in only one individual.

Let's define $w_i(n) = \dfrac{2(n-i)}{n(n-1)}$:

$$\hat{\theta}_\pi = \frac{1}{L} \sum_{i=1}^{[n/2]} i \; w_i \; \eta_i$$

Ferreti et al. (2012) proposed a different formula for $\hat{\theta}_\pi$ correcting for the missing data, where each site is represented by $n_x$ (the number of individuals having an informative nucleotide in this site $x$).

$$\hat{\theta}_{\pi c} = \frac{1}{L} \sum_{x=1}^{L} \sum_{i=1}^{[n_x/2]} i \; w_i(n_x) \; \eta_i(x)$$

where $\eta_i(x) = 1$ if there is one base present in $i$ copies and the others in $(n-i)$ copies at the site $x$, and 0 otherwise.

$$\hat{\theta}_{\pi c} = \frac{1}{L} \sum_{x=1}^{L} \sum_{i=1}^{[n_x/2]} i \; w_i(n_x) \; \eta_i(x)$$

$$\hat{\theta}_{\pi c} = \frac{1}{L} \sum_{x=1}^{L} \sum_{i=1}^{[n_x/2]} \frac{2 \; i \; (n_x - i) \; \eta_i(x)}{n_x \; (n_x - 1)}$$

$$\hat{\theta}_{\pi c} = \frac{1}{L} \sum_{x=1}^{L} \sum_{i=1}^{n_x} \frac{i \; (n_x - i) \; \eta_i(x)}{n_x \; (n_x - 1)}$$

Let $j \in \{A, C, G, T\}$ be a particular nucleotide, and $n_{j,x}$ be the number of nucleotides $j$ at site $x$.

$$\hat{\theta}_{\pi c} = \frac{1}{L} \sum_{x=1}^{L} \sum_{j \in \{A,C,G,T\}} \frac{n_{j,x} \; (n_x - n_{j,x})}{n_x \; (n_x - 1)}$$

Finally, as $n_x = \sum_{j \in \{A,C,G,T\}} n_{j,x}$, the corrected Tajima estimator $\hat{\theta}_{\pi c}$ is calculated as:

$$\hat{\theta}_{\pi c} = \frac{1}{L} \sum_{x=1}^{L} \left( 1 - \frac{\displaystyle\sum_{i \in \{A,C,G,T\}} n_{i,x}(n_{i,x} - 1)}{n_x(n_x - 1)} \right)$$

We used this formula to compute the genetic diversity of each Glomeromycotina species represented by at least 10 sequences (in dataset 2). By using a minimal number of 10 sequences, the estimates of genetic diversity were either not significantly or only slightly correlated with the number of sampled sequences per species. This criterion considerably reduced the number of EU99 units represented in the analyses, as many EU99 had less than 10 sequences (Supplementary Table 2).

#### Supplementary Notes:

#### Supplementary Note 1: Glomeromycotina species delineation based on the SSU rRNA gene

The GMYC analyses performed on Glomeromycotina families tend to support a species delineation of Glomeromycotina between EU98.5 and EU99 (Supplementary Table 4). This suggests that the expected number of Glomeromycotina species-like units should be between 1,200 and 2,600 (or between 1,300 and 2,900 when accounting for the sampling fraction >90%). The threshold time found in our GMYC analyses was similar to that found in Powell *et al.* (2011) when comparable (*i.e.* for Diversisporales in the sequences from Estonia; Supplementary Figure 3). Conversely, the VT delineation presents a flexible threshold of delimitation that can vary according to some additional expertise: while VT sequences have at least 97% of similarity, some VT have a finer delineation that are >98.5%.

The majority of the VT units tend to be either present on all continents (24%), or endemic to a unique continent (32%), as previously demonstrated in Davison *et al.* (2015) who highlighted the low endemism of VT. Conversely, EU delineations range from mainly ubiquitous (37% of EU97 are present on all continents) to more endemic units (40% of EU99 were only found on one continent). Our analyses suggest that whereas many VT have a distribution similar to the distribution of EU98.5 or EU99 (relatively similar proportions of VTs shared between continents), some VT merge together too highly dissimilar sequences that likely correspond to different "biological units", and would require a finer delineation: in particular units present in the 6 continents are not very common in the EU98.5 and EU99 delineations. In addition, this was also supported by the fact that some VT present very high values of genetic diversities (results not shown).

Although Glomeromycotina units at 98.5 or 99% tend to be less ubiquitous than VTs, their niches remain particularly large compared to those of their associated plants (Davison et al., 2015), confirming that more finely delineated units might still present relatively low endemism, maybe thanks to long-distance dispersion (Bueno & Moora, 2019).

Thus, although some VT might need to be more finely delineated, many units in the current delineations look fine and this does not seem to impact the previously formulated conclusions about the low endemism of Glomeromycotina (Davison et al., 2015). Such conclusions are also supported by more recent genomic evidence (Savary et al., 2018).

#### Supplementary Note 2: Glomeromycotina phylogenetic trees

Although our Bayesian phylogenetic reconstructions were mainly consistent across delineations (VT or EU) and phylogenetic replicates (consensus tree and 12 independent replicates sampled from the MCMC chains), we noticed that some nodes presented poor resolutions (Supplementary Fig. 4). In particular, the branching of the main Glomeromycotina orders (Paraglomerales, Archaeosporales, Diversisporales, and Glomerales) appeared to be uncertain. Our analyses mainly supported the grouping of Paraglomerales and Archaeosporales together, which is consistent with previous phylogenetic reconstructions of Glomeromycotina (Davison et al., 2015; Rimington et al., 2018), but contradict others where Paraglomerales initially diverged from the other orders (Krüger, Krüger, Walker, Stockinger, & Schüßler, 2012). Besides the Paraglomerales-Archaeosporales uncertainty, the branching time and topology of our phylogenetic trees were overall coherent with the backbone obtained by Lutzoni et al. (2018) (see Supplementary Fig. 2 of Lutzoni et al. (2018)) based on multiple fungal genes. Finally, our branching times were overall older than those of Davison *et al.*

(2015), likely because of the selection of a different branching process prior (Supplementary Table 7 - Davison *et al.* (2015) used an unlikely coalescent prior).

In any case, the topology of the early branching events would certainly benefit from genomic data, that are currently too scarce and do not cover the main Glomeromycotina orders (Venice et al., 2020). To account for this phylogenetic uncertainty in our diversification analyses, we sampled multiple trees per delineations to cover a range of different scenarios are ran all our models on each tree independently.

#### Supplementary Note 3: Limits of phylogenetic-based diversification analyses:

Phylogenetic-based diversification analyses have their limitations, including the difficulty of estimating extinction (Rabosky, 2016) and the associated difficulty of distinguishing between alternative diversification histories (Louca & Pennell, 2020; but see Morlon *et al.*, 2020). In our analyses, we made two specific simplifying assumptions on the extinction rate, by assuming either a constant turnover rate or a constant extinction rate. We also used three different ways to model variations in speciation rates: heritable rates in ClaDS, piecewise-constant rates in CoMET, and continuous time dependencies in RPANDA. In particular, for the time-dependent models in RPANDA, we considered models with constant or exponential variation of speciation rates through time and null or constant extinction rates: models with constant rates correspond to the null hypothesis of clock-like speciation, whereas the exponential variation ensures positive rates and are an approximation of diversity dependence, a process often invoked during radiations (Rabosky & Lovette, 2008). We also further tested the robustness of our findings concerning environmental dependencies by performing additional simulations (Supplementary Methods 5). In addition, we have carried additional analyses testing the robustness of our results concerning the decline in speciation rates to high extinction rates (Supplementary Methods 4). Although this does not exclude the possibility that different results could be found under yet other assumptions, our results were consistent under all the alternative assumptions we made.

Phylogenetic-based diversification analyses are also fundamentally dependent on the robustness and completeness of the phylogenetic data, as well as species delineation, which remain major challenges in microbial groups studied primarily via metabarcoding studies. Additionally, phylogenetic approaches for testing the effect of paleoenvironments on diversification depend on the dating of the environmental variables and the phylogenies, which both have uncertainties. We accounted for these uncertainties as best as we could, and found rather consistent and realistic results. Importantly, carrying such studies is our only possible attempt at characterizing the diversification history of microbial groups of major ecological and evolutionary importance such as Glomeromycotina for which we have very few fossil data.

#### Supplementary Tables:

**Supplementary Table 1: Data selection in the MaarjAM database:**

**(a):** List of sequences from MaarjAM selected for the analyses according to the successive filters.

**(b):** Number of selected sequences per fungal family (dataset 1).

**(c):** Number of selected sequences per continent in natural environments (dataset 2).

**(a)**

| Filters | Number of interactions |
|---|---|
| Initial interactions downloaded from MaarjAM (accessed in June 2019) | 41,989 |
| Interactions with fungal sequences corresponding to 18S rDNA | 36,411 **Dataset 1** **(Use for EU delineations)** |
| Interactions documented directly from the plant root (e.g. discarding soil samples) | 30,337 |
| Interactions sampled in a natural ecosystem (i.e. removing anthropogenic sites) | 27,512 |
| Interactions with an associated plant identified at the species level | 26,350 **Dataset 2** **(Use for statistical analyses)** |

**(b)**

| Glomeromycotina family | Acaulosporaceae | Ambisporaceae | Archaeosporaceae | Claroideoglomeraceae | Diversisporaceae |
|---|---|---|---|---|---|
| Number of sequences | 1,441 | 155 | 963 | 2,201 | 1,091 |
| Glomeromycotina family | Geosiphonaceae | Gigasporaceae | Glomeraceae | Pacisporaceae | Paraglomeraceae |
| Number of sequences | 4 | 887 | 28,896 | 59 | 714 |

**(c)**

| Continent | Africa | Asia | Europe | North America | Oceania | South America |
|---|---|---|---|---|---|---|
| Number of sequences | 3,638 | 3,498 | 11,932 | 966 | 1,583 | 4,733 |

**Supplementary Table 2: Documented plant-Glomeromycotina interactions:** Characteristics of the Glomeromycotina units (VT or EU) in terms of interactions with plants (occurring in natural environment; dataset 2 – see Supplementary Table 1).

| Units | VT | EU97 | EU97.5 | EU98 | EU98.5 | EU99 |
|---|---|---|---|---|---|---|
| Number of units | 384 | 182 | 340 | 641 | 1,190 | 2,647 |
| Number of units in natural environments | 351 | 169 | 302 | 569 | 1,052 | 2,303 |
| Number of units in natural environments with at least 10 sequences | 232 | 112 | 162 | 257 | 355 | 456 |
| Number of interactions in natural environments | 26,350 | 25,024 | 24,961 | 24,813 | 24,286 | 22,518 |
| Number of plant partners | 490 | 479 | 479 | 479 | 478 | 472 |
| Number of unique plant-fungus interactions | 12,914 | 8,460 | 9,361 | 11,231 | 12,837 | 14,017 |

**Supplementary Table 3: Characteristics of the fungal units (VT or EU) in the database**:

Characteristics of the fungal units (VT or EU) according to each species delineation criterion. Discarded sequences correspond to singletons, plus too short or badly aligned sequences that are removed from the alignment before reconstructing the phylogenetic tree used for EU delineation.

| | VT | 97% | 97.5% | 98% | 98.5% | 99% |
|---|---|---|---|---|---|---|
| Number of units | 384 | 182 | 340 | 641 | 1,190 | 2,647 |
| Total number of discarded sequences | 828 | 1,882 | 1,979 | 2,210 | 2,947 | 5,407 |
| Number of discarded singletons | NA | 71 | 168 | 399 | 1,136 | 3,596 |
| Most abundant EU | 1,184 | 3,323 | 3,323 | 2,455 | 1,225 | 743 |
| Mean size | 93 | 190 | 101 | 53 | 28 | 12 |

**Supplementary Table 4: Number of units (VT, EU, or GMYC) per fungal clades.**

For GMYC analyses, the number of units corresponds to the inferred number of clusters (discarding singletons) and the numbers in brackets indicate the corresponding confidence interval at 5%. GMYC analyses were not ran for the Glomeraceae family because of computational limits.

| | VT | 97% | 97.5% | 98% | 98.5% | 99% | GMYC clusters |
|---|---|---|---|---|---|---|---|
| Ambisporaceae | 4 | 1 | 2 | 3 | 5 | 11 | 207 (191-227) |
| Archaeosporaceae | 13 | 7 | 19 | 33 | 49 | 94 | |
| Geosiphonaceae | 1 | 0 | 1 | 1 | 1 | 1 | |
| Paraglomeraceae | 19 | 12 | 12 | 24 | 28 | 59 | |
| Acaulosporaceae | 35 | 6 | 18 | 30 | 60 | 138 | 166 (139-191) |
| Diversisporaceae | 19 | 10 | 11 | 24 | 32 | 79 | |
| Gigasporaceae | 11 | 1 | 15 | 28 | 44 | 89 | |
| Pacisporaceae | 2 | 1 | 1 | 1 | 4 | 5 | |
| Claroideoglomeraceae | 16 | 6 | 12 | 16 | 53 | 119 | 63 (56-128) |
| Glomeraceae | 264 | 138 | 249 | 481 | 914 | 2052 | NA |

**Supplementary Table 5: GMYC delineation and corresponding sampling fraction:**

For each of the 3 tested Glomeromycotina clades, we indicated the initial number of sequences and the number of haplotypes (unique SSU rRNA sequences) on which the GMYC analyses were performed, the inferred number of clusters (excluding singletons; with its associated confidence interval indicated in brackets), the inferred number of entities (clusters plus singletons; with its associated confidence interval indicated in brackets), as well as their corresponding estimated sampling fraction using the Bayesian Diversity Estimation Software (Quince et al., 2008) assuming a Sichel species abundance distribution. The Sichel distribution was selected compared to other distributions (log-normal, log-Student, and inverse gaussian) based on lowest deviance information criterion (DIC).

On average, the GMYC analyses revealed that there are 10 different SSU rRNA haplotypes for one GMYC cluster.

| Glomeromycotina clade | Glomeromycotina family | Number of fungal sequences (and haplotypes) | GMYC clusters | GMYC clusters and singletons | Sampling fraction without singletons | Sampling fraction with singletons |
|---|---|---|---|---|---|---|
| Archaeosporales Paraglomerales | Ambisporaceae Archaeosporaceae Geosiphonaceae Paraglomeraceae | 1,836 (1,493) | 207 (191-227) | 264 (241-294) | 0.96 | 0.86 |
| Diversisporales | Acaulosporaceae Diversisporaceae Gigasporaceae Pacisporaceae | 3,478 (2,765) | 166 (139-191) | 182 (151-212) | 0.99 | 0.96 |
| Claroideo-glomeraceae | Claroideo-glomeraceae | 2,201 (1,686) | 63 (56-128) | 69 (62-142) | 0.97 | 0.93 |

**Supplementary Table 6: The estimated sampling fraction using Chao2 index suggested a sampling fraction >90%:**

For each species delineation (VT or EU), the table indicates the estimated number of units, the sampling fraction, and a confidence interval built from the standard error (s.e.).

| Species delineation | Observed number of units | Chao2 estimates | | | |
|---|---|---|---|---|---|
| | | Estimated number of units | Sampling fraction | Sampling fraction +/- standard error (s.e.) | |
| | | | | -s.e. | +s.e. |
| VT | 384 | 407 | 94% | 92% | 97% |
| EU97 | 182 | 188 | 97% | 94% | 99% |
| EU97.5 | 340 | 356 | 95% | 94% | 97% |
| EU98 | 641 | 663 | 97% | 96% | 98% |
| EU98.5 | 1,190 | 1,239 | 96% | 95% | 97% |
| EU99 | 2,647 | 2,734 | 97% | 96% | 97% |

**Supplementary Table 7: The prior selection for the VT Bayesian phylogenetic reconstructions (BEAST) using nested sampling (NS) favored a log-normal and Pure-birth prior.**

The selected model based on the highest marginal log-likelihood is highlighted in orange: it corresponds to a Pure-birth prior with a log-normal clock model. Preliminary analyses (not shown here) on Glomeromycotina phylogenetic trees reconstructed using a (non-significantly supported) Birth-death prior found similar diversification trends as obtained with a Pure-birth prior (*e.g.* a decline in speciation rates toward the present, estimation extinction rates close to 0…).

| Tree-shape prior | Clock-model | Marginal log-likelihood | Standard deviation |
|---|---|---|---|
| Coalescent | Exponential | -24,982 | 50.7 |
| Birth-death | Exponential | -24,844 | 46.9 |
| Birth-death | Log-normal | -24,763 | 45.3 |
| Birth-death | Strict | -24,956 | 43.0 |
| Pure-birth | Exponential | -24,774 | 49.0 |
| Pure-birth | Log-normal | -24,675 | 43.4 |

### Supplementary Figures:

**Supplementary Figure 1: Visualization of Glomeromycotina sequence alignments: The ITS marker is not a good marker for Glomeromycotina phylogenetic reconstruction compared to the SSU rRNA region:**

Glomeromycotina sequences were downloaded from (Lekberg et al., 2018), clustered into OTUs at 97% using VSEARCH (Rognes, Flouri, Nichols, Quince, & Mahé, 2016), and aligned using MAFFT (Katoh & Standley, 2013).

(a) For the SSU rRNA gene, we obtained 70 OTUs of 280 base pairs on average that almost perfectly align with very few gaps.

(b) Conversely, for the ITS marker, we obtained 76 OTUs of 280 base pairs on average that badly align with very many gaps, such that the total alignment length exceeds 450 base pairs.

The following alignments were visualized using the software AliView (https://ormbunkar.se/aliview/).

**(a)**

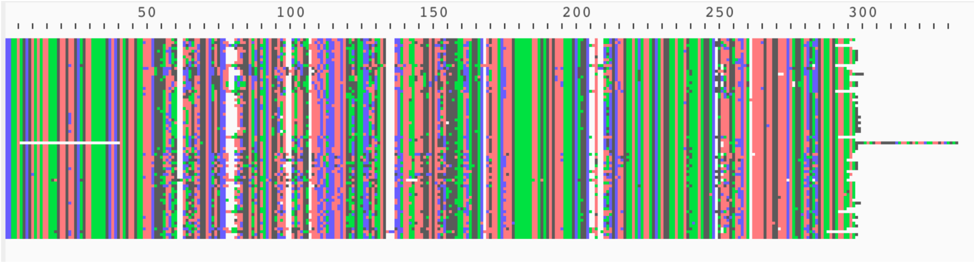


**(b)**

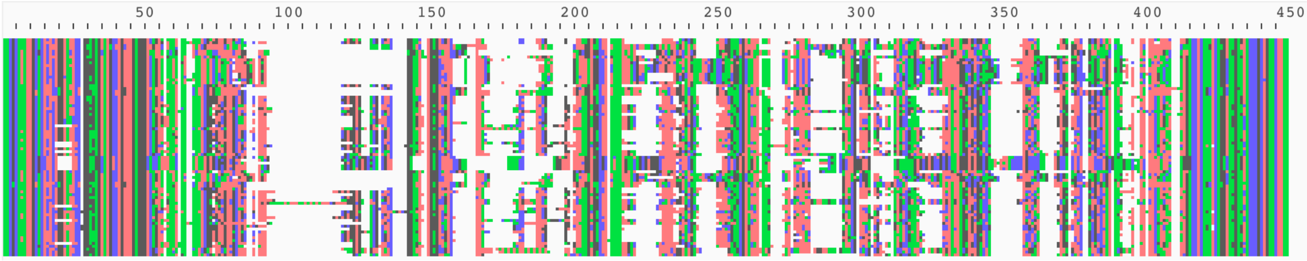

**Supplementary Figure 2: Simulated diversification scenarios:**

**(a) Different simulated diversification scenarios:** To test whether a short and slowly evolving barcoding gene can be used to infer the diversification dynamic of a given clade, we simulated two scenarios of diversification in the last 500 Myr: (1) constant speciation rate and no extinction or (2) constant speciation and extinction rates. Speciation rates are represented in blue and extinction rates in red. The simulated diversification rates were chosen to be close to the rates estimated in Glomeromycotina (Fig. 2) for VT **(case I)** or for EU99 **(case II)** such that we obtained similar numbers of species.

**(b) The EU99 delineation tend to lump together species:** For each scenario, we performed 10 simulations and reported in the table the original number of species in the simulated clade and the evolutionary units at 99% (EU99) delineated from the simulated DNA sequences with several coalescing individuals per species. The number of EU99 is in general lower than the true number of species (i.e. several species are lumped within the same EU99 unit), but in a few cases, the number of EU99 is higher than the true number of species (the high intra-specific differentiation lead to several EU99 units).

**(I) Using a net diversification rate (r=0.01) leading to a total number of species similar to the number of VT:**

(a)

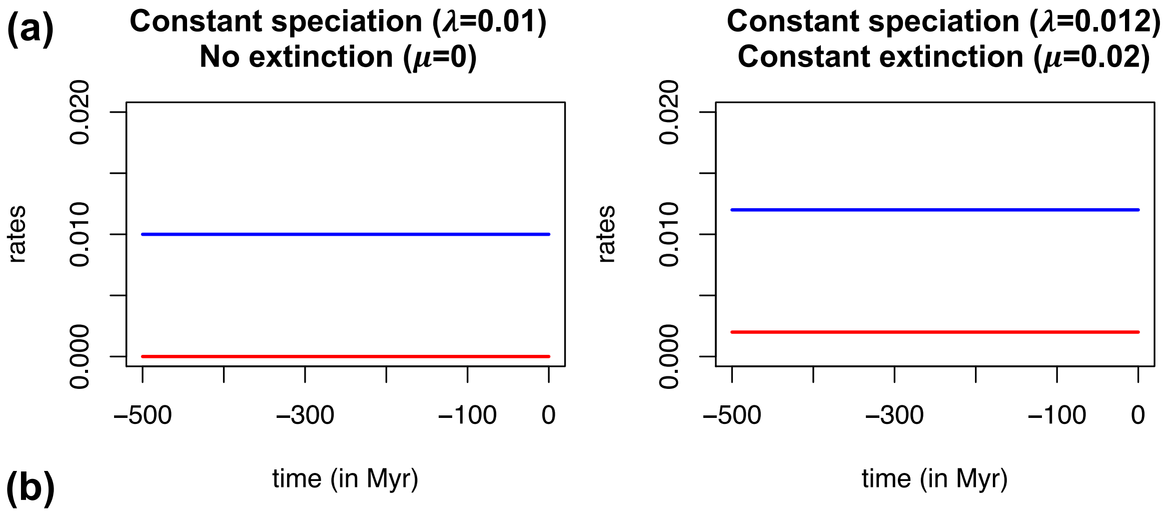


(b)

| Constant speciation ($\lambda$=0.01) No extinction ($\mu$=0) | | Constant speciation ($\lambda$=0.012) Constant extinction ($\mu$=0.02) | |
|---|---|---|---|
| Number of simulated species | Number of EU99 | Number of simulated species | Number of EU99 |
| 75 | 70 | 40 | 32 |
| 64 | 44 | 147 | 133 |
| 270 | 225 | 446 | 372 |
| 360 | 333 | 388 | 338 |
| 96 | 91 | 227 | 201 |
| 542 | 551 | 171 | 141 |
| 291 | 239 | 344 | 308 |
| 148 | 125 | 14 | 17 |
| 466 | 409 | 495 | 381 |
| 187 | 156 | 417 | 378 |

**(II) Using a net diversification rate (r=0.015) leading to a total number of species similar to the number of EU99 units:**

(a)

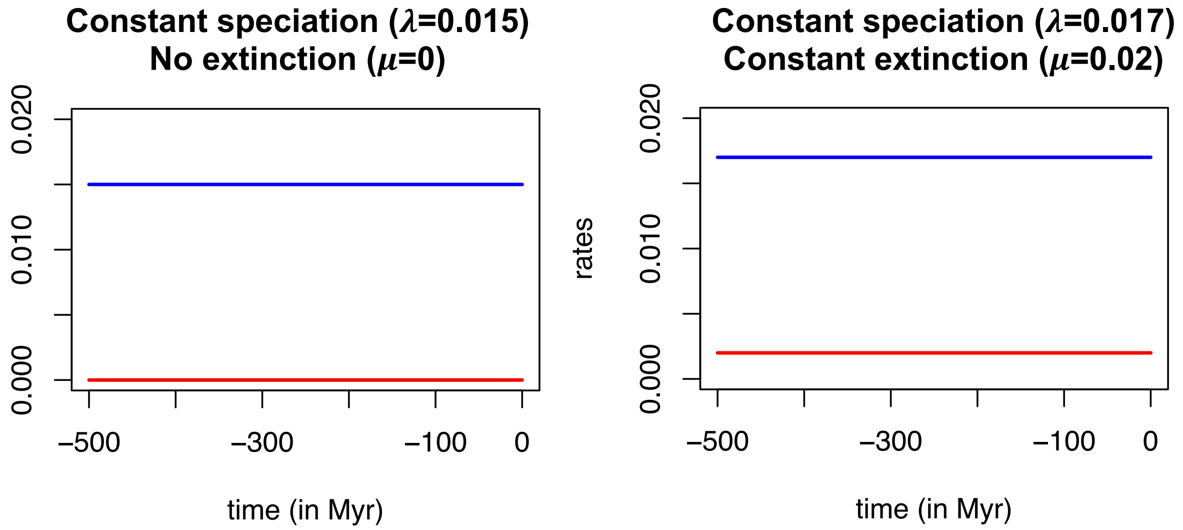


(b)

| Constant speciation ($\lambda$=0.015) No extinction ($\mu$=0) | | Constant speciation ($\lambda$=0.017) Constant extinction ($\mu$=0.02) | |
|---|---|---|---|
| **Number of simulated species** | **Number of EU99** | **Number of simulated species** | **Number of EU99** |
| 2,727 | 2,117 | 1,754 | 1,208 |
| 1,612 | 1,280 | 2,556 | 1,763 |
| 1,366 | 1,099 | 1,334 | 924 |
| 1,585 | 1,251 | 1,509 | 1,117 |
| 1,826 | 1,549 | 2,668 | 2,218 |
| 2,804 | 2,259 | 1,731 | 1,378 |
| 1,402 | 1,070 | 2,453 | 1,720 |
| 1,926 | 1,588 | 2,595 | 1,943 |
| 1,342 | 1,072 | 2,633 | 2,121 |
| 2,120 | 1,674 | 1,855 | 1,349 |

**Supplementary Figure 3: GMYC species delineations in Glomeromycotina clades significantly support the existence of intraspecific haplotypes in the SSU rRNA gene:**

**(a, b, c)** Likelihood of the GMYC model as a function of the relative time threshold *t* that separates the coalescent versus Yule process from -100 (the root age of the Glomeromycotina clade – the age of the root is arbitrarily set at 100) to 0 (present time). The likelihood of the null model considering that each tip of the tree is a species (*i.e.* each SSU rRNA haplotype corresponds to a different Glomeromycotina species) is the likelihood on the right of the plots. A LRT is performed for each tree between the GMYC model with maximum likelihood and the null model. If the LRT supports the GMYC model, different SSU rRNA haplotypes are clustered into the same Glomeromycotina species, *i.e.* the SSU rRNA gene has time to accumulate substitutions between Glomeromycotina speciation events. Alternatively, if the null model cannot be rejected, each SSU rRNA haplotype represents one or multiple Glomeromycotina species.

We performed these analyses on Archaeosporales + Paraglomerales (a – LRT: $P = 4.10^{-15}$), Claroideoglomeraceae (b – LRT: $P < 1.10^{-16}$), and Diversisporales (c – LRT: $P < 1.10^{-16}$).

**(d, e, f)** Intraspecific genetic diversity measured using Tajima's estimator ($\theta\pi$; Supplementary Methods 8) within each GMYC unit. Note that $\theta\pi$ is the average % of nucleotide difference between pairs of sequence, i.e. 100 -% sequence similarity. The median value of the genetic diversities is just below 1% for Archaeosporales + Paraglomerales (d) and just above 1% for Claroideoglomeraceae (e) and Diversisporales (f), confirming that GMYC delineations range on average between the EU98.5 and the EU99 delineations (Supplementary Table 4).

**(g, h, i)** Number of reconstructed fungal lineages through time (lineages through time plots) for the Glomeromycotina phylogenetic trees reconstructed from the GMYC delineations (only 1 representing sequence is kept for each GMYC unit and phylogenetic analyses were performed using the same procedure as described in Supplementary Methods 1).

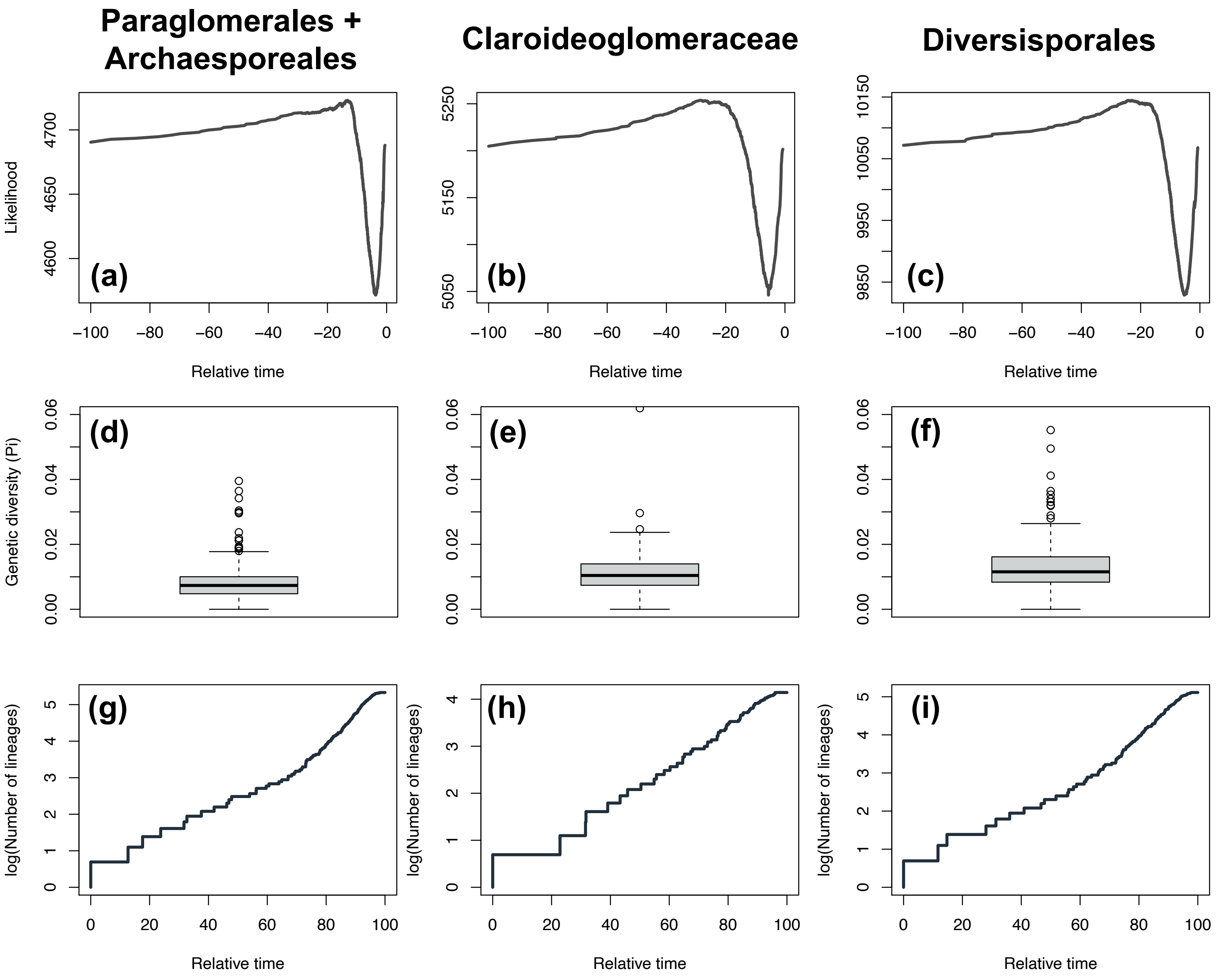

**Supplementary Figure 4: Consensus Glomeromycotina phylogenetic trees for the different species delineations:**

Consensus Glomeromycotina phylogenetic trees using Bayesian reconstruction (BEAST; with a pure-birth prior and a log-normal clock selected using Nested Sampling) for different Glomeromycotina delineations. Values at nodes indicate the Bayesian posterior probability.

Colors highlight the main Glomeromycotina clades: Red = Paraglomerales + Archaeosporales; Green = Diversisporales; Purple = Glomeraceae; Blue = Claroideoglomeraceae.

The topologies of the consensus phylogenetic trees for the main Glomeromycotina clades vary with species delineations; however, this variation reflects the low support of some node rather than an effect of species delineation *per se*. There is indeed as much variation between Bayesian replicates from the same delineation scheme than between delineations schemes. We account for both sources of uncertainty when replicating our analyses across delineation and across replicate trees.

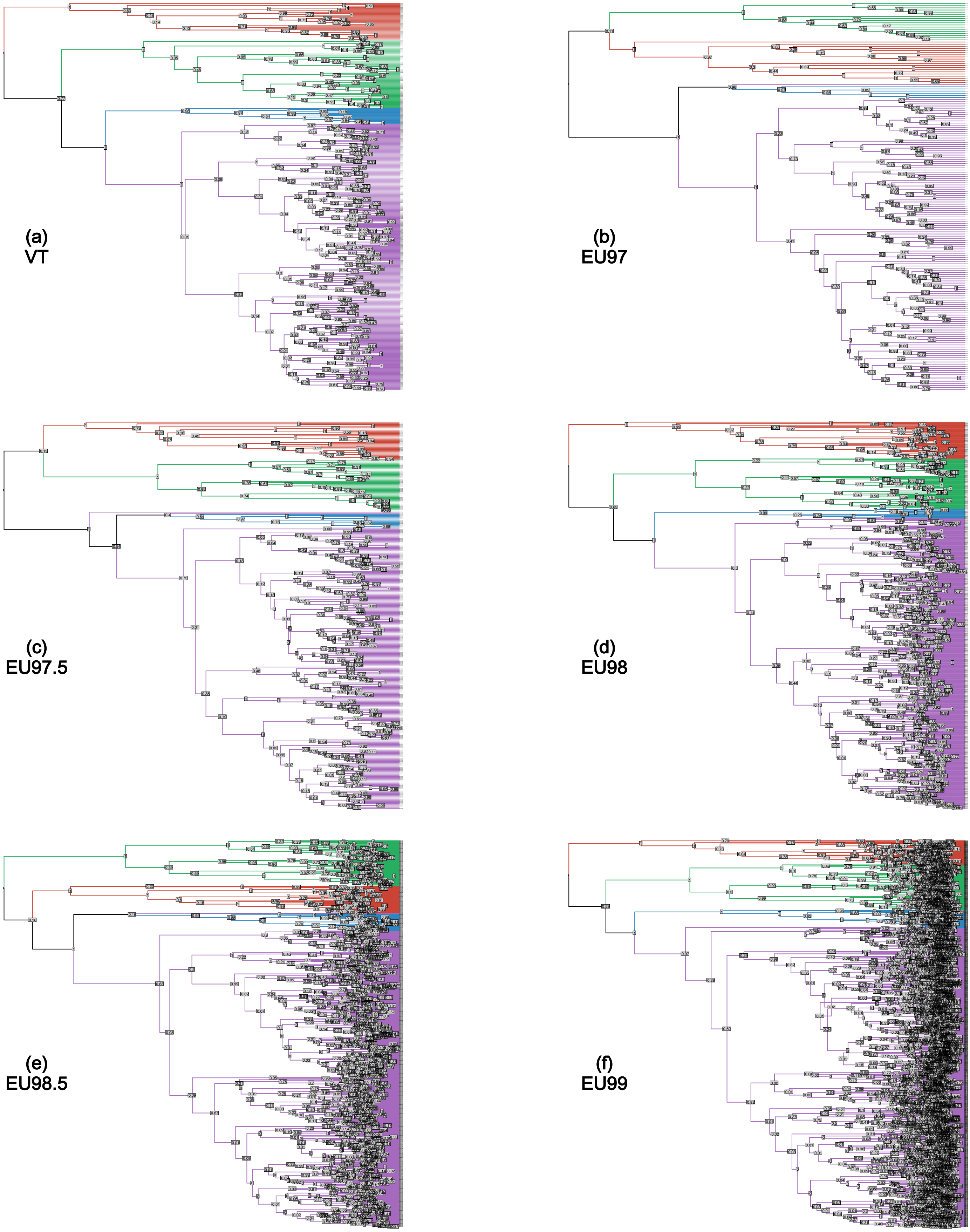
(a)
VT
(b)
EU97
(c)
EU97.5
(d)
EU98
(e)
EU98.5
(f)
EU99

**Supplementary Figure 5: Node depth distribution of the consensus Glomeromycotina phylogenetic trees for the different delineations.**

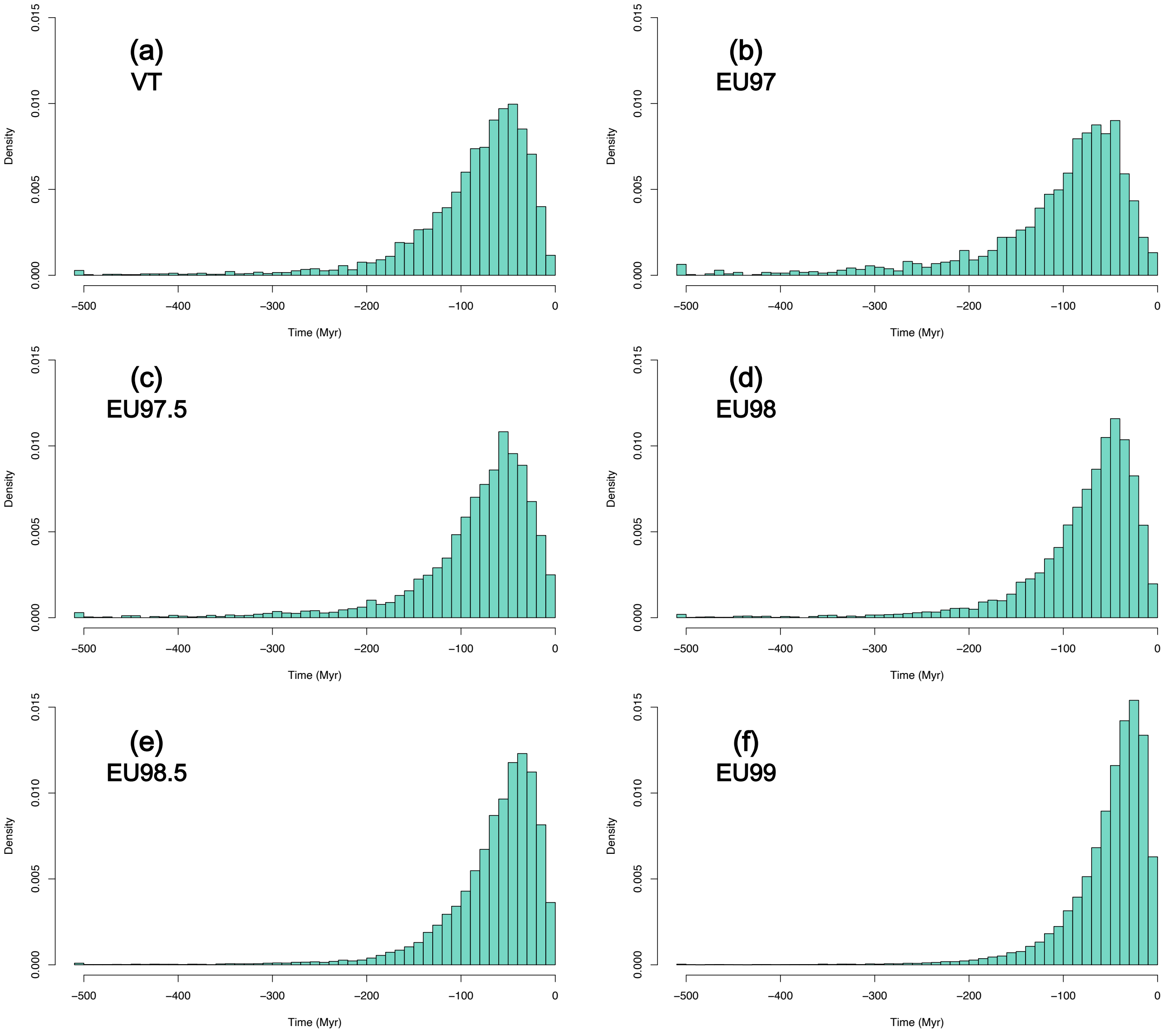

**Supplementary Figure 6: The accumulation of fungal lineages through time present a slowdown toward the present in the reconstructed Glomeromycotina phylogenetic trees:**

Number of reconstructed fungal lineages through time for the different Glomeromycotina phylogenetic trees (lineages through time plots). For the different Glomeromycotina delineations, the consensus tree is in dark grey, whereas orange lines represent the 12 independent replicate trees.

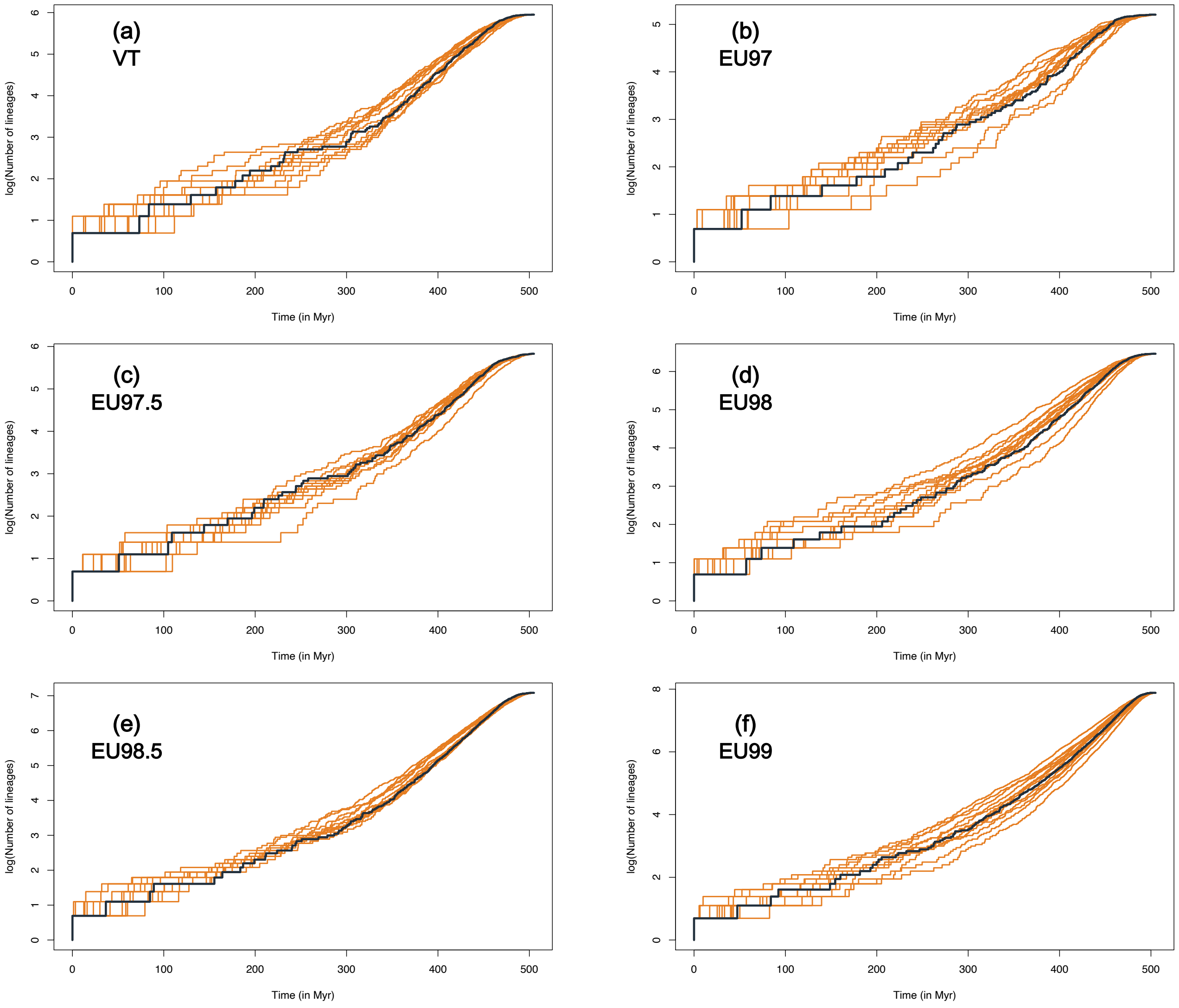

**Supplementary Figure 7: Speciation rates per lineage estimated by ClaDS show that Glomeromycotina experienced heterogeneous diversification rates across clades and time.**

Speciation rates per lineage estimated by ClaDS on the Glomeromycotina consensus trees, for several delineations and with the BDES estimated sampling fraction (Quince et al., 2008).

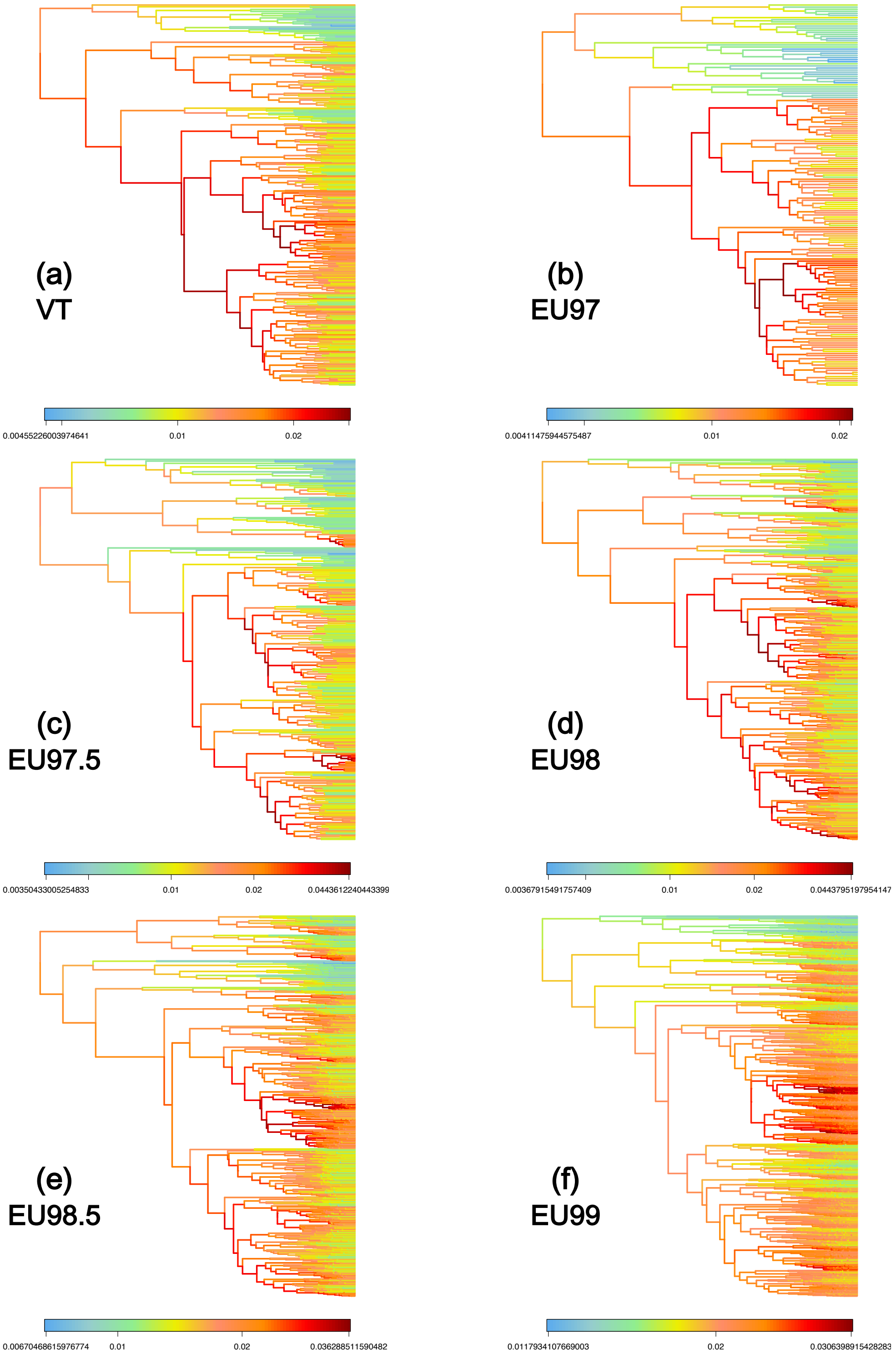

**Supplementary Figure 8: Present-day speciation rates at the tips estimated by ClaDS show that Glomeromycotina have heterogeneous diversification rates across clades.** Speciation rates at the tips estimated by ClaDS for the main Glomeromycotina orders (Archaeosporales, Paraglomerales, Diversisporales, and Glomerales), for several Glomeromycotina delineations and using the BDES estimated sampling fraction.

The difference of speciation rates across Glomeromycotina orders was statistically tested using mixed linear model, with all speciation rates estimates (the consensus tree and the 12 replicate trees) and the Glomeromycotina units as a random factor. Significant relationships are represented in bold at the top of each panel (F-statistic, with the degrees of freedom and the corresponding p-value).

Boxplots indicate the median surrounded by the first and third quartiles, and whiskers extend to the extreme values but no further than 1.5 of the inter-quartile range.

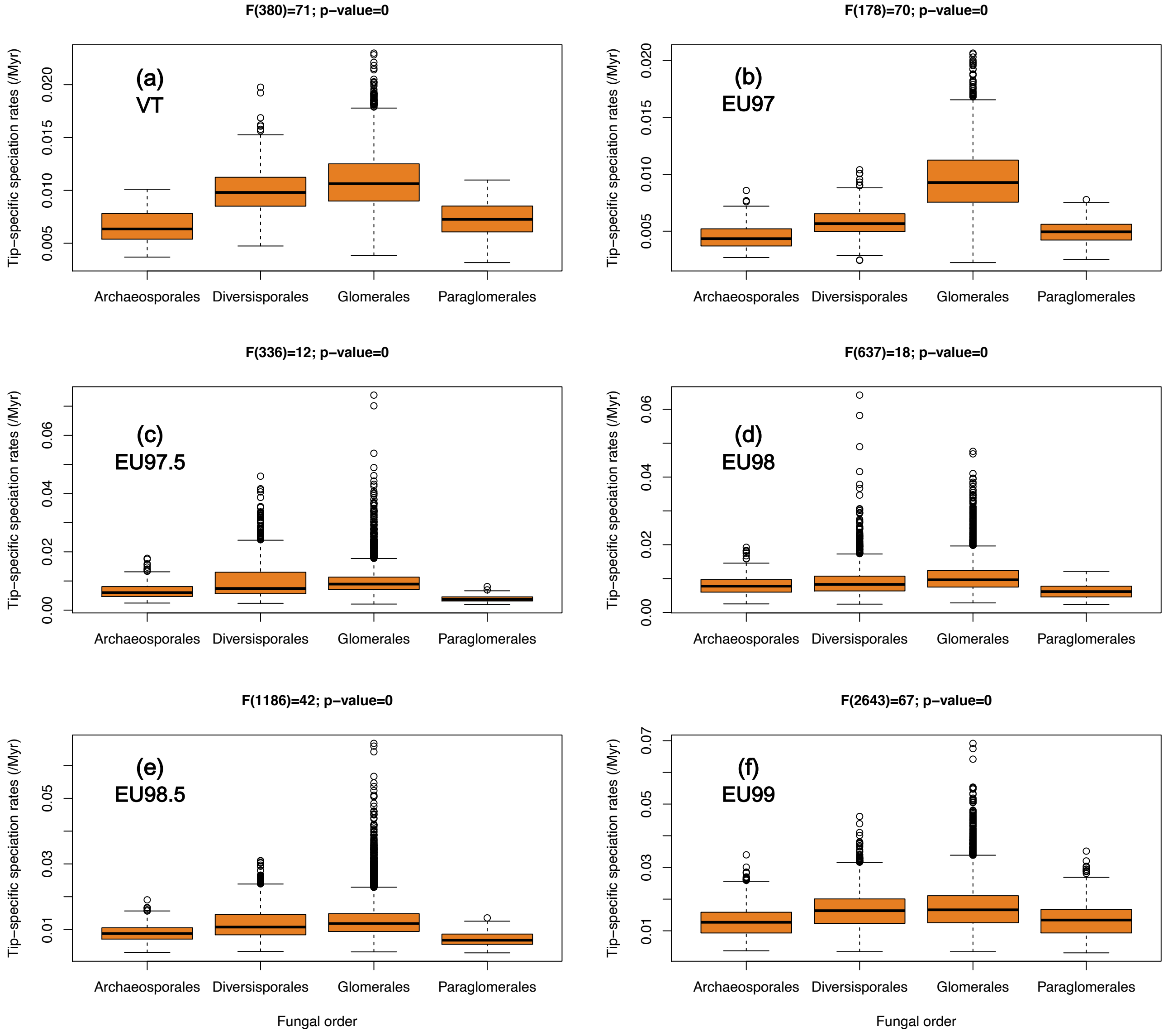

F(380)=71; p−value=0
(a)
VT
F(178)=70; p−value=0
(b)
EU97
F(336)=12; p−value=0
(c)
EU97.5
F(637)=18; p−value=0
(d)
EU98
F(1186)=42; p−value=0
(e)
EU98.5
F(2643)=67; p−value=0
(f)
EU99
Tip−specific speciation rates (/Myr)
Archaeosporales
Diversisporales
Glomerales
Paraglomerales
Fungal order

**Supplementary Figure 9: Speciation rates at the tips estimated by ClaDS for VT and EU delineations are significantly correlated.**

For each Glomeromycotina SSU rRNA haplotype sequences, we represented the estimated speciation rate of its corresponding EU as a function of the estimated speciation rate of its corresponding VT. Ideally, we would expect the estimate to be the same, i.e. on the first bisector (represented in light grey). The estimated relationships between EU and VT speciation rates are indicated in dark grey.

For each delineation, speciation rates were estimated on the consensus phylogenetic tree for each delineation and using the BDES estimated sampling fraction. We attributed the same speciation rate to all the haplotype sequences constituting each VT or EU unit.

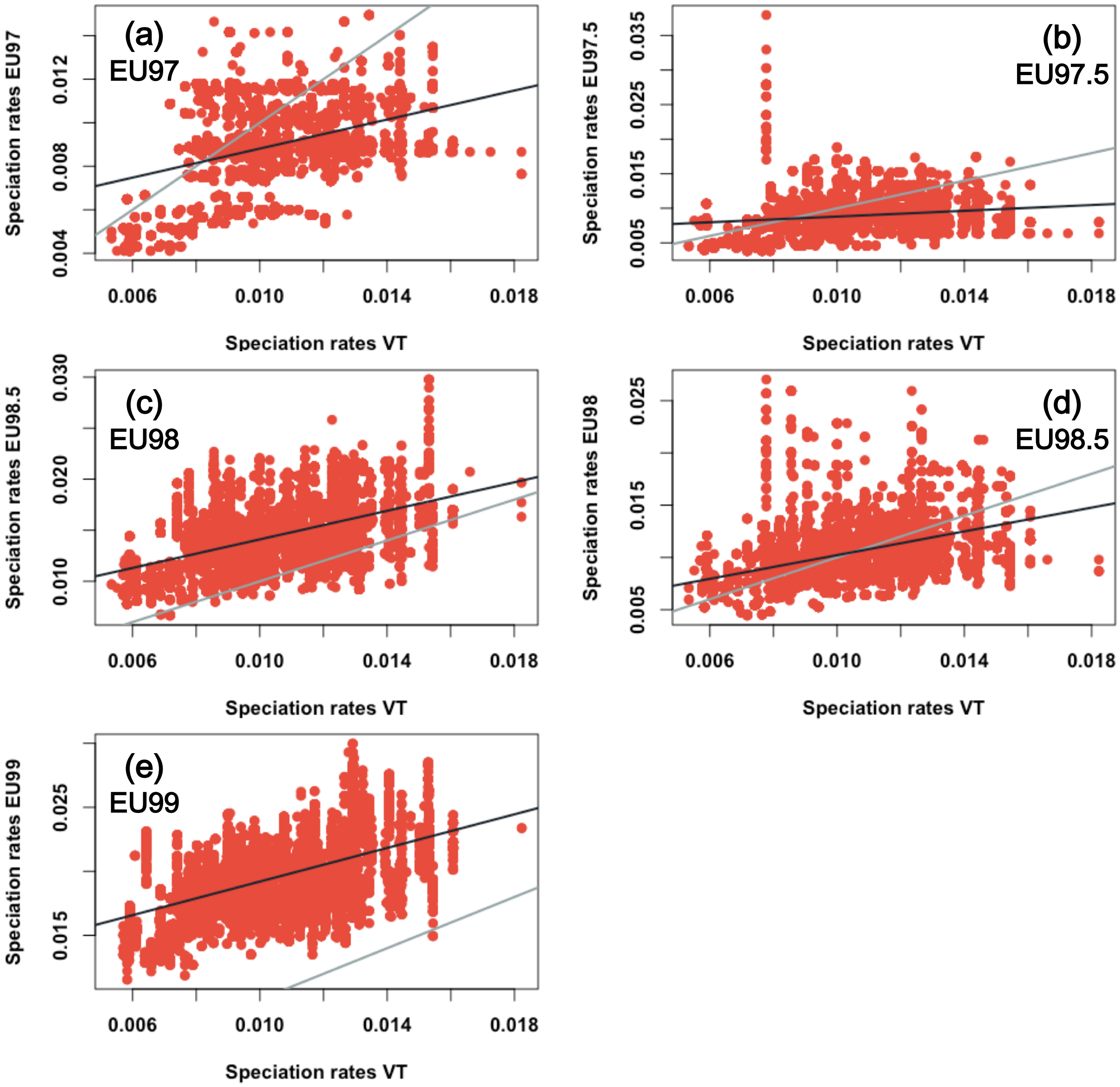
(a)
EU97
Speciation rates EU97
0.004
0.008
0.012
Speciation rates VT
(b)
EU97.5
Speciation rates EU97.5
0.005
0.015
0.025
0.035
Speciation rates VT
(c)
EU98
Speciation rates EU98.5
0.010
0.020
0.030
Speciation rates VT
(d)
EU98.5
Speciation rates EU98
0.005
0.015
0.025
Speciation rates VT
(e)
EU99
Speciation rates EU99
0.015
0.025
0.006
0.010
0.014
0.018
Speciation rates VT

**Supplementary Figure 10: The average speciation rates through time estimated by ClaDS show that Glomeromycotina experienced a decline in speciation rates toward the present after a period of high speciation rates.**

Average speciation rates (MAPS) through time estimated by ClaDS, for the different Glomeromycotina delineations, using the BDES estimated sampling fraction (Quince et al., 2008). The consensus trees are in dark grey, whereas orange lines represent the 100 independent replicate trees (we increased the number of replicate trees because a few EU99 trees show different trends).

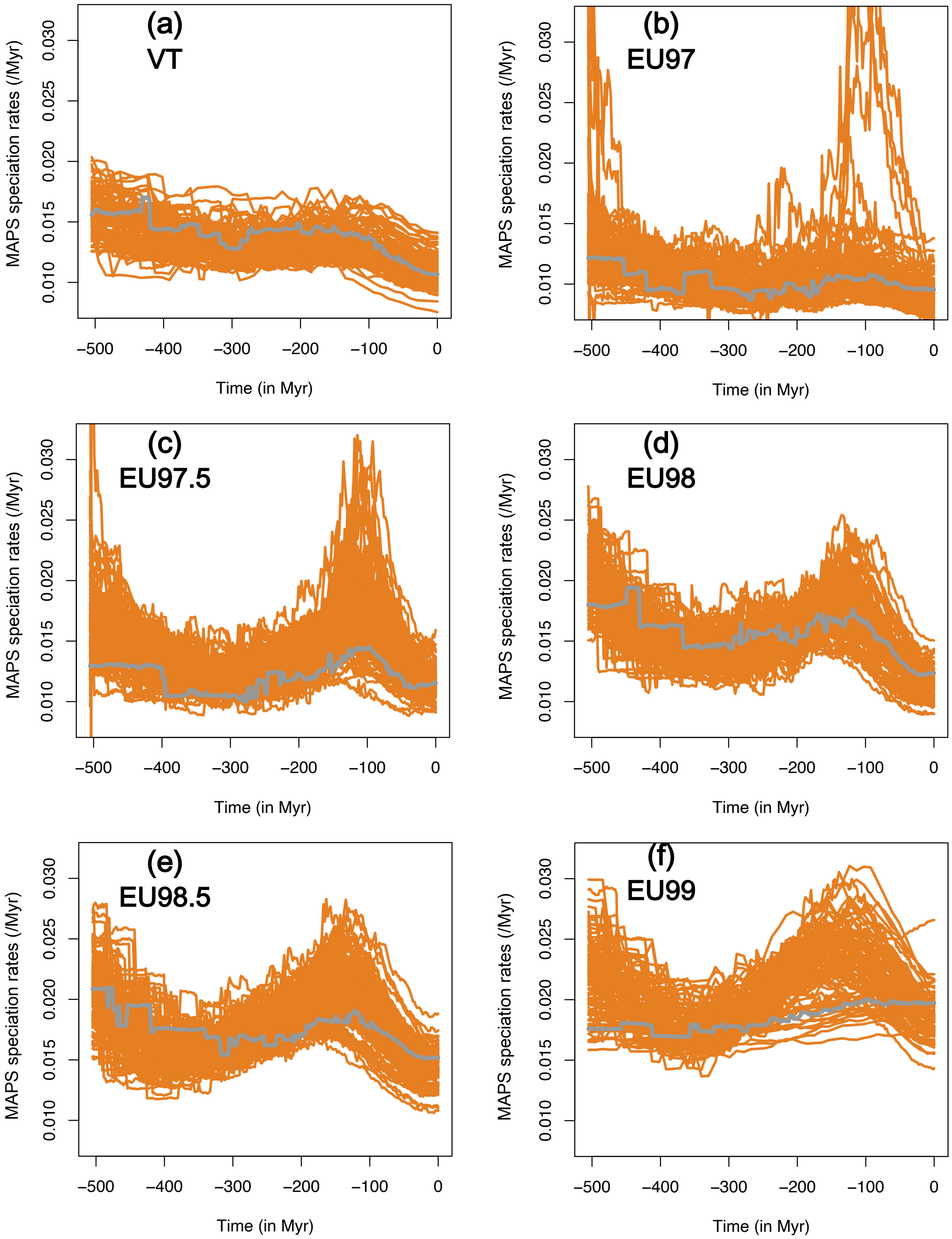
(a)
VT
(b)
EU97
(c)
EU97.5
(d)
EU98
(e)
EU98.5
(f)
EU99
MAPS speciation rates (/Myr)
0.010
0.015
0.020
0.025
0.030
Time (in Myr)
−500
−400
−300
−200
−100
0

**Supplementary Figure 11: The speciation rates through time estimated by CoMET show that Glomeromycotina experienced a decline in speciation rates toward the present after a period of high speciation rates.**

Mean Bayesian posterior of the speciation rates (/Myr) through time estimated by CoMET, for the different Glomeromycotina delineations, using the BDES estimated sampling fraction (Quince et al., 2008). Only the consensus trees are represented here, but the 12 independent replicate trees showed very similar trends. In each panel, the dark purple line represents the posterior mean and the purple polygon corresponds to the 95% credible interval.

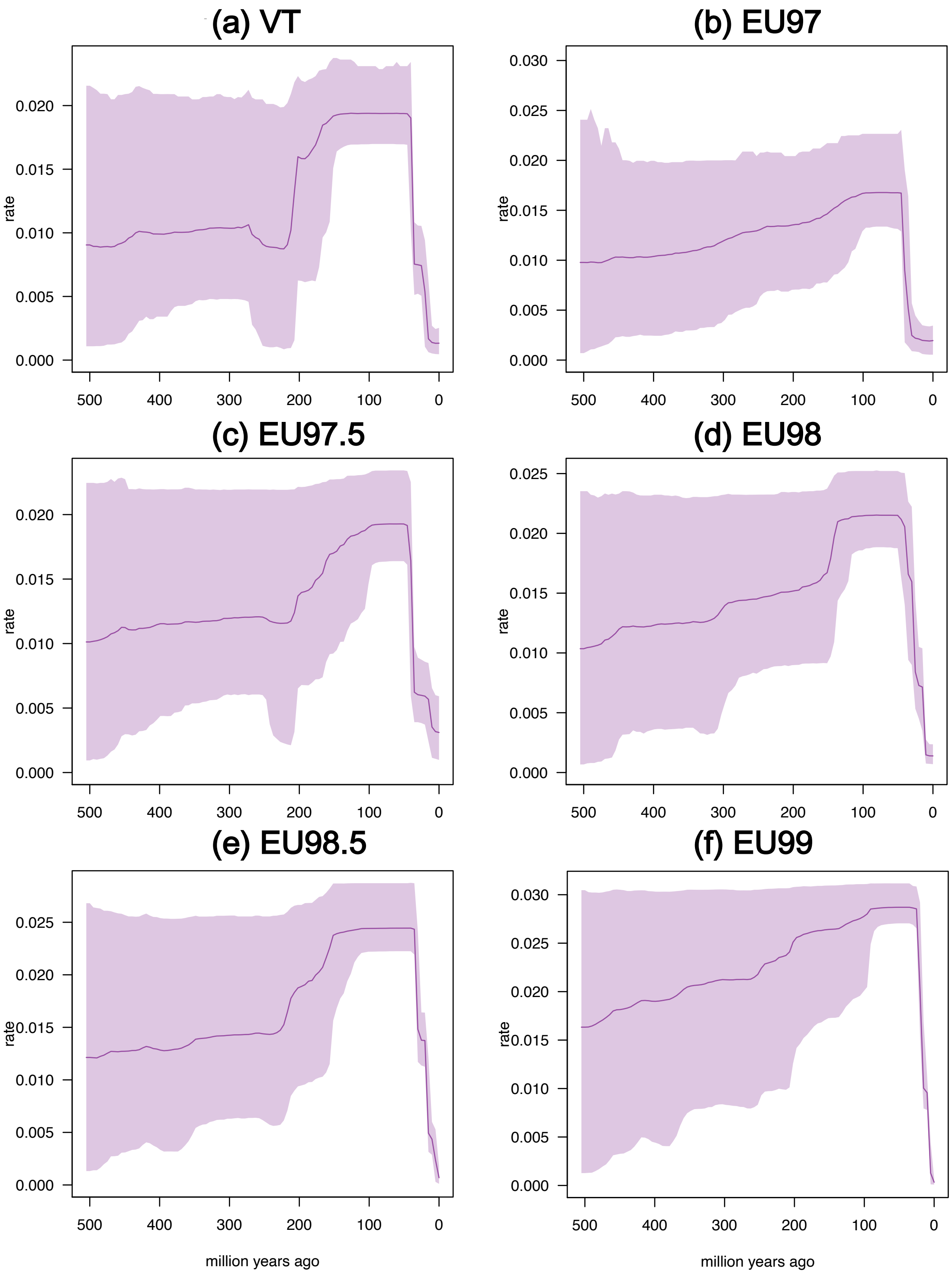
(a) VT
(b) EU97
(c) EU97.5
(d) EU98
(e) EU98.5
(f) EU99
rate
0.000
0.005
0.010
0.015
0.020
0.025
0.030
500
400
300
200
100
0
million years ago

**Supplementary Figure 12: Estimated hyperparameters of ClaDS2 runs, for the different Glomeromycotina delineations, using the BDES estimated sampling fraction.** At each speciation event, the two daughter lineages inherit new speciation rates sampled from a log-normal distribution with an expected value equals to log[*alpha* × λ] (where λ represents the parental speciation rate) and a standard deviation *sigma*. See Maliet et al., 2019 for more details about the hyperparameters:

(a): Estimated stochasticity (sigma) representing the variability in the rates at speciation events.

(b): Estimated turnover (epsilon) – constant ratio between extinction rates and speciation rates.

(c): Estimated alpha parameters.

(d): Estimated m parameters, m = alpha × exp(sigma$^2$/2) representing the general trend of the rate through time (m<1 means that there is an average decrease through time).

For each delineation, the boxplots represent the parameter values obtained for the consensus tree and the 12 independent replicate trees.

Boxplots indicate the median surrounded by the first and third quartiles, and whiskers extend to the extreme values but no further than 1.5 of the inter-quartile range.

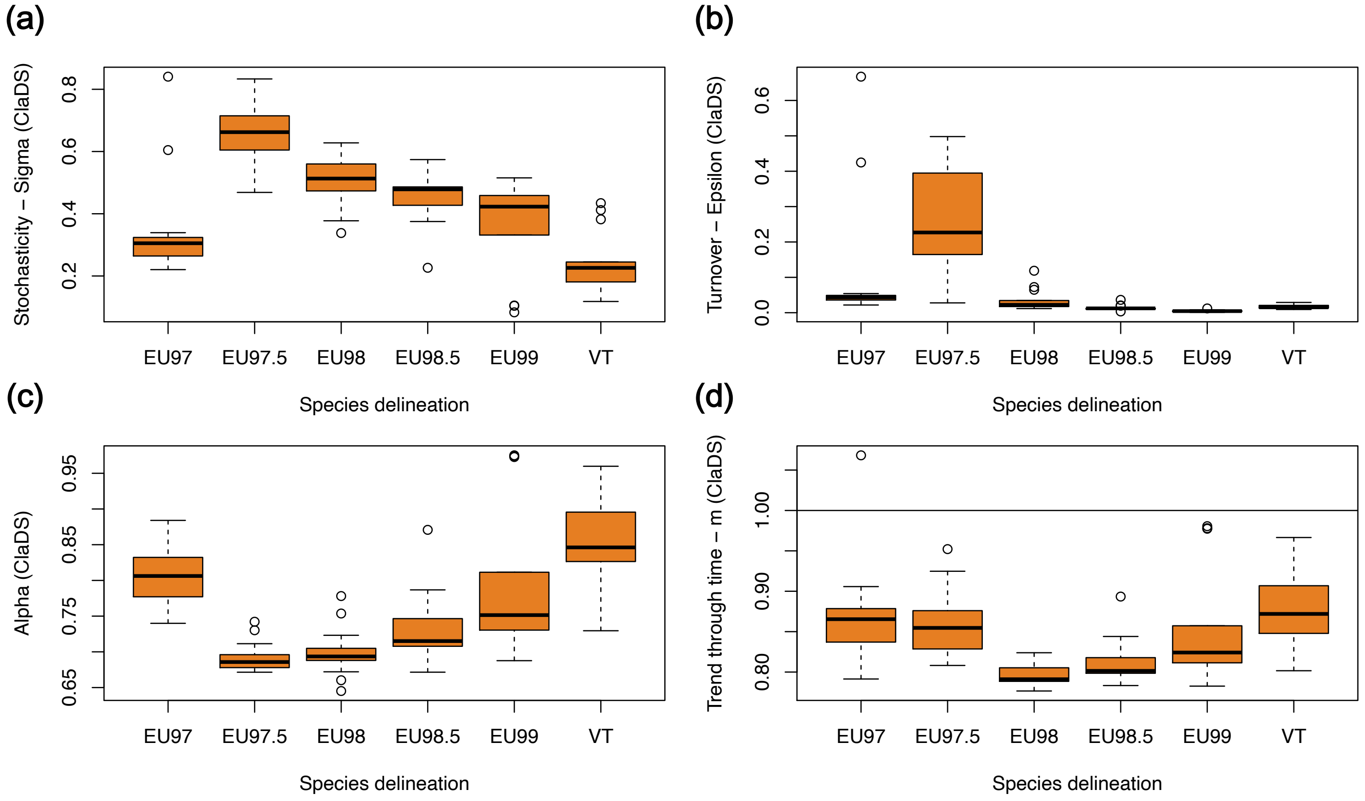

**Supplementary Figure 13: The speciation rates through time estimated by RPANDA show that Glomeromycotina experienced a decline in speciation rates toward the present.**

Speciation rates through time according to the best-supported time-dependent diversification model in RPANDA, for the different Glomeromycotina delineations and using the BDES estimated sampling fraction (Quince et al., 2008). The consensus trees are in dark grey, whereas orange lines represent the 12 independent replicate trees. None of these best-supported models include extinction.

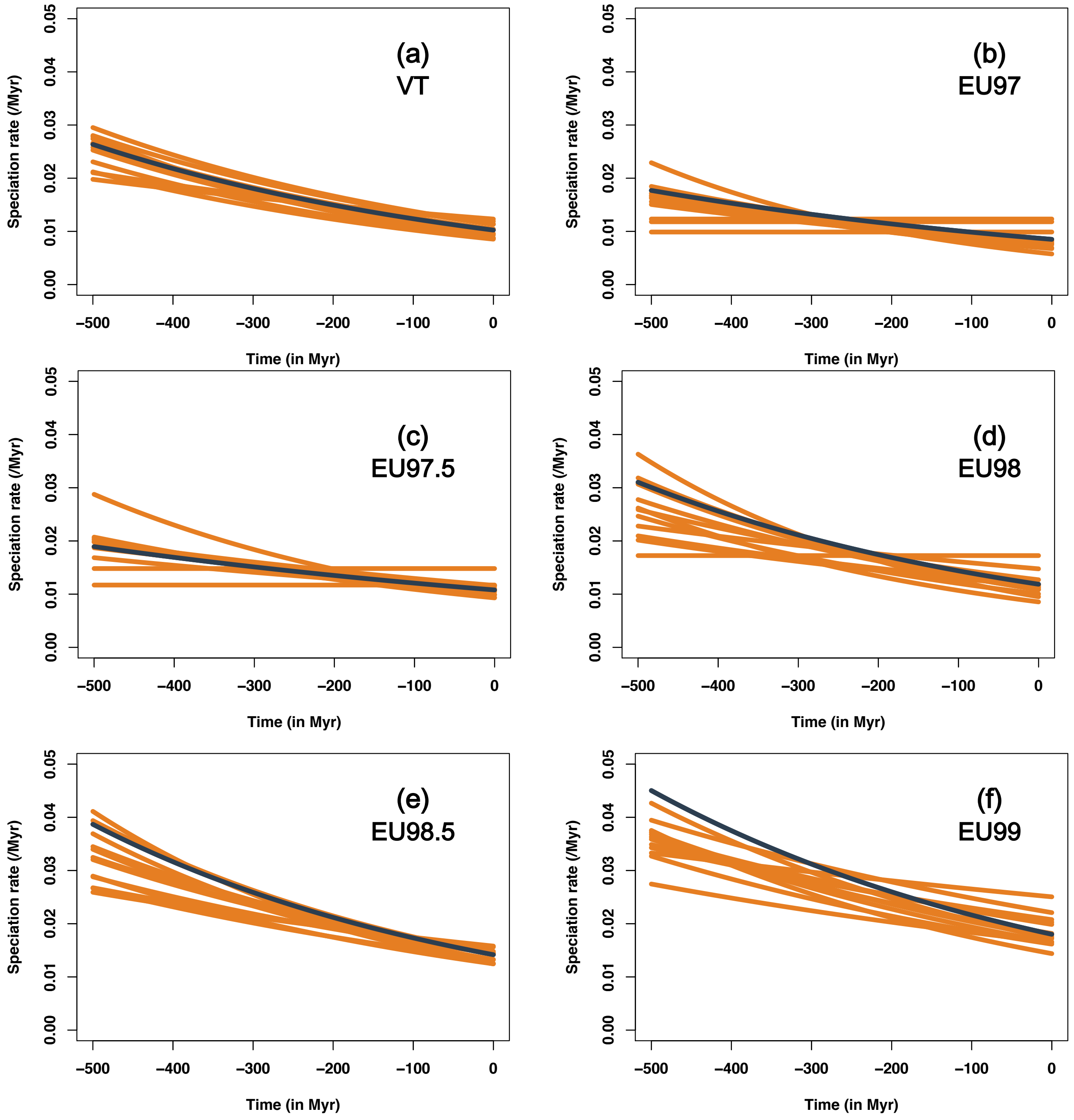
(a)
VT
(b)
EU97
(c)
EU97.5
(d)
EU98
(e)
EU98.5
(f)
EU99
Speciation rate (/Myr)
Time (in Myr)
0.00
0.01
0.02
0.03
0.04
0.05
−500
−400
−300
−200
−100
0

**Supplementary Figure 14: The speciation rates through time estimated by ClaDS also show that Glomeromycotina experienced a decline in speciation rates toward the present even when using sampling fractions <90%:**

Average speciation rates (MAPS) through time estimated by ClaDS, for the different Glomeromycotina delineations, for a range of sampling fraction: 50%, 60%, 70%, or 80% of sampled species. The consensus trees are in dark grey, whereas orange lines represent the 12 independent replicate trees.

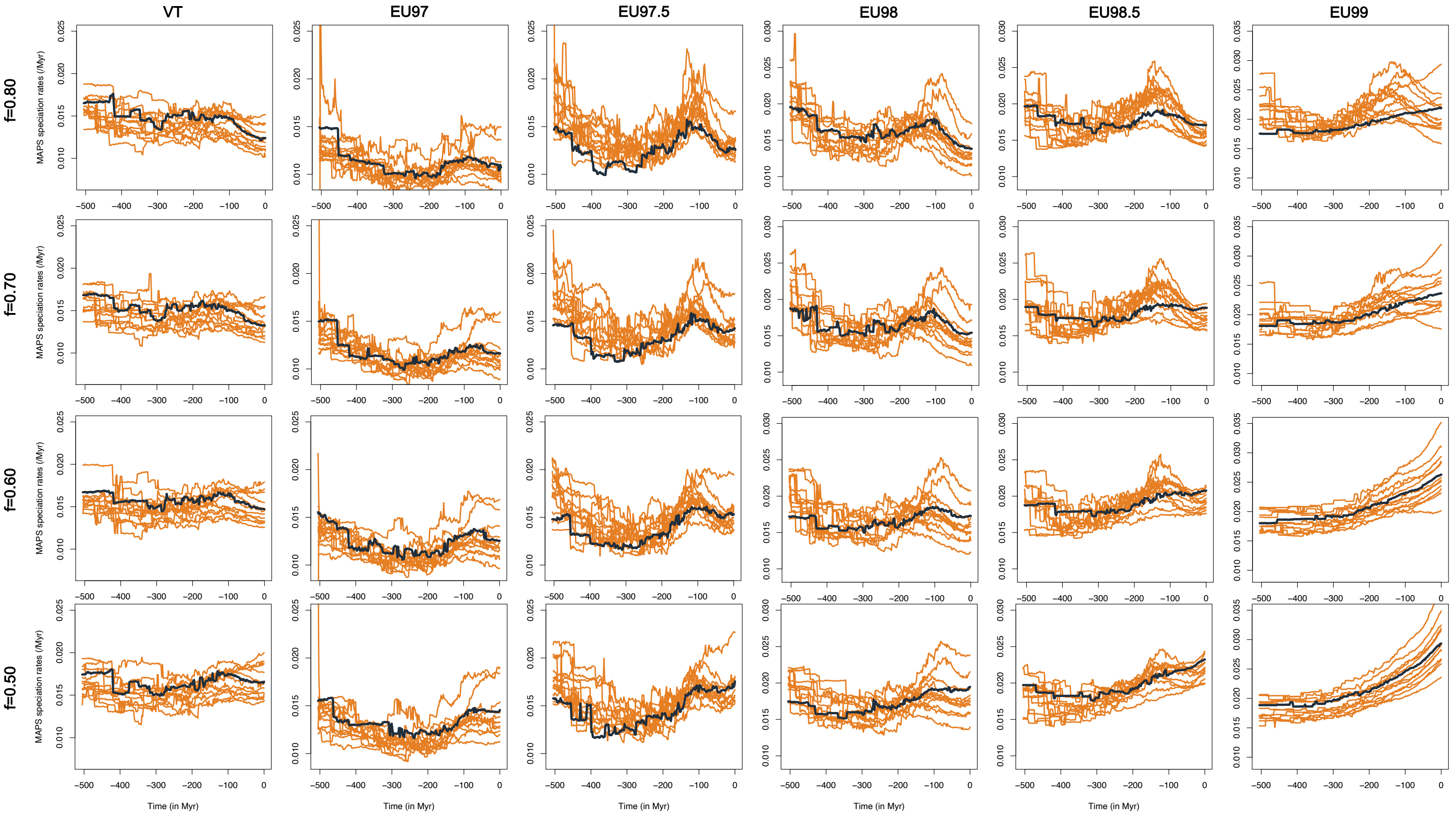

**Supplementary Figure 15: The speciation rates through time estimated by CoMET also show that Glomeromycotina experienced a decline in speciation rates toward the present even when using sampling fractions <90%:**

Mean Bayesian posterior of the speciation rates (/Myr) through time estimated by CoMET, for the different Glomeromycotina delineations, for a range of sampling fraction: 50%, 60%, 70%, or 80% of sampled species. Only the consensus trees are represented here, but the 12 independent replicate trees showed very similar trends. In each panel, the dark purple line represents the posterior mean and the purple polygon corresponds to the 95% credible interval.

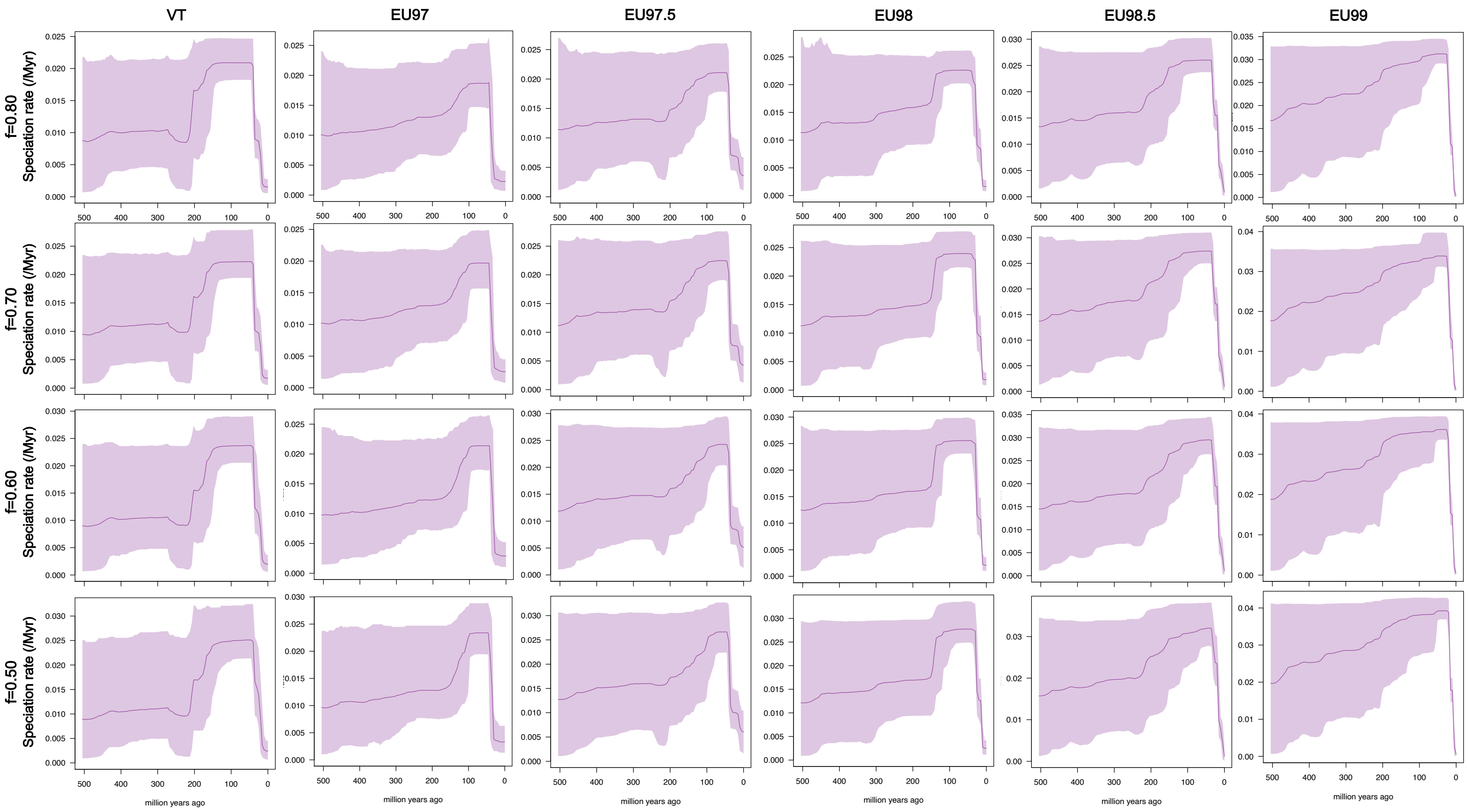

**Supplementary Figure 16: Speciation rates through time estimated by ClaDS using the 28S large sub-unit of the rRNA gene (LSU rRNA gene)**

Average speciation rates (MAPS) through time estimated by ClaDS, for the GMYC delineation, for a range of sampling fraction: 90%, 80%, 70%, 60%, or 50% of sampled species. The consensus trees are in dark grey, whereas orange lines represent the 12 independent replicate trees.

We observed a period of high speciation rates between 200 and 100 Myr ago that was followed by a decline in speciation rates toward the present, for assumed sampling fractions as low as 60%. For a sampling fraction of 50%, we nevertheless observed a plateau of the speciation rates in the last 50 Myr.

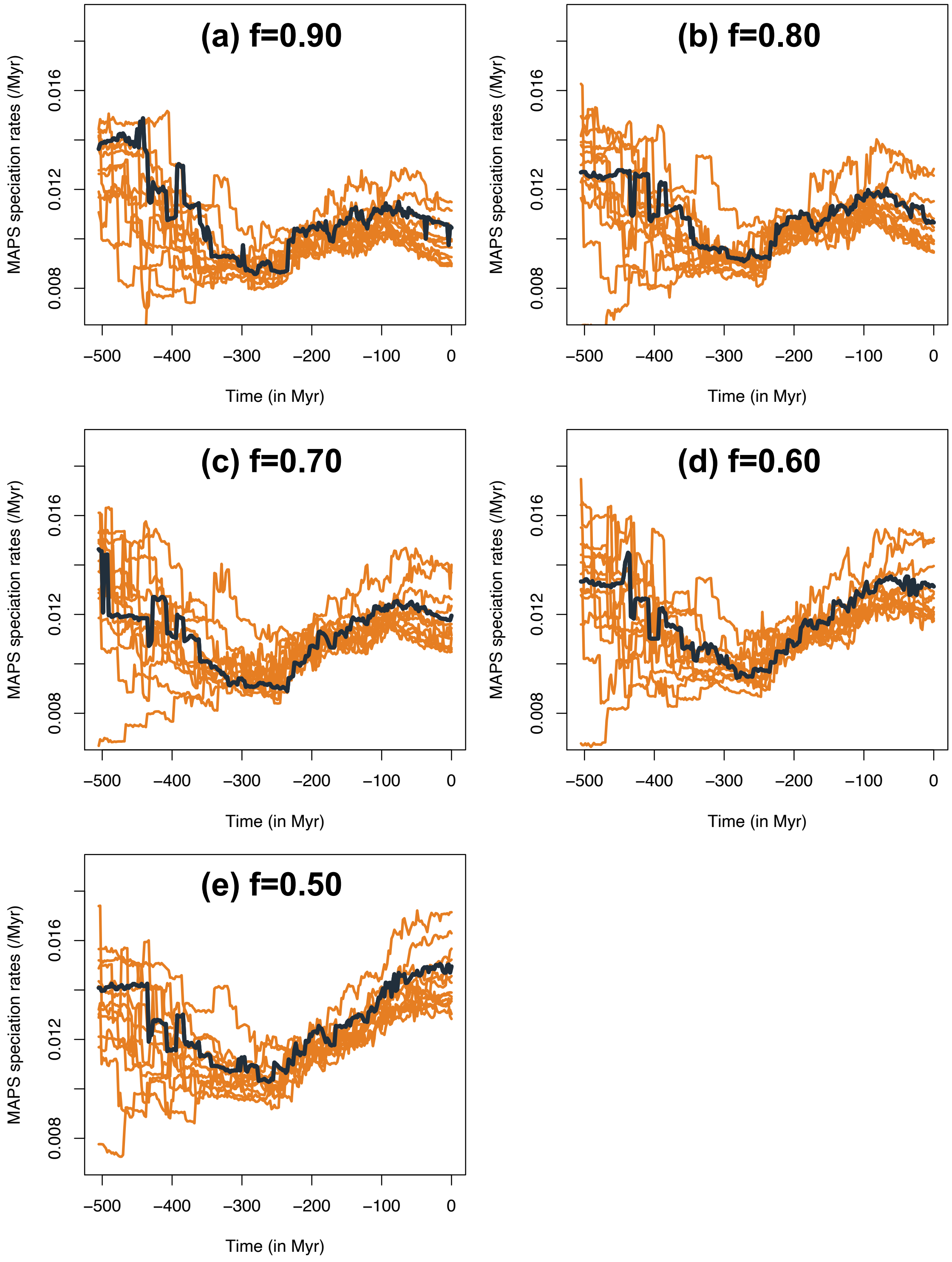
(a) f=0.90
(b) f=0.80
(c) f=0.70
(d) f=0.60
(e) f=0.50
MAPS speciation rates (/Myr)
0.008
0.012
0.016
−500
−400
−300
−200
−100
0
Time (in Myr)

**Supplementary Figure 17: Low support for extinction according to CoMET:** Bayesian posterior of the extinction rates (/Myr) through time estimated by CoMET, for the different Glomeromycotina delineations, using the BDES estimated sampling fraction (Quince et al., 2008). Only the consensus trees are represented here, but the 12 independent replicate trees showed very similar trends. In each panel, the red purple line represents the posterior mean and the red polygon corresponds to the 95% credible interval. Given that extinction rates of 0 are always contained within the Bayesian 95% credible intervals, there is no (or very few) support for extinction in Glomeromycotina.

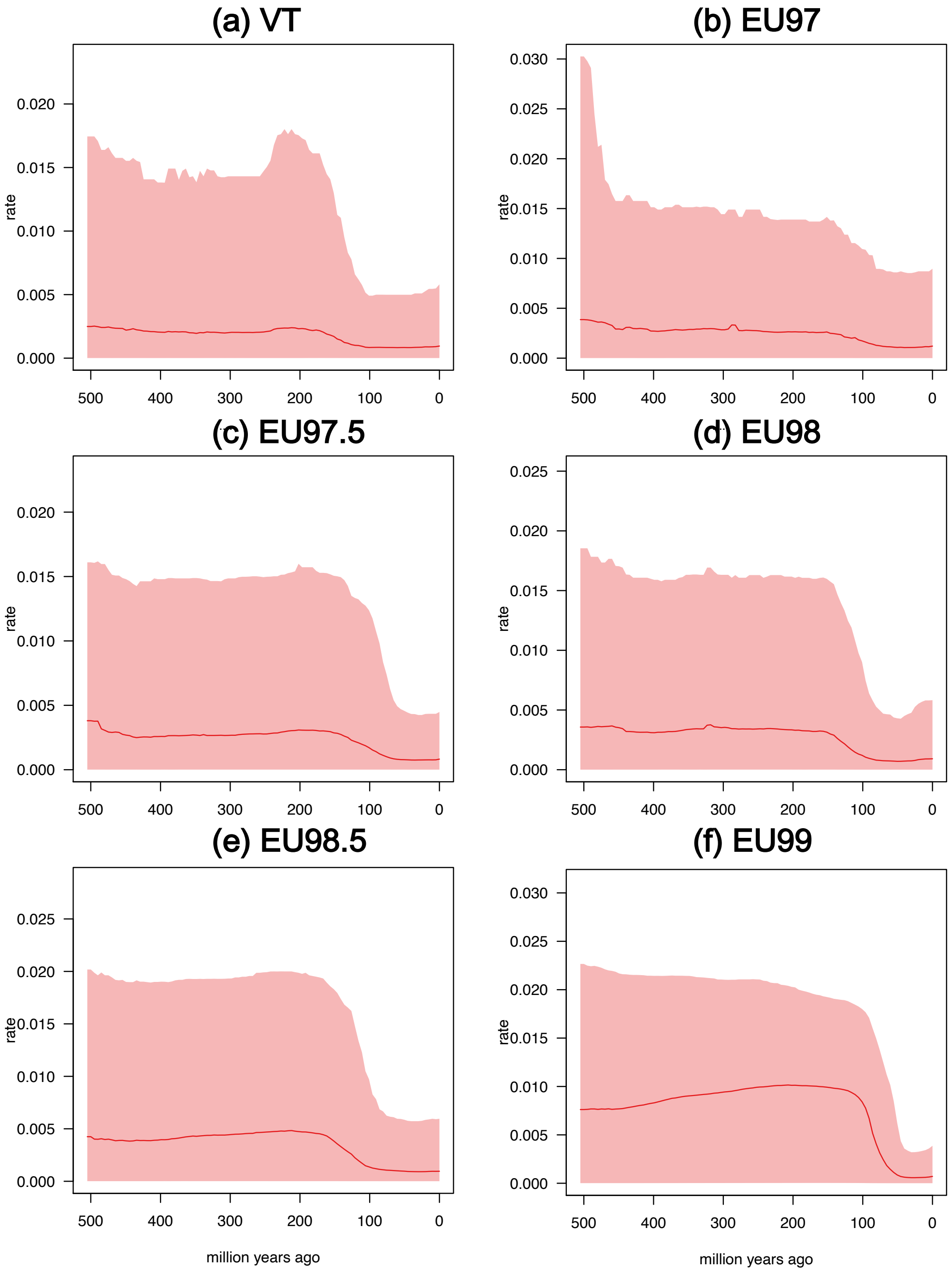

**Supplementary Figure 18: Inferred diversification rates decline faster in time-dependent models with fixed extinction rates.**

Each plot represents the speciation rates through time estimated by the exponential-time-dependent speciation rate model in RPANDA with a fixed extinction rate (from 0 to 0.03 extinction events/Myr – represented in different colors). Analyses were replicated for the different Glomeromycotina delineations and using the BDES estimated sampling fraction (Quince et al., 2008), but only the results of the consensus trees are represented here for simplicity (the replicates trees show the same trends).

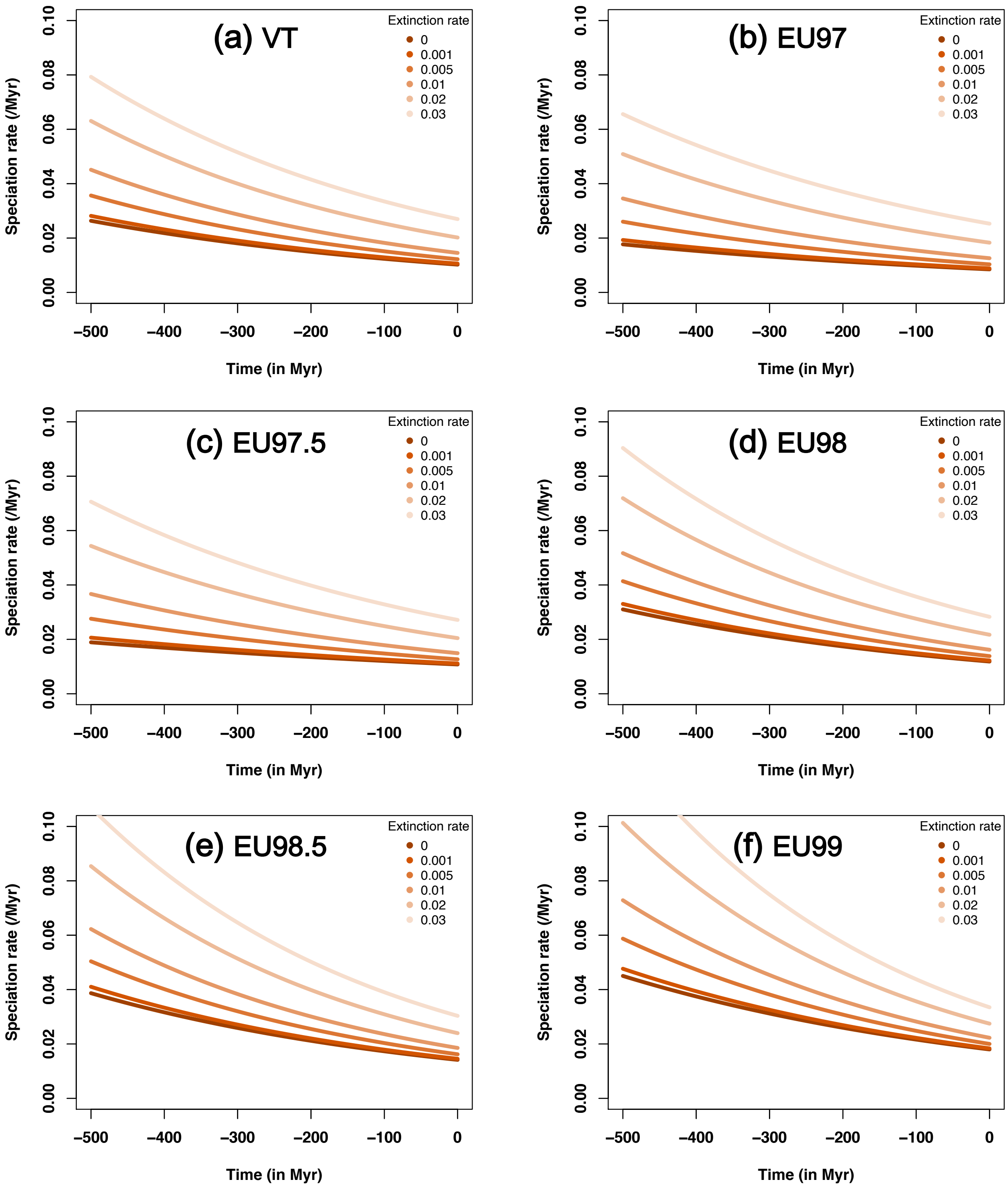
(a) VT
(b) EU97
(c) EU97.5
(d) EU98
(e) EU98.5
(f) EU99
Extinction rate
0
0.001
0.005
0.01
0.02
0.03
Speciation rate (/Myr)
0.00
0.02
0.04
0.06
0.08
0.10
Time (in Myr)
−500
−400
−300
−200
−100
0

**Supplementary Figure 19: Inferred diversification rates decline faster in congruent models with fixed extinction rates.**

Each plot represents speciation rates through time congruent to the best-fit time-dependent diversification model in RPANDA and with fixed extinction rate (from 0 to 0.03 extinction events/Myr – represented in different colors). Analyses were replicated for the different Glomeromycotina delineations and using the BDES estimated sampling fraction (Quince et al., 2008), but only the results of the consensus trees are represented here for simplicity (the replicates trees show the same trends).

Note that by definition all the congruent models have the same likelihood, but given that they include an additional extinction parameter (the best-fit time-dependent diversification model in RPANDA has no extinction), they would not be selected in a model comparison.

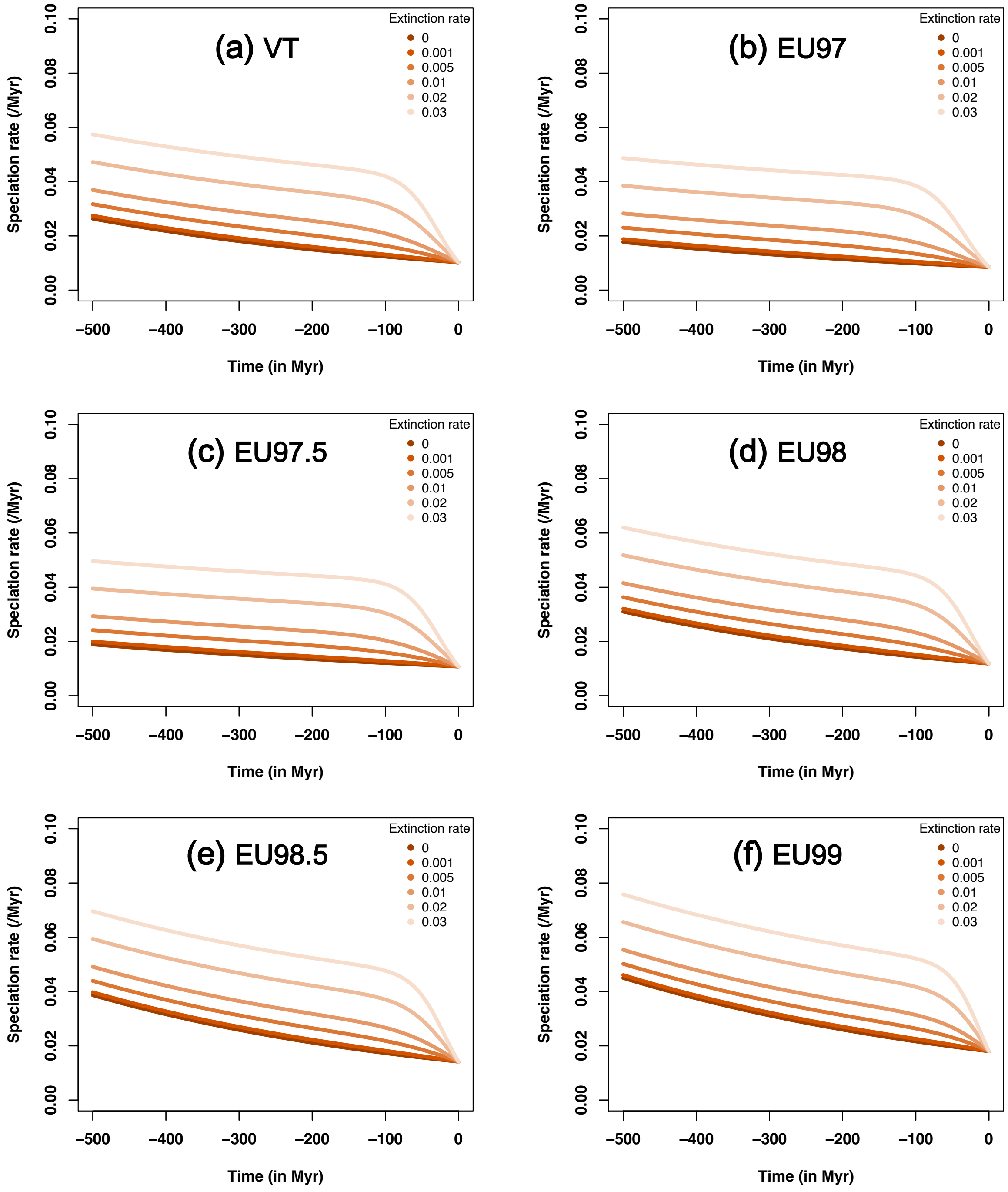

(a) VT
(b) EU97
(c) EU97.5
(d) EU98
(e) EU98.5
(f) EU99
Extinction rate
0
0.001
0.005
0.01
0.02
0.03
Speciation rate (/Myr)
Time (in Myr)
0.00
0.02
0.04
0.06
0.08
0.10
−500
−400
−300
−200
−100
0

**Supplementary Figure 20: Variation of the environmental variables through time tested with RPANDA:**

(a) average temperature (data up to 520 million years (Myr) ago), (b) $pCO_2$ (data up to 430 Myr ago), and (c-d) land plant diversity (data up to 430 Myr ago). Land plant diversity were estimated using the Paleobiology database and the shareholder quorum subsampling method (Supplementary Methods 6). Panel (c) represents land plant diversities estimated using different quorum values, from 0.2 (dark green) to 0.7 (light green), and panel (d) focuses the quorum value of 0.5.

For each environmental variable, we smoothed the curves using cubic splines: the degree of freedom of each smoothing is 33 for the temperature (a), 128 for pCO2 (b), and 25 for the land plant diversity (d). These smoothing values were used for fitting environment-dependent diversification models in RPANDA.

Temperature and $CO_2$ appeared to be importantly correlated.

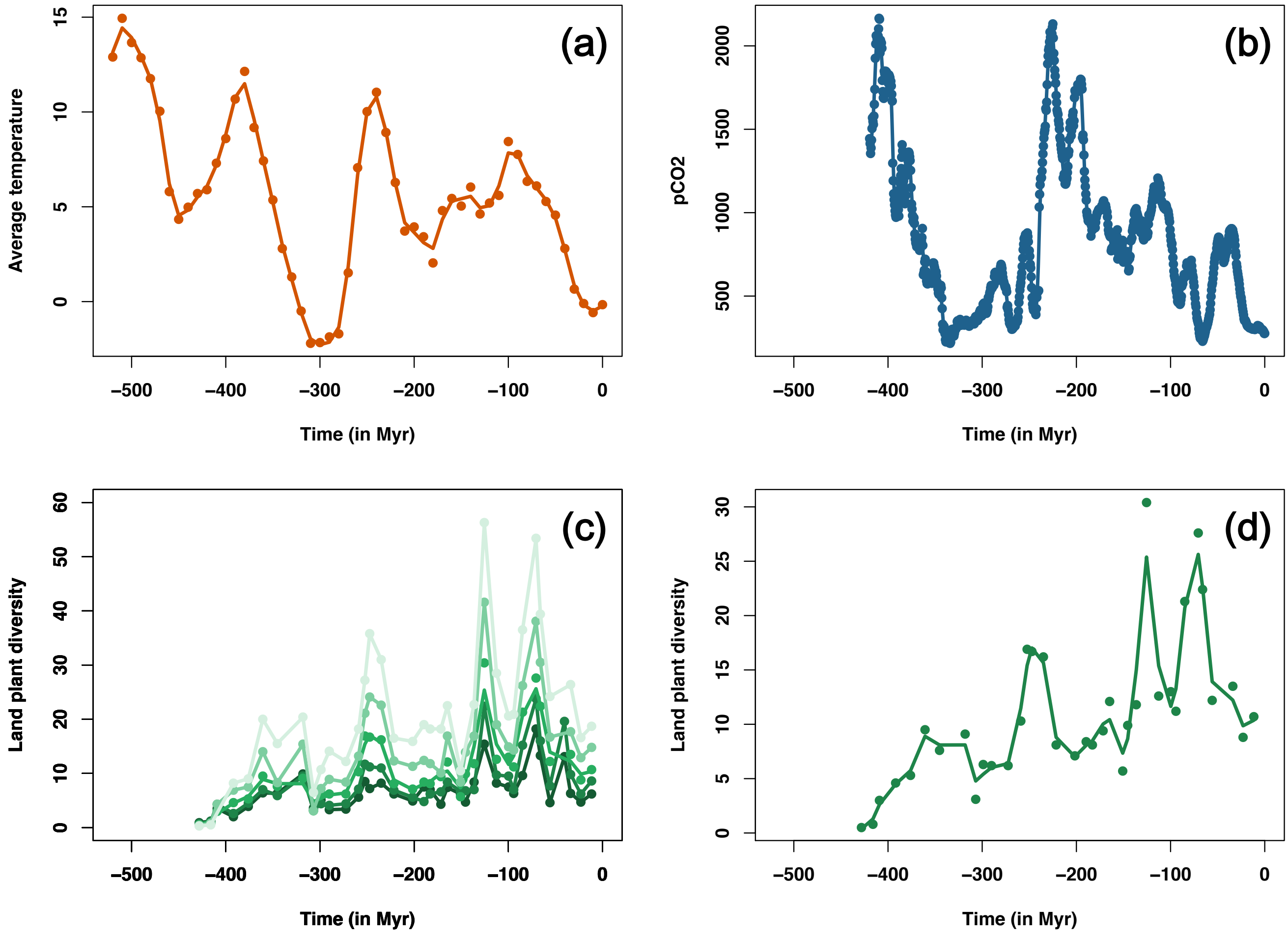

**Supplementary Figure 21: Speciation rates through time according to the temperature-dependent diversification model in RPANDA**, for the different Glomeromycotina delineations and using the BDES estimated sampling fraction (Quince et al., 2008). The consensus trees are in dark grey, whereas orange lines represent the 12 independent replicate trees.

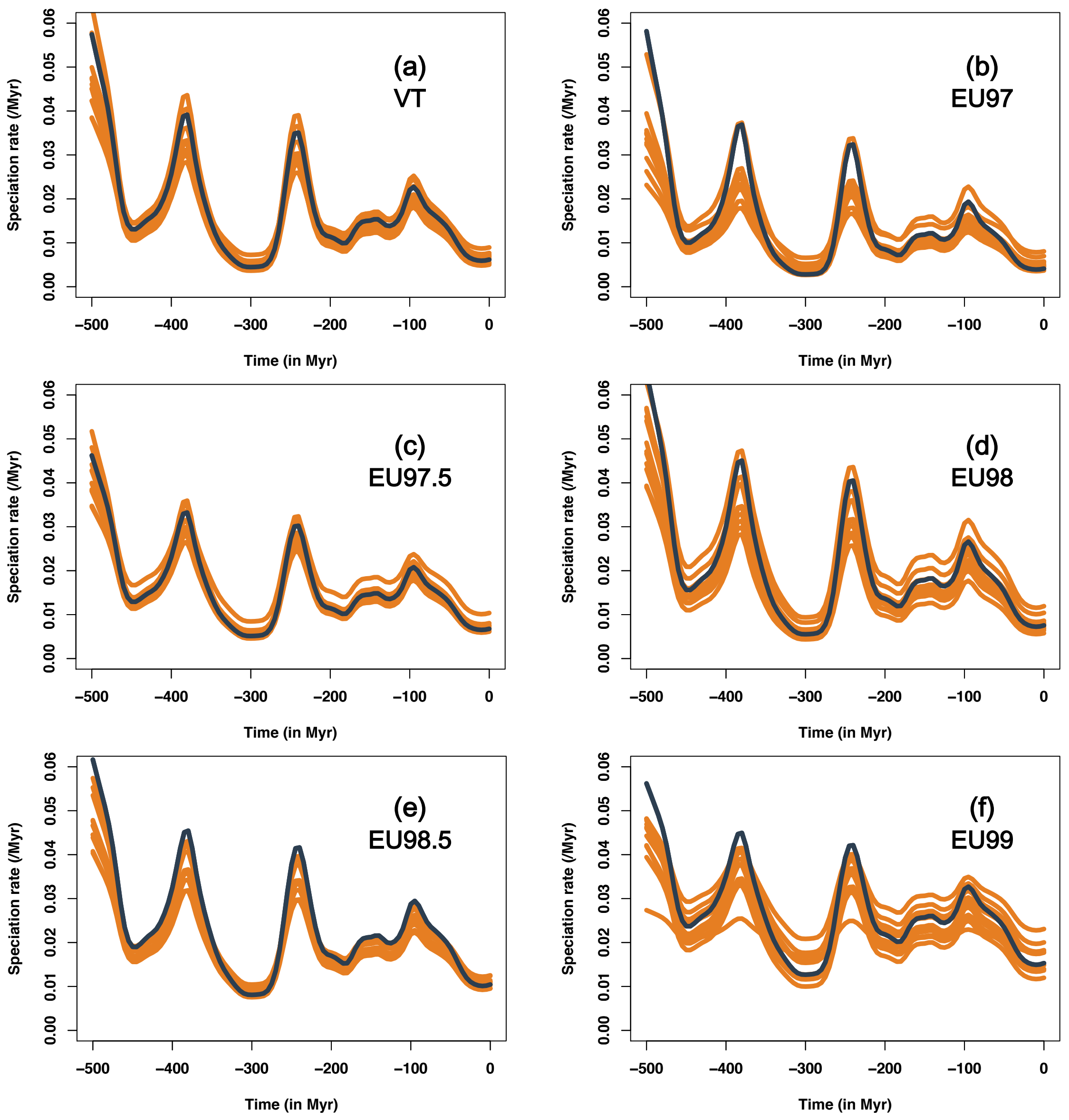

**Supplementary Figure 22: Temperature-dependent models are significantly supported in Glomeromycotina:**

Best-supported models between the time-dependent and temperature-dependent diversification models in RPANDA, for the different Glomeromycotina delineations and using the BDES estimated sampling fraction (Quince et al., 2008).

(a) AICc difference between the best-supported time-dependent model and the temperature-dependent model in RPANDA. An AICc difference greater than 2 indicates that there is a significant support for the temperature-dependent model.

(b) Parameter estimation of the temperature-dependent model (speciation rate ~ exp(parameter * temperature) ). A positive parameter value indicates a positive effect of temperature on speciation rates.

For each delineation, the boxplots represent the results obtained for the consensus tree and the 12 independent replicate trees.

Boxplots indicate the median surrounded by the first and third quartiles, and whiskers extend to the extreme values but no further than 1.5 of the inter-quartile range.

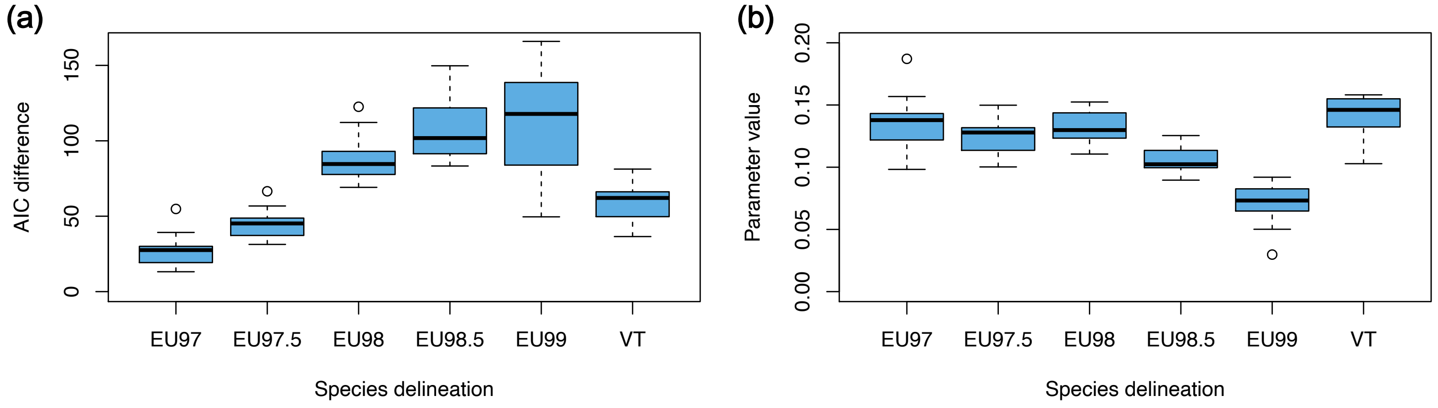

**Supplementary Figure 23: Temperature-dependent models are significantly supported in Glomeromycotina, even when using sampling fractions <90%:**

Best-supported models between the time-dependent and temperature-dependent diversification models in RPANDA, for the different Glomeromycotina delineations and for a range of sampling fraction: 50% (a, b), 60% (c, d), 70% (e, f), or 80% (g, h) of sampled species.

**(a, c, e, g)** AICc difference between the best-supported time-dependent model and the temperature-dependent model in RPANDA. An AICc difference greater than 2 indicates that there is a significant support for the temperature-dependent model.

**(b, d, f, h)** Parameter estimation of the temperature-dependent model (speciation rate ~ exp(parameter * temperature) ). A positive parameter value indicates a positive effect of temperature on speciation rates.

For each delineation, the boxplots represent the results obtained for the consensus tree and the 12 independent replicate trees.

Boxplots indicate the median surrounded by the first and third quartiles, and whiskers extend to the extreme values but no further than 1.5 of the inter-quartile range.

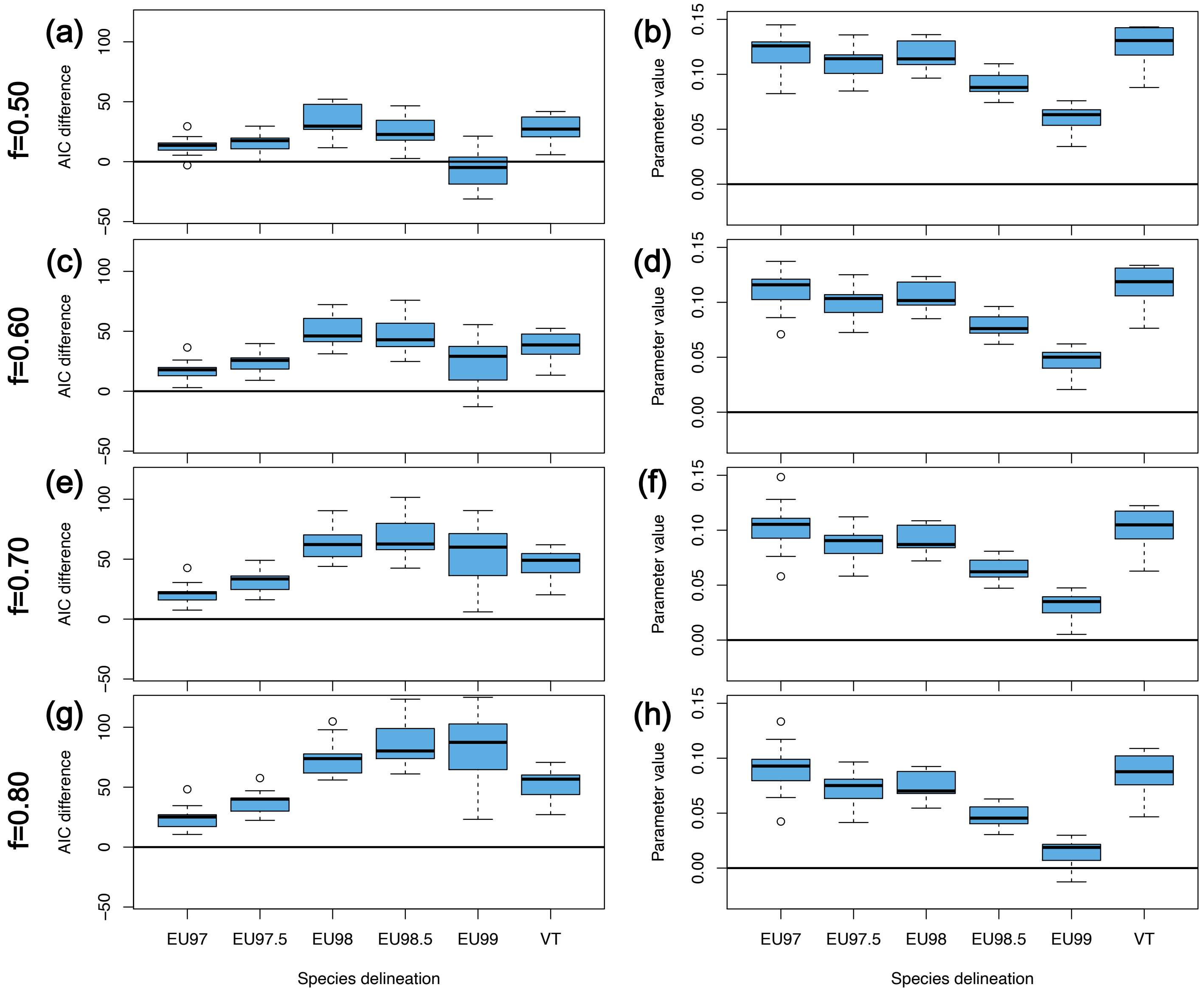

f=0.50
f=0.60
f=0.70
f=0.80
(a)
(b)
(c)
(d)
(e)
(f)
(g)
(h)
AIC difference
Parameter value
EU97
EU97.5
EU98
EU98.5
EU99
VT
Species delineation

**Supplementary Figure 24: Effect of the Glomeromycotina crown age on the RPANDA models.**

For the different Glomeromycotina delineations, analyses were reproduced on the Glomeromycotina phylogenetic trees with two extreme values of crown age: 530 Myr old (a-b) and 437 Myr old (c-d) (Lutzoni et al., 2018). We used the BDES estimated sampling fraction (Quince et al., 2008).

**(a-c)** AICc difference between the best-supported time-dependent model and the temperature-dependent model in RPANDA. An AICc difference greater than 2 indicates that there is a significant support for the temperature-dependent model.

**(b-d)** Parameter estimation of the temperature-dependent model (speciation rate ~ exp(parameter * temperature) ). A positive parameter value indicates a positive effect of temperature on speciation rates.

For each delineation, the boxplots represent the results obtained for the consensus tree and the 12 independent replicate trees.

Boxplots indicate the median surrounded by the first and third quartiles, and whiskers extend to the extreme values but no further than 1.5 of the inter-quartile range.

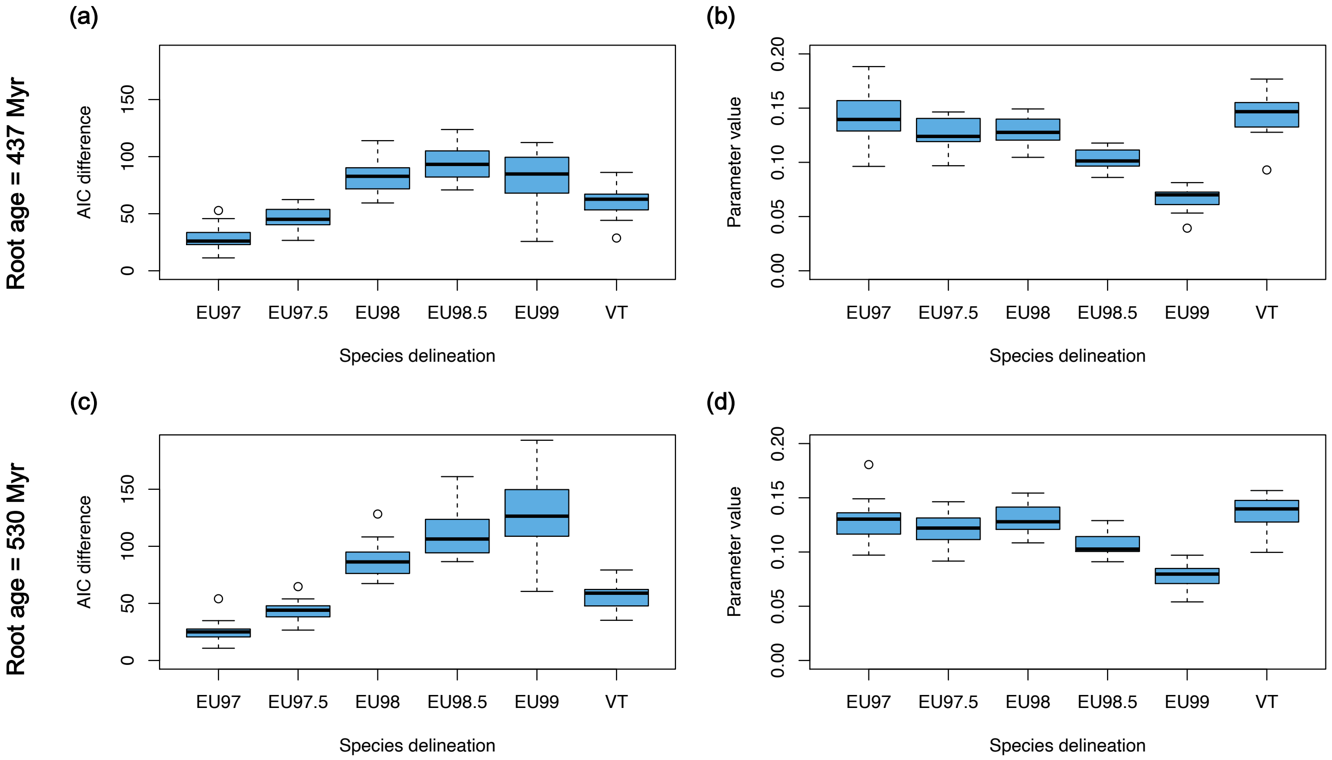

**Supplementary Figure 25: Temperature-dependent models are not artifactually supported when time-dependency is simulated.**

**(a-f)** For each Glomeromycotina delineation, the boxplot represents the size distribution (number of tips) of the trees simulated under the best-supported time-dependent model inferred using RPANDA on the consensus tree. The horizontal orange lines represented the estimated total number of species per delineation (the observed number of species divided by the estimated BDES sampling fraction). VT phylogenies simulated under the best-fit time-dependent model had sizes centered around the size of the actual Glomeromycotina phylogeny, whereas the simulated EU phylogenies had on average less species than the actual EU phylogenies.

**(g)** For every simulated trees of each Glomeromycotina delineation, the difference of AICc ($\Delta$) was computed between the best-supported time-dependent model and the temperature-dependent model, and the bar chart indicates the number of simulated trees that present either a significant effect of temperature ($\Delta>2$), or no significant effect of temperature ($\Delta<2$). For all delineations and most of the simulations of time-dependency, temperature-dependent models were not supported.

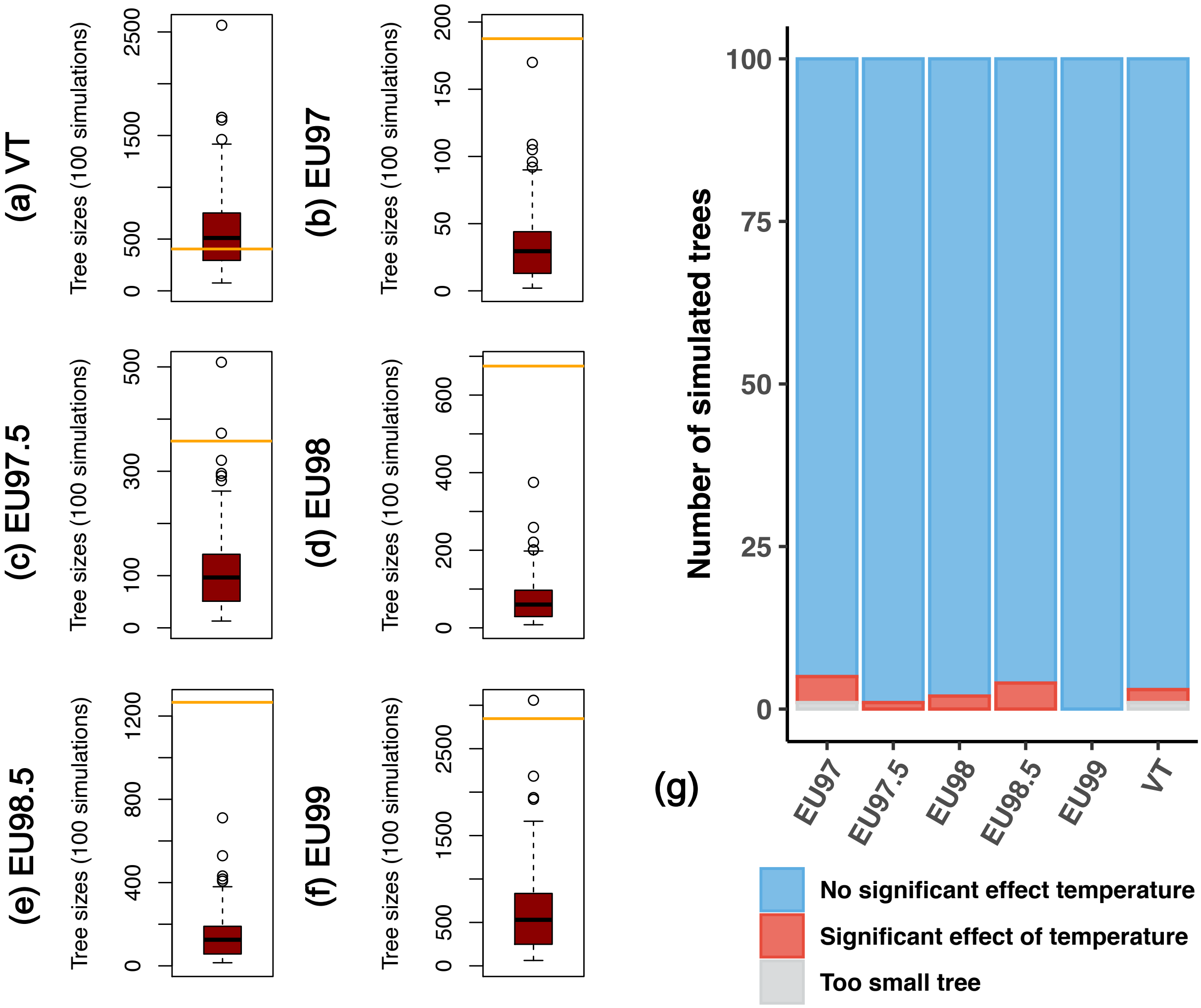
(a) VT
(b) EU97
(c) EU97.5
(d) EU98
(e) EU98.5
(f) EU99
Tree sizes (100 simulations)
(g)
Number of simulated trees
EU97
EU97.5
EU98
EU98.5
EU99
VT
No significant effect temperature
Significant effect of temperature
Too small tree

**Supplementary Figure 26: Temperature-dependent models are not supported because of a global temporal trend in temperature variation.**

Analyses were replicated for the different Glomeromycotina delineations (b-g) and on the consensus and replicate trees, using the BDES estimated sampling fraction.

(a) Temperature curves in the last 520 Myr obtained with different smoothing: the degrees of freedom (df) vary from 33 (original smoothing) to 2 (only a general trend of decrease).

(b-g) The barplots represent in blue the proportion of Glomeromycotina trees (the consensus tree and the 12 replicate trees) which present a fit with the smoothed temperature-dependent model that is as good as the original temperature-dependent model (AICc difference lower than 2). If a smoothed temperature-dependent model presents an AICc difference greater than 2 with the original temperature-dependent model, we considered that there is no more support for the smoothed temperature-dependent model (represented in grey).

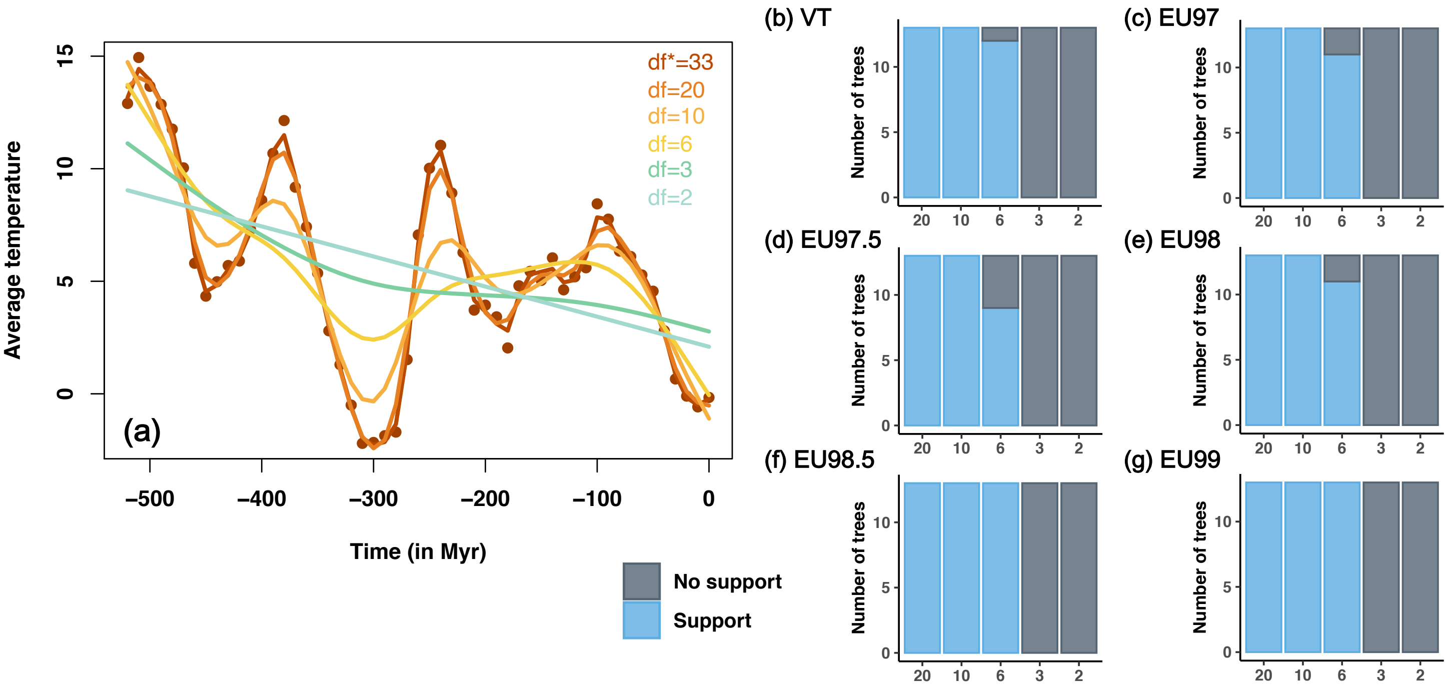

**Supplementary Figure 27: The support for temperature-dependent models is not linked to the heterogeneity of rates across lineages.**

**(a-f)** For each Glomeromycotina delineation, the boxplot represents the size distribution (number of tips) of the trees simulated with hyperparameters inferred using ClaDS2 on the consensus tree. The horizontal orange lines represent the estimated total number of Glomeromycotina species per delineation (the observed number of species divided by the estimated BDES sampling fraction). Phylogenies simulated under ClaDS hyperparameters had sizes centered around the size of the actual Glomeromycotina phylogenies (expect for EU97.5 and EU98).

**(g)** For every simulated trees of each Glomeromycotina delineation, the difference of AICc (Δ) was computed between the best-supported time-dependent model and the temperature-dependent model, and the bar chart indicates the number of simulated trees that present either a significant effect of temperature (Δ>2), or no significant effect of temperature (Δ<2). Temperature-dependent models were not supported, suggesting that the support for temperature-dependent models found on Glomeromycotina phylogenies is not linked to unaccounted for heterogeneity in rates across lineages.

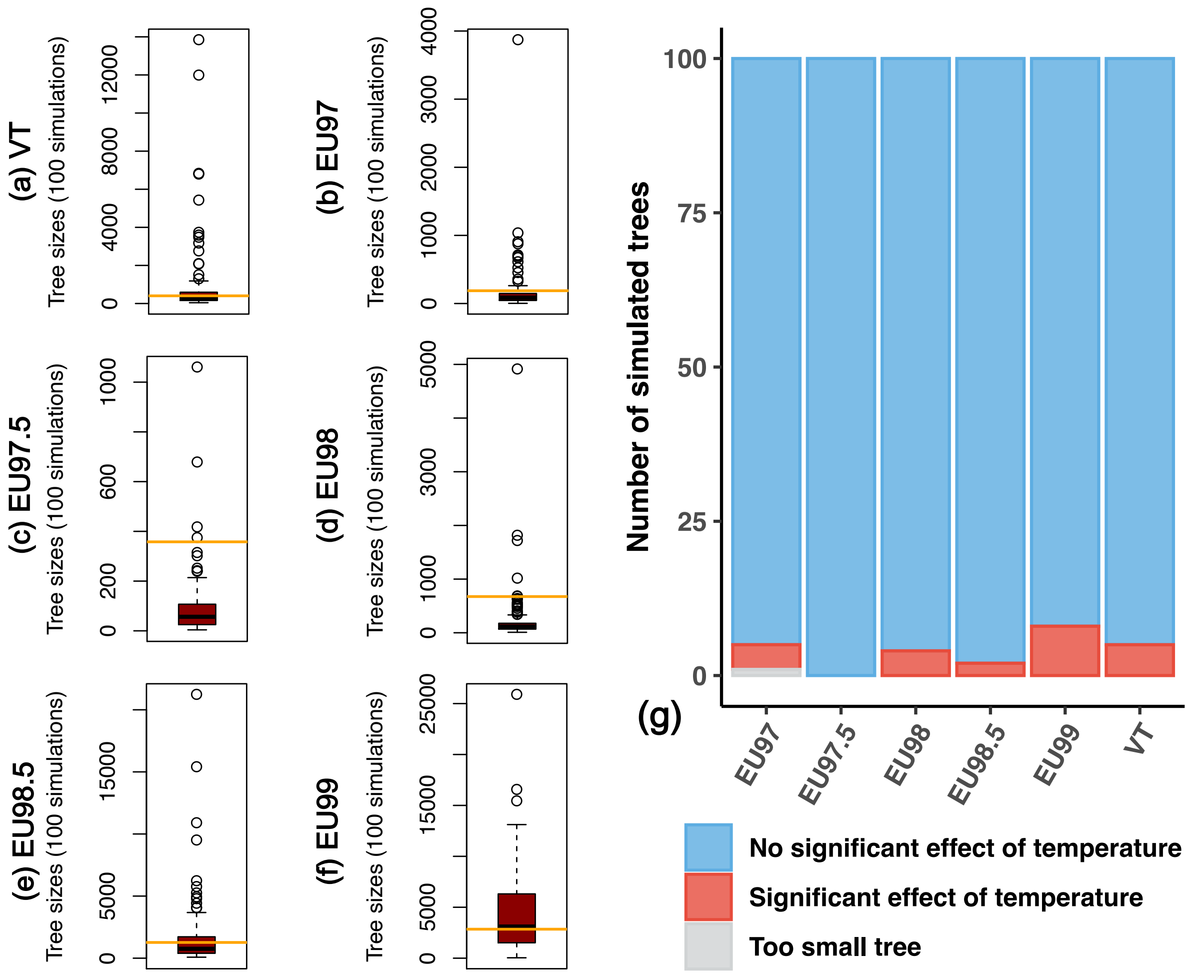
(a) VT
Tree sizes (100 simulations)
(b) EU97
Tree sizes (100 simulations)
(c) EU97.5
Tree sizes (100 simulations)
(d) EU98
Tree sizes (100 simulations)
(e) EU98.5
Tree sizes (100 simulations)
(f) EU99
Tree sizes (100 simulations)
(g)
Number of simulated trees
EU97
EU97.5
EU98
EU98.5
EU99
VT
No significant effect of temperature
Significant effect of temperature
Too small tree

**Supplementary Figure 28: The different Glomeromycotina sub-clades present significant support for temperature-dependence diversification, but also for dependences with $CO_2$ and land plants:**

Best-supported models between the environment-dependent (temperature, $CO_2$, and land plant fossil diversity) and time-dependent diversification model in RPANDA in 200 million years (Myr)-old sub-trees (a-b) or 400 Myr-old sub-trees (c-d) for which land plant fossil diversity, $CO_2$, and temperature can be tested.

The y-axis indicates the number of sliced trees with more than 50 tips at the time for which each model is best selected. Analyses were replicated for the different Glomeromycotina delineations and on the consensus (represented on the right of each delineation) and replicate trees (the results of the 100 trees are represented on the left of each delineation), using the BDES estimated sampling fraction (Quince et al., 2008).

**(a)** For each sliced tree at 200 Myr, the barplot indicates the model with the smallest AICc.

**(b)** For each sliced tree at 200 Myr, the barplot indicates the model with the smallest AICc and an AICc difference with all others models larger than 2. If the difference is smaller than 2, it is considered as non-significant.

**(c)** For each sliced tree at 400 Myr, the barplot indicates the model with the smallest AICc.

**(d)** For each sliced tree at 400 Myr, the barplot indicates the model with the smallest AICc and an AICc difference with all others models larger than 2. If the difference is smaller than 2, it is considered as non-significant.

Only one sliced tree in two EU99 replicates supports a model with extinction (denoted as "Exponential + extinction") and a few trees support the effect of land plant fossil diversity.

These analyses were replicated to take into account the uncertainty in the crown root age of Glomeromycotina: they were replicated on the phylogeny with a crown age of 505 Myr (I), as well as using the youngest (437 Myr - II) and oldest (530 Myr - III) crown age estimates from Lutzoni *et al.* (2018).

#### (I) Glomeromycotina crown age of 505 Myr:

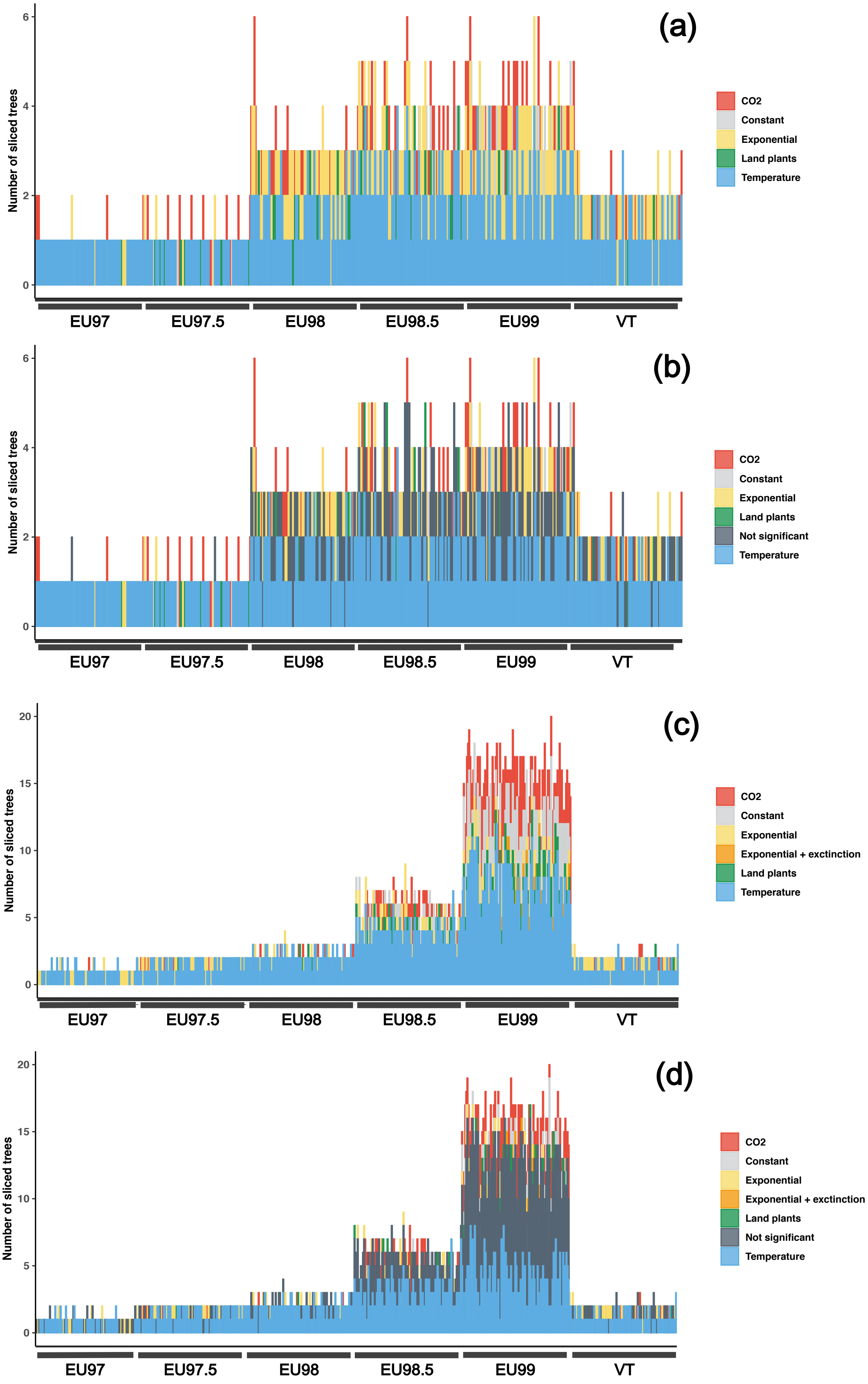

#### (II) Glomeromycotina crown age of 437 Myr:

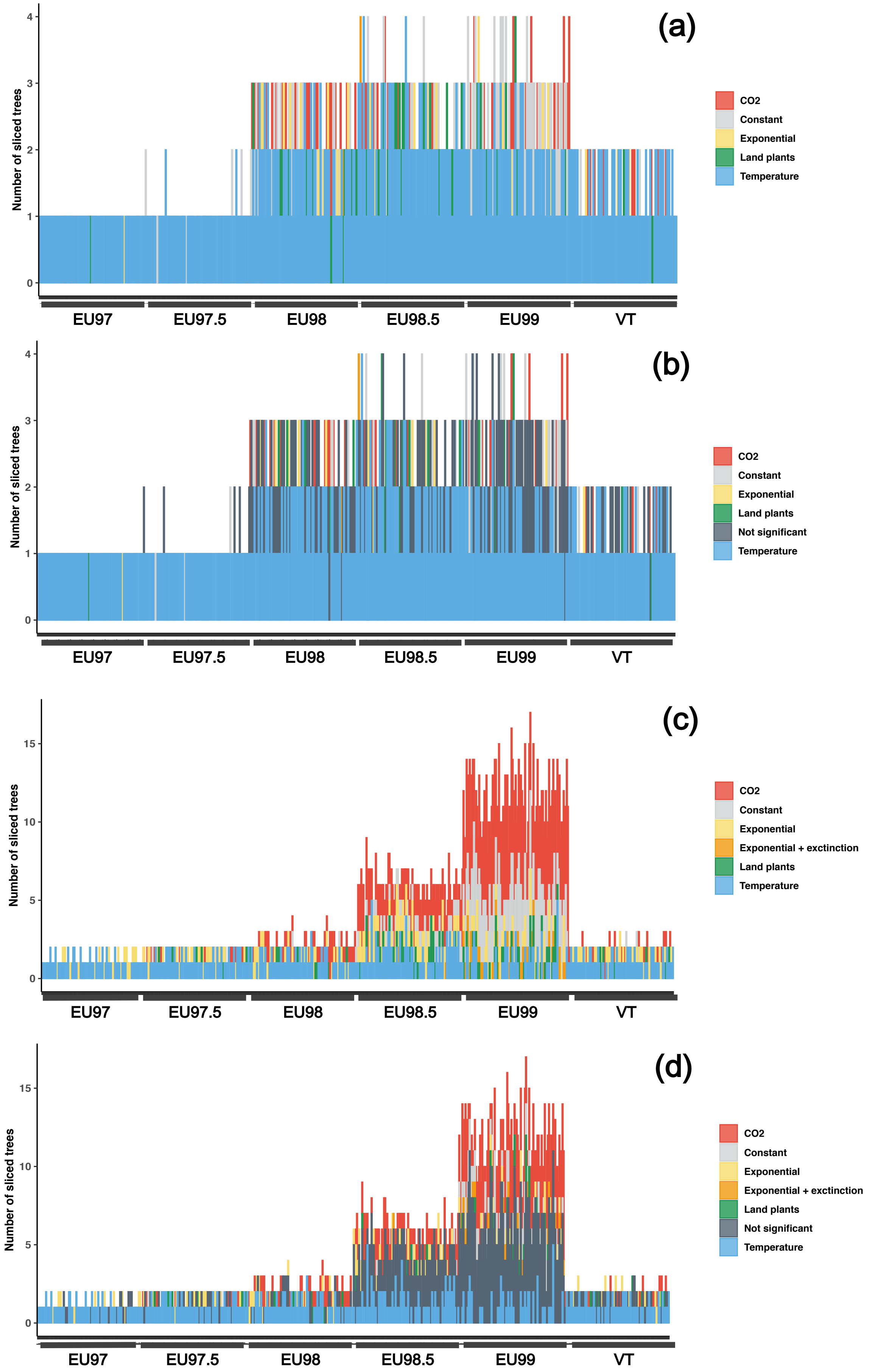

#### (III) Glomeromycotina crown age of 530 Myr:

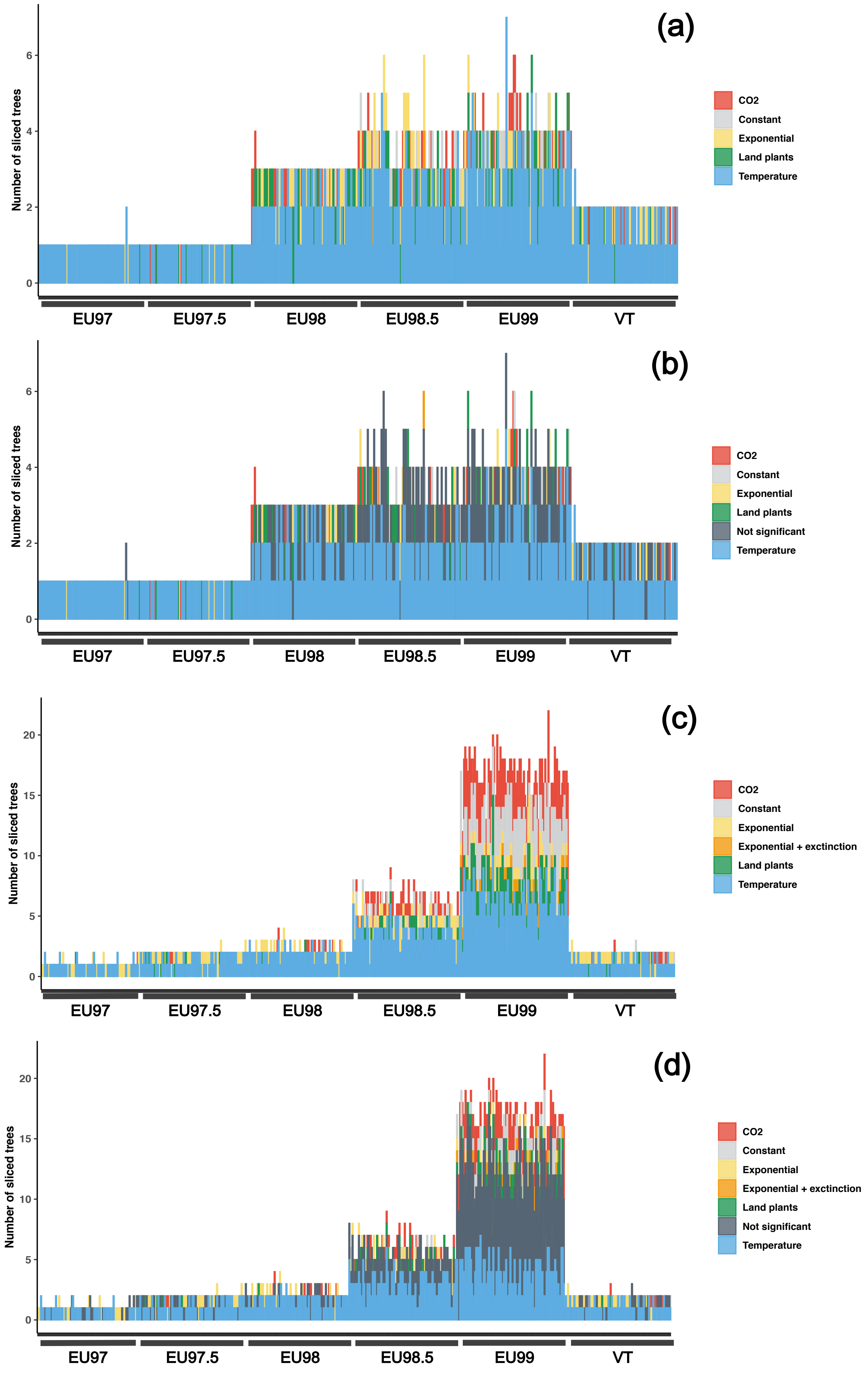

**Supplementary Figure 29: Diversification models estimated with RPANDA when using the 28S large sub-unit of the rRNA gene (LSU rRNA gene)**

Best-supported models between the temperature-dependent and time-dependent diversification model. For each delineation, the barplot indicates the model with the smallest AICc (a) or the model with the smallest AICc and an AICc difference with all others models larger than 2 (b). Glomeromycotina units were delineated based on GMYC delineation and diversification analyses were performed for a range of sampling fraction (90%, 80%, 70%, 60%, or 50% of sampled species) and replicated on the consensus trees and the 12 independent replicate trees.

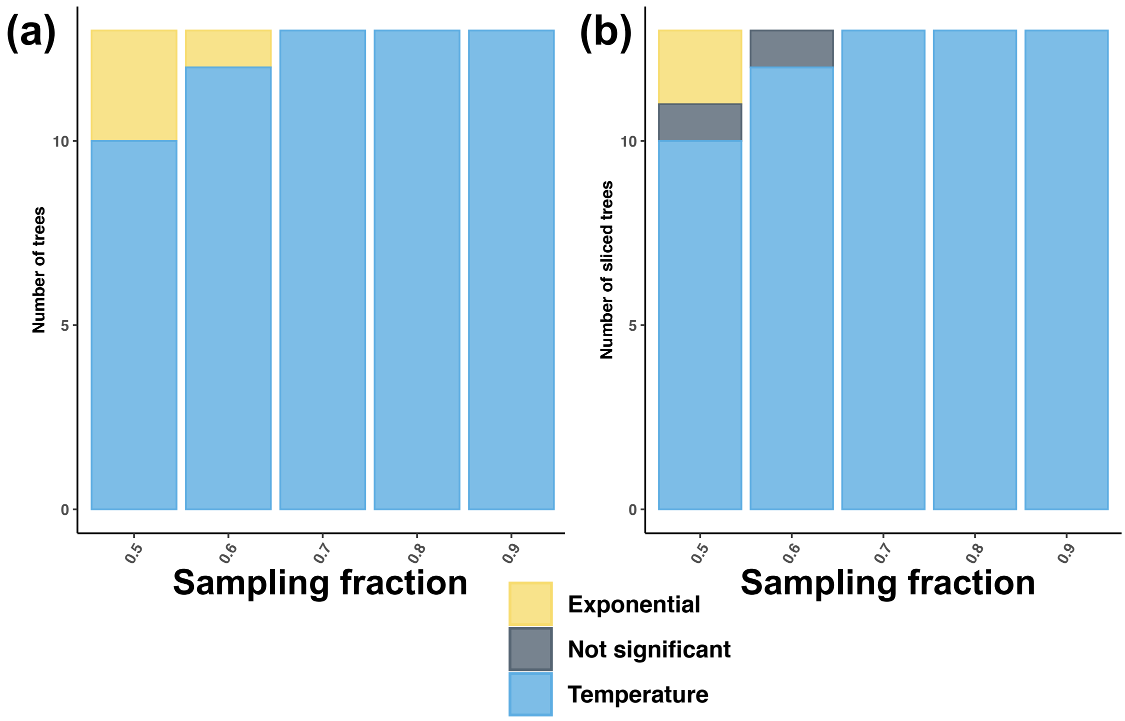

**Supplementary Figure 30: Characterizing Glomeromycotina niche width using principal coordinate analysis (PCA): Percentage of explained variance of the principal coordinate analysis (PCA)** performed for the different Glomeromycotina species with more than 10 sequences.

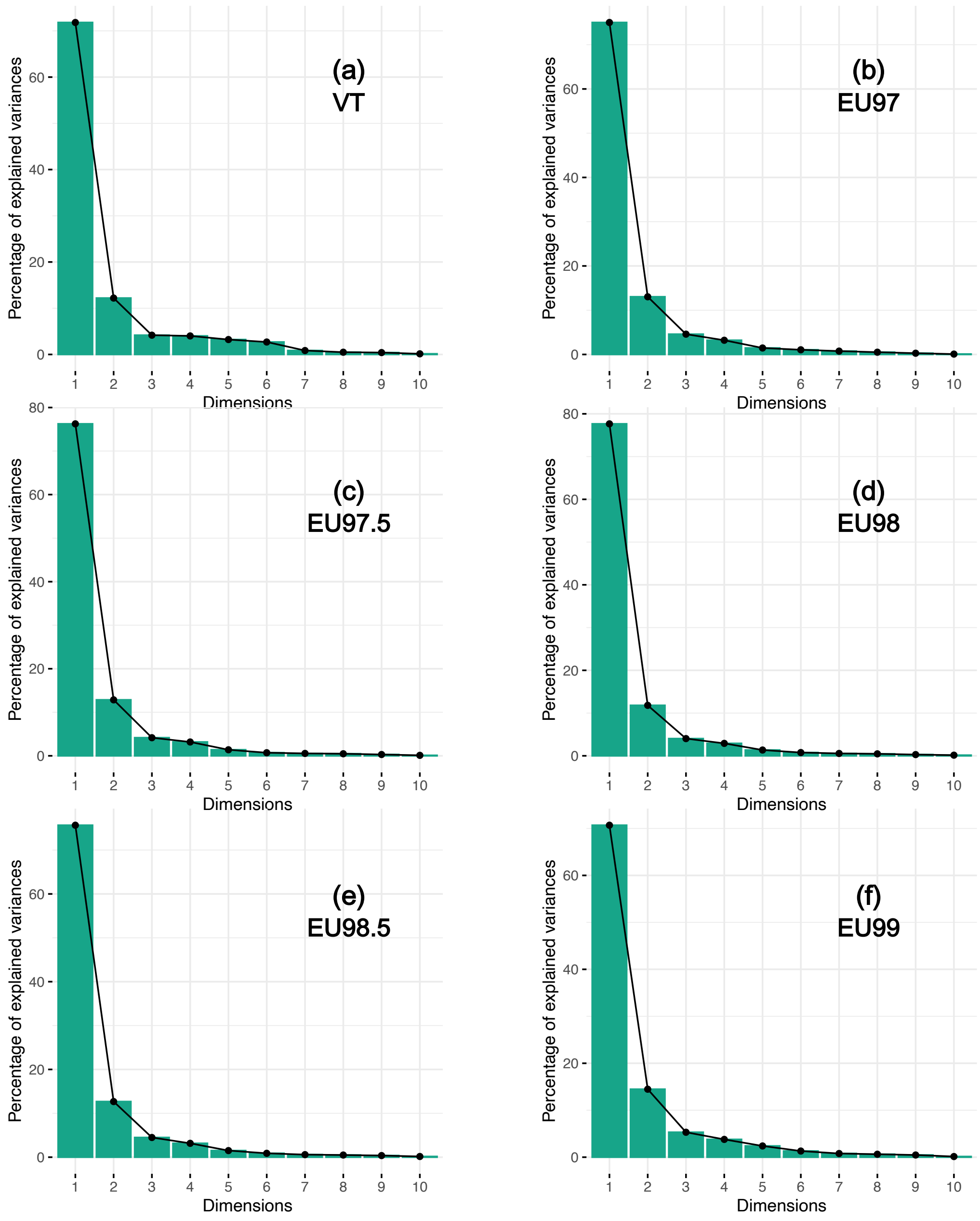

**Supplementary Figure 31: Characterizing Glomeromycotina niche width using principal coordinate analysis (PCA): Individual projection on the two principal coordinates according to the different Glomeromycotina delineations.**

Principal coordinate analysis (PCA) performed for the different species with more than 10 sequences. Colors represent the main fungal clades: Paraglomerales (including Archaeosporales), Diversisporales, Claroideoglomeraceae, and Glomeraceae.

Although the individual projections presented horseshoe shapes, the parallel analyses indicated that the two first components were significantly supported.

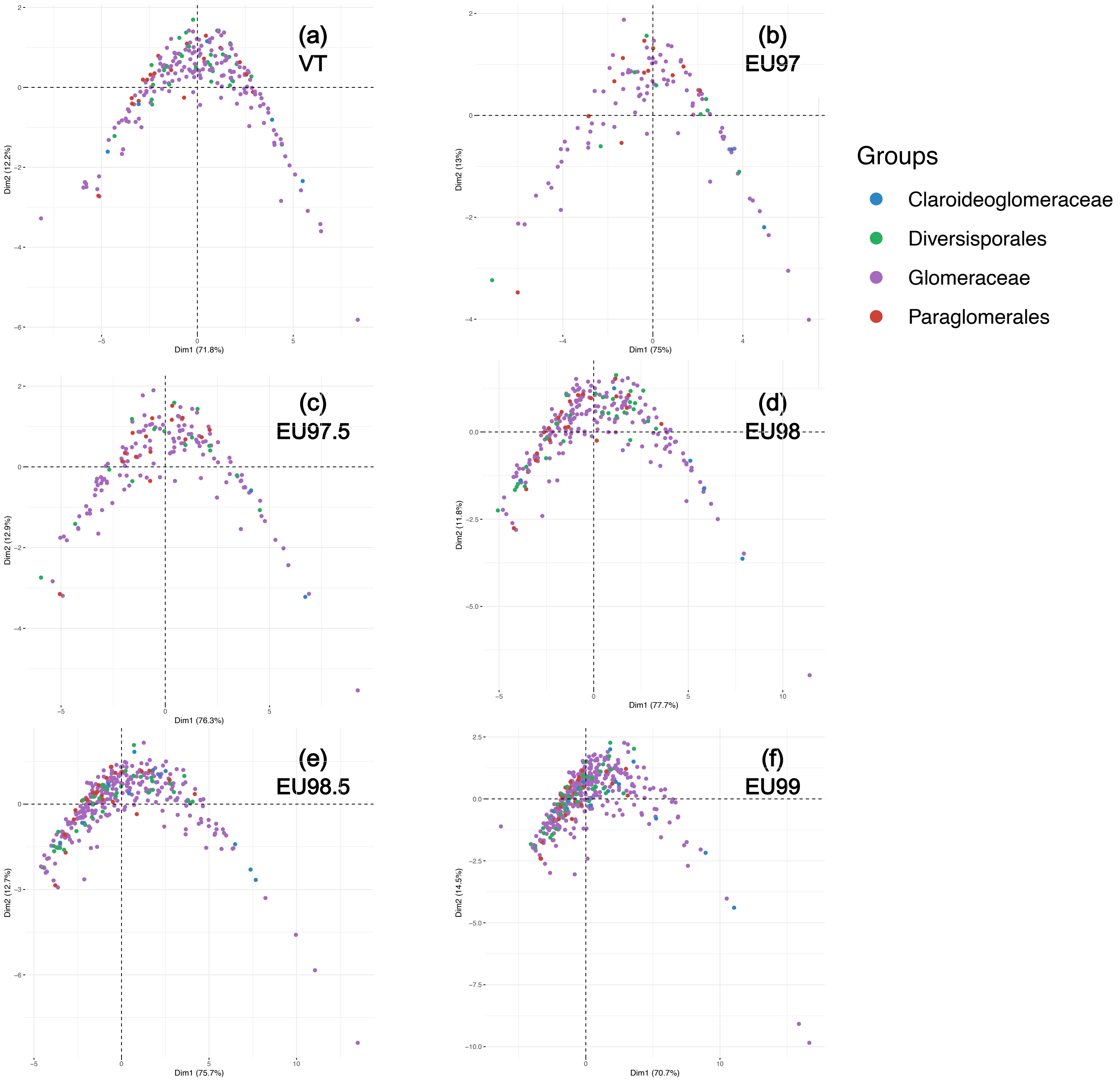

**Supplementary Figure 32: Characterizing Glomeromycotina niche width using principal coordinate analysis (PCA): Projection of the 10 abiotic and biotic variables on the two principal coordinates according to the different Glomeromycotina delineations.**

The principal coordinate analysis (PCA) was performed for species with more than 10 sequences. Colors represent the contribution of the variable to the principal coordinates. The percentage for each principal coordinate (PC) indicates its amount of explained variance.

Tested variables were: the number of continents (nb_continent), of realms (nb_realm), of ecosystems (nb_ecosystems), of habitats (nb_habitats), of biomes (nb_biomes), and of climatic zones (nb_climatic) that a given species occupies, as well as information about the associated plant species, such as the number of plant partners (nb_plants), the phylogenetic diversity of these plants (PD), and the betweenness and closeness measurement of each fungal species in the plant-fungus interaction network.

All variables are positively correlated with the first principal coordinate (PC1), whereas the second principal coordinate (PC2) is always positively correlated with the number of continents, and negatively correlated with the number of plants.

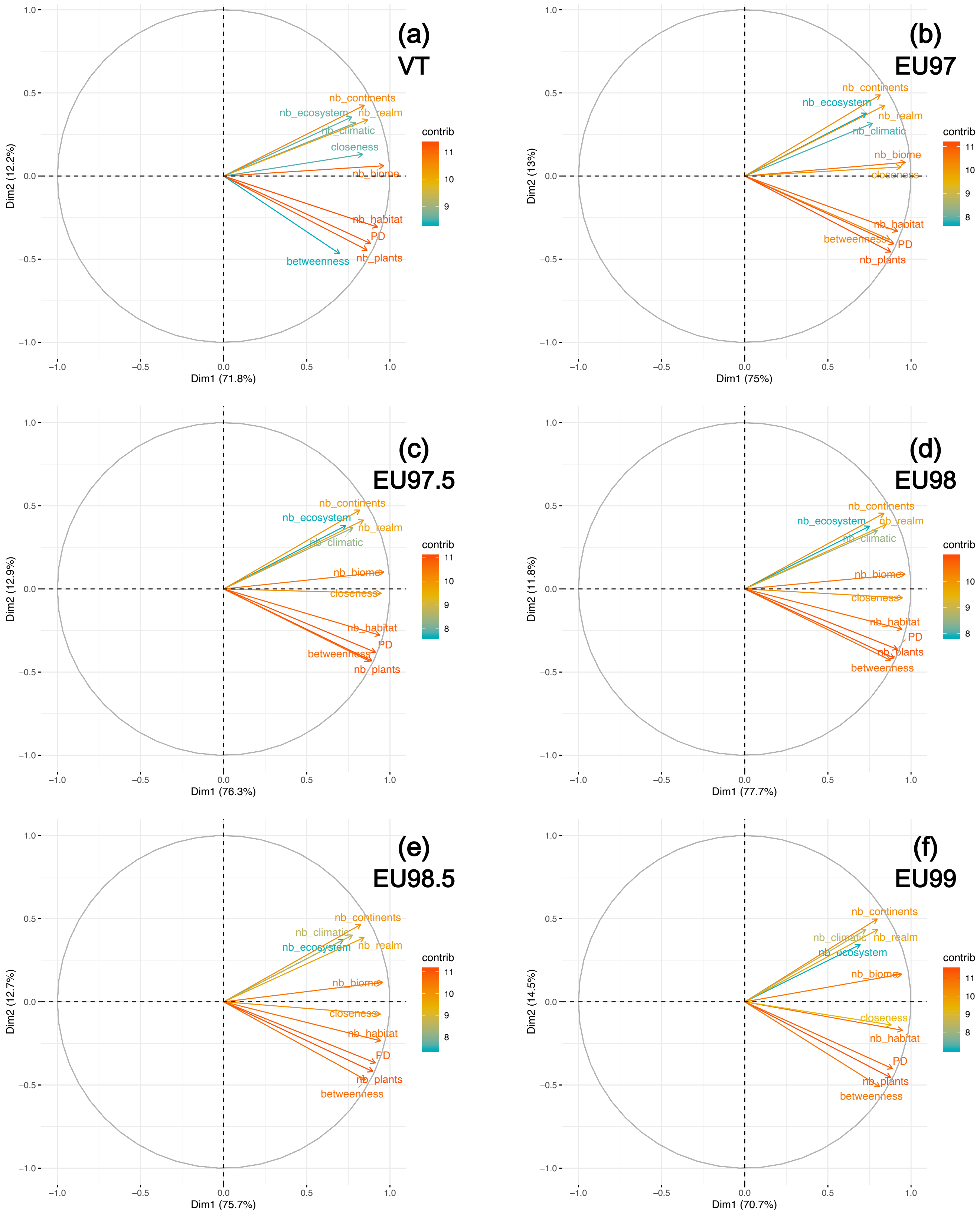
(a)
VT
(b)
EU97
(c)
EU97.5
(d)
EU98
(e)
EU98.5
(f)
EU99
nb_continents
nb_ecosystem
nb_realm
nb_climatic
closeness
nb_biome
nb_habitat
PD
nb_plants
betweenness
contrib
Dim1 (71.8%)
Dim2 (12.2%)
Dim1 (75%)
Dim2 (13%)
Dim1 (76.3%)
Dim2 (12.9%)
Dim1 (77.7%)
Dim2 (11.8%)
Dim1 (75.7%)
Dim2 (12.7%)
Dim1 (70.7%)
Dim2 (14.5%)

**Supplementary Figure 33: Correlations between speciation rates at the tips, estimates of genetic diversity (Tajima's $\theta\pi$ estimator - referred to as "Nei diversity") and PC1 and PC2 coordinates**, evaluated using simple linear mixed models (a - not correcting for phylogenetic relatedness) or MCMCglmm (b - correcting for phylogenetic relatedness). For each Glomeromycotina delineation, the statistical analyses are performed on the consensus tree and the 12 replicates, and we reported here the number of trees that present a significant correlation between the set of tested variables (in blue: significant positive correlation, in red: significant negative correlation). We tested the robustness of the analyses to the sampling within species, by performing the analyses on all species being represented by at least 10, 15, or 20 sequences occurring in natural environments. The columns indicate the significance and sign of the correlations between (i) PCA1 and genetic diversity ($\theta\pi$), (ii) PCA2 and genetic diversity ($\theta\pi$), (iii) PCA1 and lineage-specific speciation rates, (iv) PCA2 and lineage-specific speciation rates, and (v) genetic diversity and lineage-specific speciation rates.

**(a)**

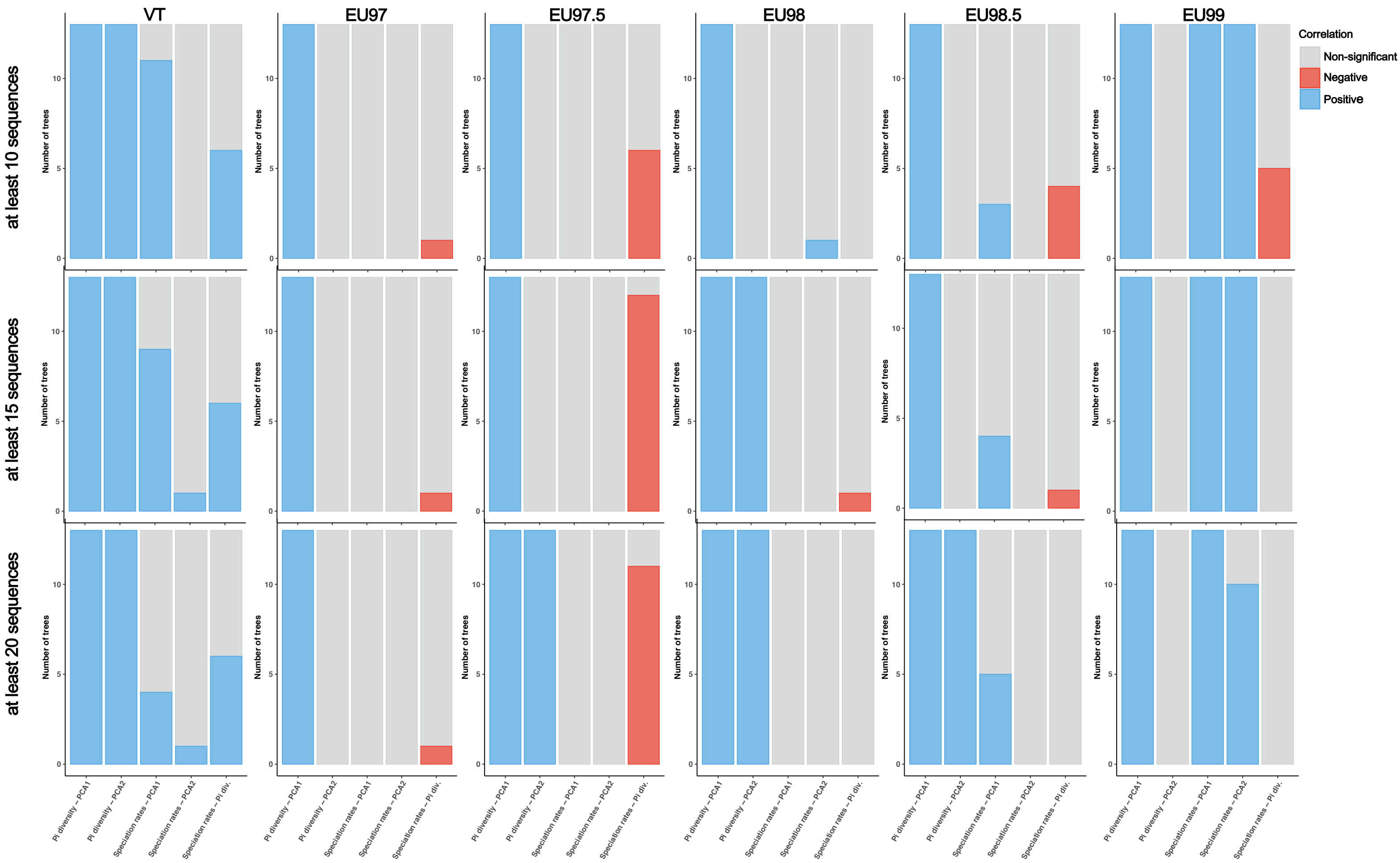

VT
EU97
EU97.5
EU98
EU98.5
EU99
Correlation
Non-significant
Negative
Positive
at least 10 sequences
at least 15 sequences
at least 20 sequences
Number of trees
Pi diversity ~ PCA1
Pi diversity ~ PCA2
Speciation rates ~ PCA1
Speciation rates ~ PCA2
Speciation rates ~ Pi div.

**(b)**

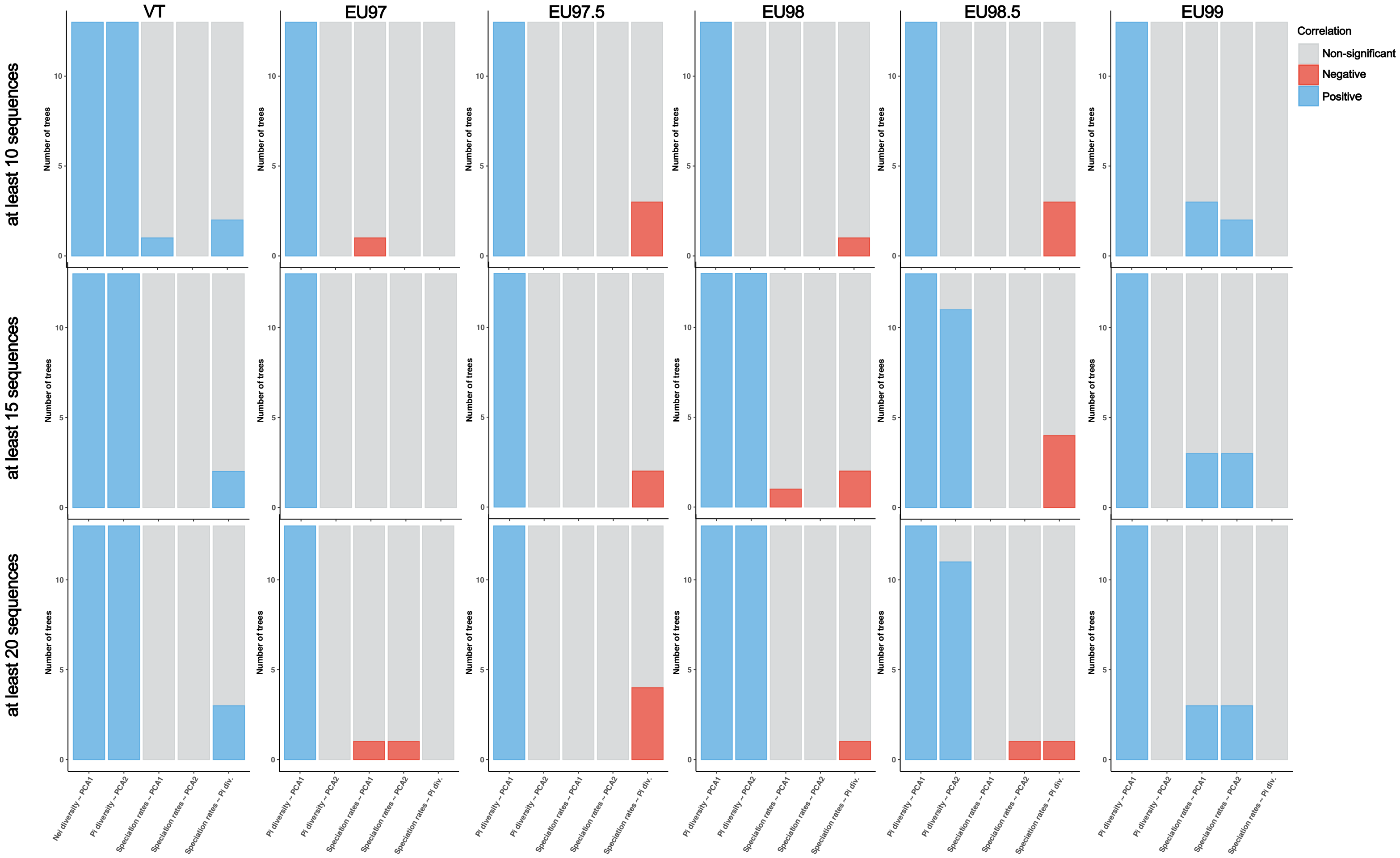
VT
EU97
EU97.5
EU98
EU98.5
EU99
Correlation
Non-significant
Negative
Positive
at least 10 sequences
at least 15 sequences
at least 20 sequences
Number of trees
Nei diversity – PCA1
Pi diversity – PCA1
Pi diversity – PCA2
Speciation rates – PCA1
Speciation rates – PCA2
Speciation rates – Pi div.

**Supplementary Figure 34: The *Rhizophagus* clade with large niche width present the highest speciation rates (Fig. 2).**

**(a-b)** Speciation rates in the diversified *Rhizophagus* clade versus other Glomeromycotina clades, in the VT (a) and EU99 (b) delineations. Only the results for the consensus trees are represented here.

**(c-d)** PC1 coordinates (as a proxy for Glomeromycotina niche width) in the diversified *Rhizophagus* clade versus other Glomeromycotina clades, in the VT (a) and EU99 (b) delineations. PC1 coordinates were only computed for Glomeromycotina species represented by more than 10 sequences.

Boxplots indicate the median surrounded by the first and third quartiles, and whiskers extend to the extreme values but no further than 1.5 of the inter-quartile range.

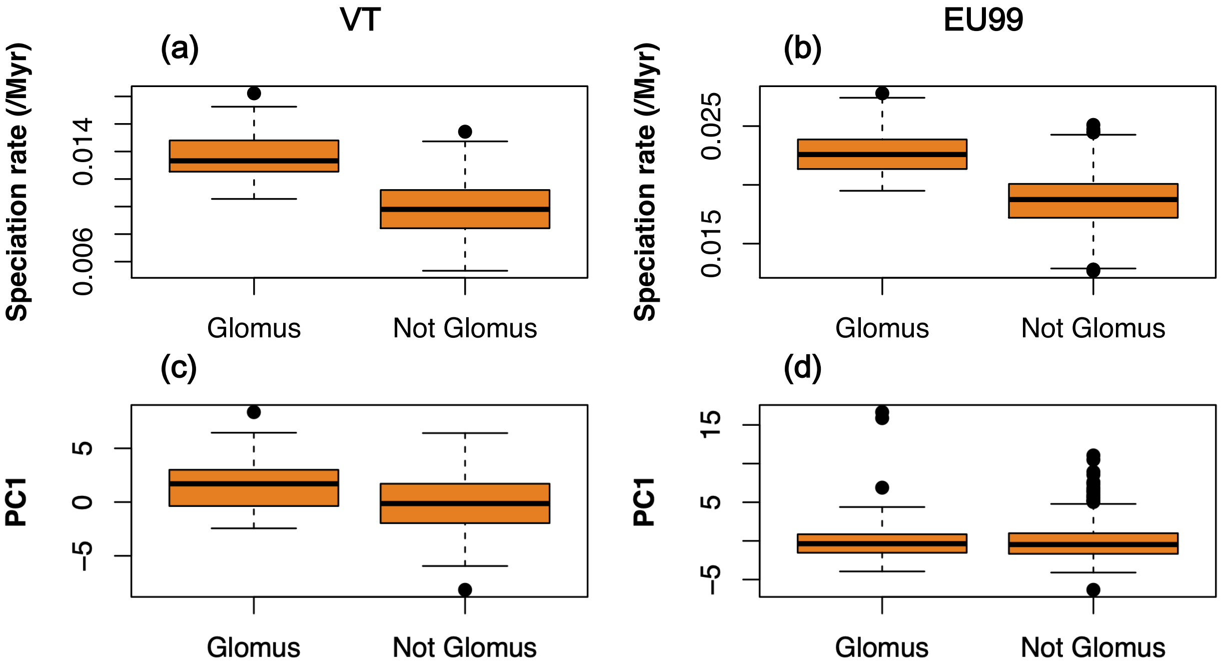

**Supplementary Figure 35: Significant latitudinal gradient of Glomeromycotina diversity.**

Number of Glomeromycotina species observed as a function of the absolute latitude. To account for the heterogenous sampling coverage, 1,000 Glomeromycotina sequences are subsampled by slice of latitude (every 20 degrees). We replicated the subsampling 100 times and represented the observed number of Glomeromycotina species per slice of latitude using boxplots.

Boxplots indicate the median surrounded by the first and third quartiles, and whiskers extend to the extreme values but no further than 1.5 of the inter-quartile range.

The lower diversity in latitude between 0 and 20 degrees for EU99 might indicate that the sampling coverage of the tropics is not enough to get the whole species richness of the area when the Glomeromycotina delineation is finer.

The letters (from "a" to "d") indicates which slides of latitude are significantly different from the others (after correcting for multiple testing).

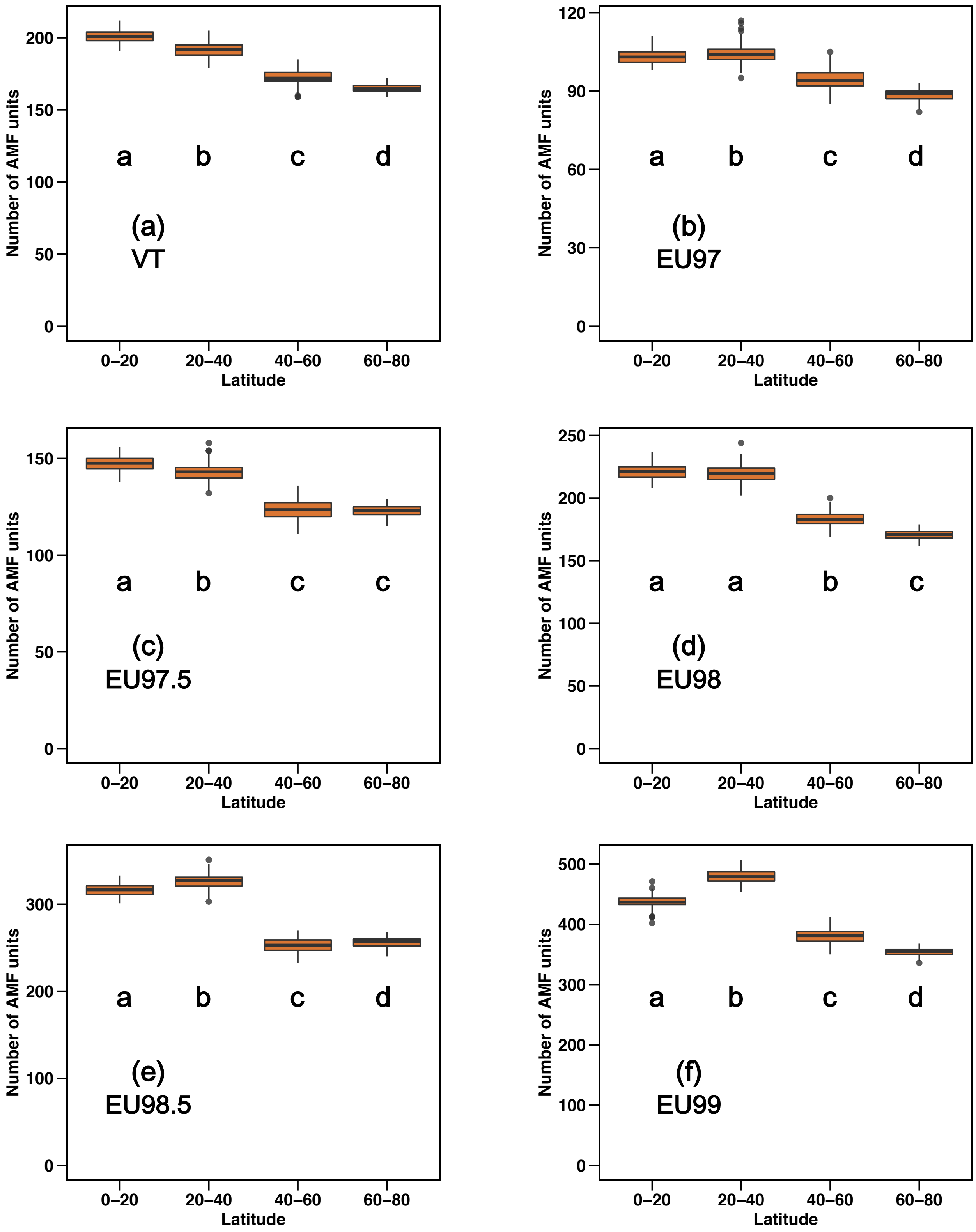

(a)
VT
(b)
EU97
(c)
EU97.5
(d)
EU98
(e)
EU98.5
(f)
EU99
Number of AMF units
Latitude
0–20
20–40
40–60
60–80
a
b
c
d

**Supplementary Figure 36: The total number of Glomeromycotina species is not higher in (tropical) grasslands.**

Total number of VT or EU units as a function of the type of ecosystem. To account for the heterogenous sampling coverage, we rarefied the dataset in order to have the same number of observations per ecosystem (250 per ecosystem), and replicated our rarefactions 100 times) and represented the observed number of Glomeromycotina species per ecosystem using boxplots.

Boxplots indicate the median surrounded by the first and third quartiles, and whiskers extend to the extreme values but no further than 1.5 of the inter-quartile range.

Although in local communities, Glomeromycotina alpha diversity in tropical grasslands is generally higher than Glomeromycotina alpha diversity in tropical forests, Glomeromycotina beta diversity in tropical forests is much higher (Davison *et al.*, 2015), which explains why the total diversity between tropical grasslands and forests is similar.

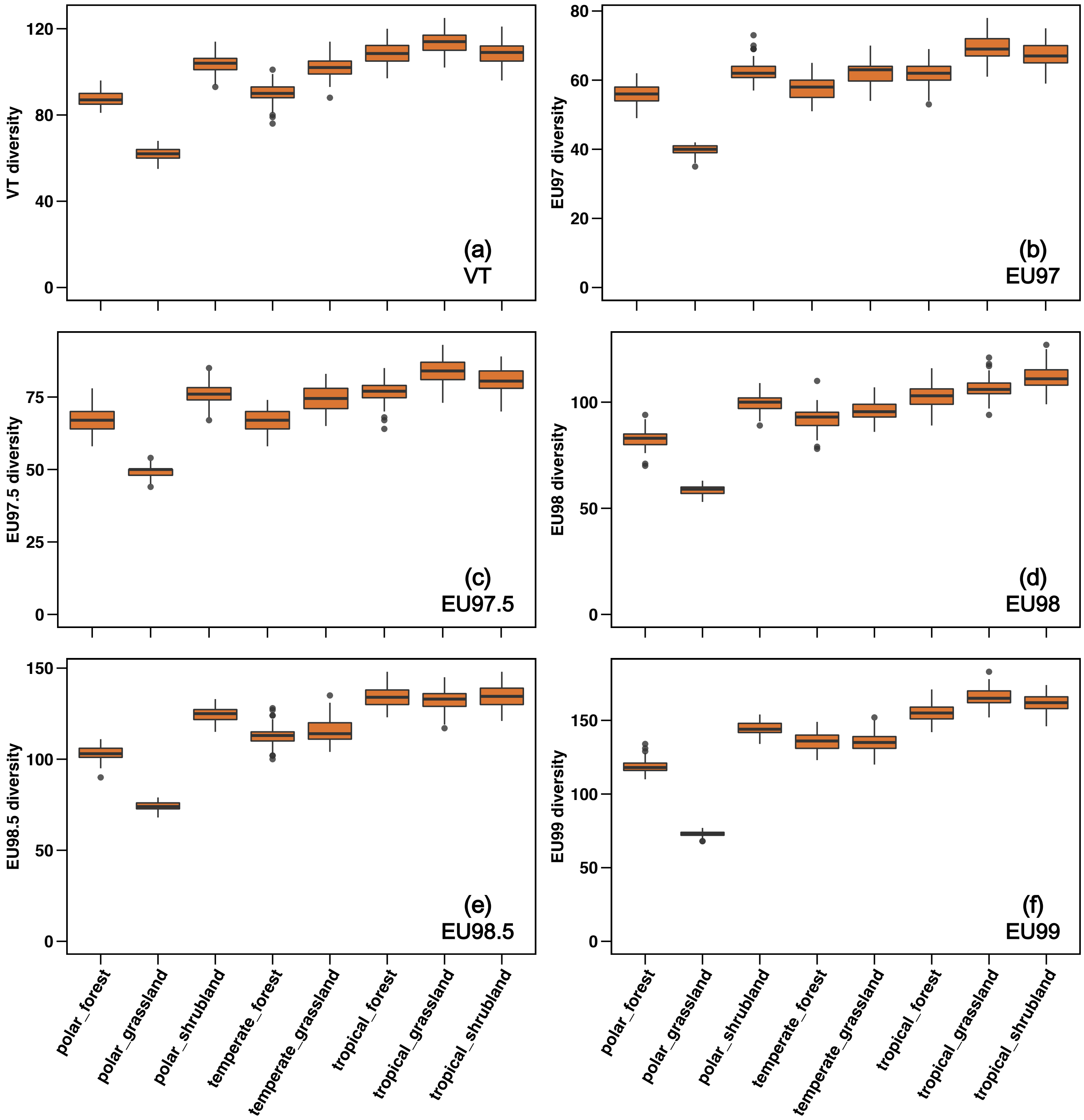

VT diversity
EU97 diversity
EU97.5 diversity
EU98 diversity
EU98.5 diversity
EU99 diversity
(a) VT
(b) EU97
(c) EU97.5
(d) EU98
(e) EU98.5
(f) EU99
polar_forest
polar_grassland
polar_shrubland
temperate_forest
temperate_grassland
tropical_forest
tropical_grassland
tropical_shrubland

**Supplementary Figure 37: No effect of ecosystem types or climatic zones on Glomeromycotina speciation rates.**

For each delineation, the speciation rates estimated by ClaDS (with the BDES sampling fraction on the Glomeromycotina consensus tree) are sorted as a function of the ecosystems and the climatic niche that the corresponding Glomeromycotina accession occupy. Ecosystem types and climatic zones appear to have no significant effect on speciation rates (phylogenetic linear mixed models: p-values>0.05).

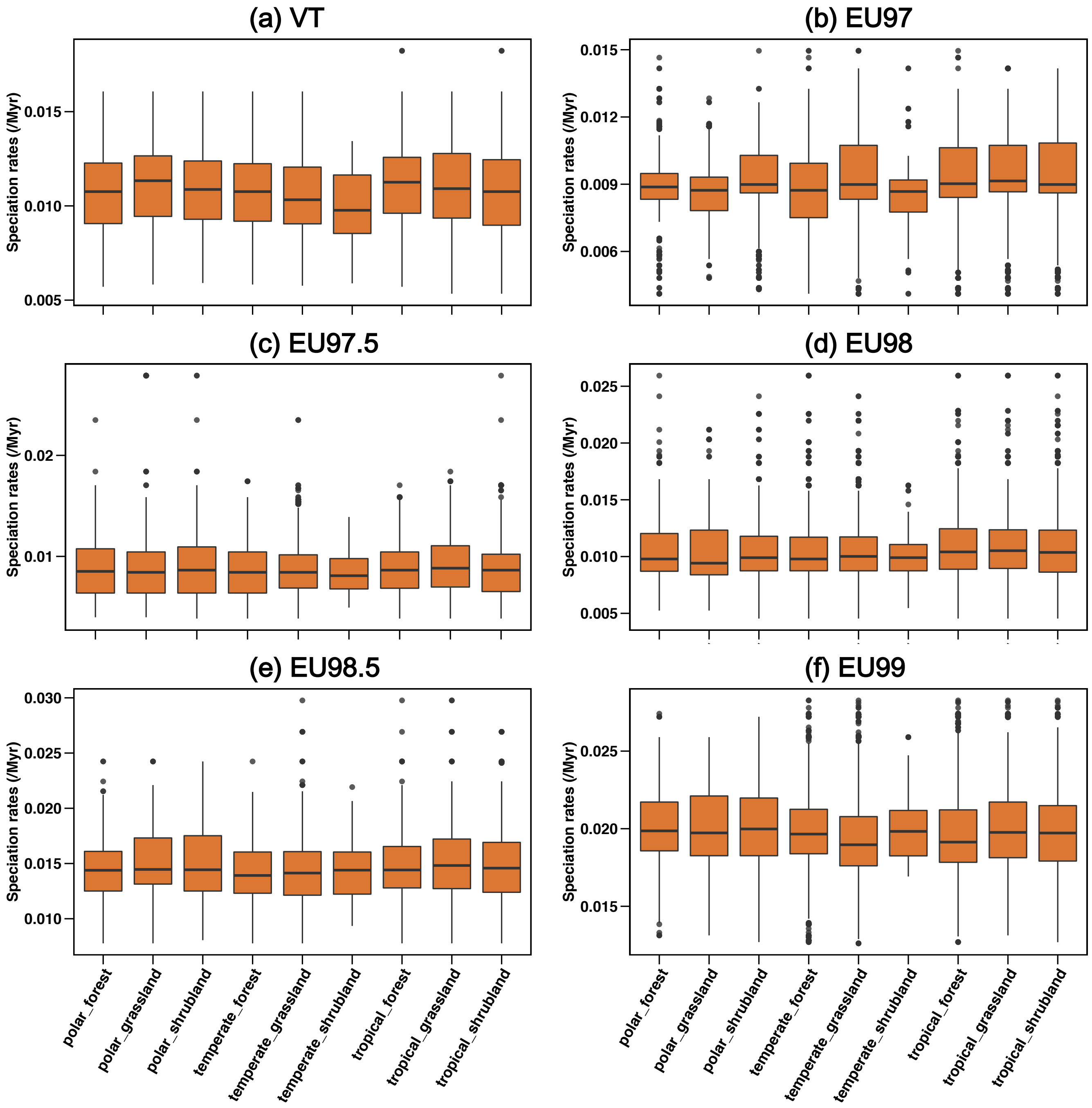

**Supplementary Figure 38: No significant effect of mean spore length on VT speciation rates.**

Each panel represents the consensus tree or one on the 12 replicate tree (ordered top to bottom from left to right). The relationship between speciation rates (log-transformed) and mean spore lengths (log-transformed) is evaluated using Phylogenetic Generalized Least-Square model (PGLS) to take into account phylogenetic relatedness among VT. We checked the normality of the residuals of the models.

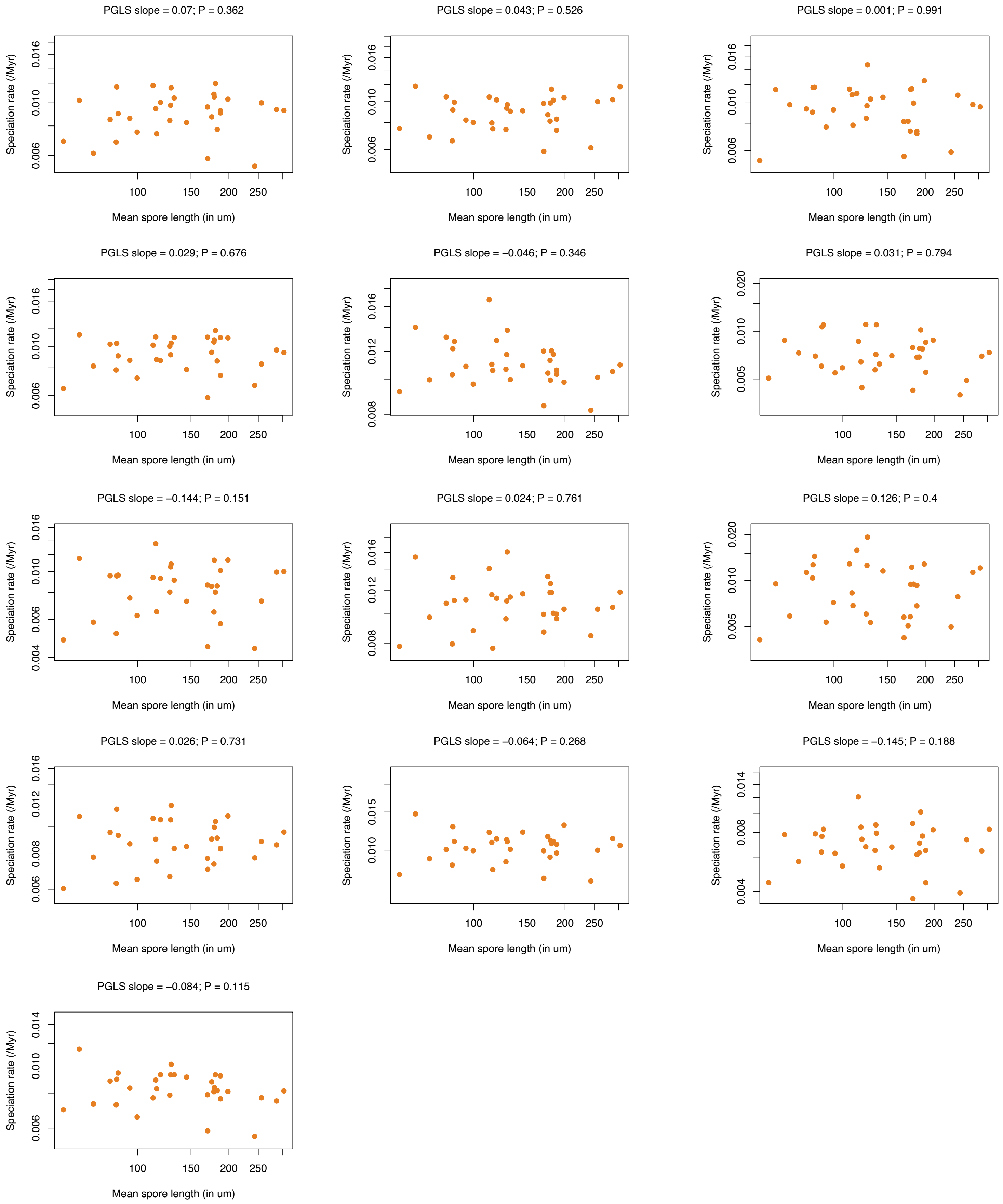
PGLS slope = 0.07; P = 0.362
PGLS slope = 0.043; P = 0.526
PGLS slope = 0.001; P = 0.991
PGLS slope = 0.029; P = 0.676
PGLS slope = −0.046; P = 0.346
PGLS slope = 0.031; P = 0.794
PGLS slope = −0.144; P = 0.151
PGLS slope = 0.024; P = 0.761
PGLS slope = 0.126; P = 0.4
PGLS slope = 0.026; P = 0.731
PGLS slope = −0.064; P = 0.268
PGLS slope = −0.145; P = 0.188
PGLS slope = −0.084; P = 0.115
Speciation rate (/Myr)
Mean spore length (in um)
100
150
200
250

**Supplementary Figure 39: No significant correlation between mean spore length and endemism.**

The relationship between mean spore length and the number of continents occupied by each VT is evaluated using Phylogenetic Generalized Least-Square model (PGLS).

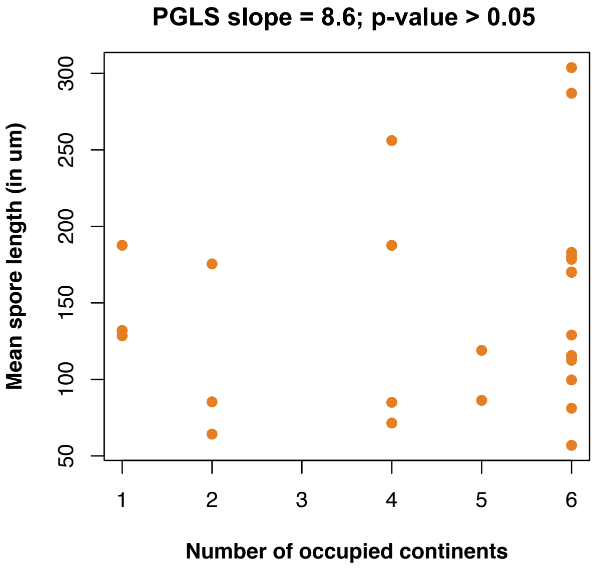

**Supplementary Figure 40: Average Glomeromycotina speciation rates and land plant diversity are decoupled for ~130 Myr:**

Average speciation rates (MAPS) through time estimated by ClaDS, for the different Glomeromycotina delineations (a: VT and b: EU99), using the BDES estimated sampling fraction (Quince et al., 2008). The consensus trees are in grey, whereas orange lines represent the 12 independent replicate trees.

The green dots and line represent the land plant fossil diversity (data up to 430 Myr ago). Land plant diversity were estimated using the Paleobiology database and the shareholder quorum subsampling method (Supplementary Methods 6; quorum value of 0.5). We smoothed the curves using cubic splines: the degree of freedom was 25 for the land plant diversity.

The figures indicate that Glomeromycotina diversification and land plant fossil diversity followed similar trends from 400 Myr to 130 Myr ago, suggesting a dynamic of co-diversification (Lutzoni et al., 2018). Conversely from 130 Myr ago to the present (highlighted in orange), the Glomeromycotina speciation rates decreased, while the land plant diversities peaked (corresponding to the radiations of many Angiosperm or Pinaceae lineages). This period likely corresponds to a decoupling of Glomeromycotina and plant diversifications. These results contrast with diversification dynamics in other mycorrhizal fungal groups that present higher rates of diversification close to the present, mirroring the recent land plant diversifications (Lutzoni et al., 2018; Varga et al., 2019).

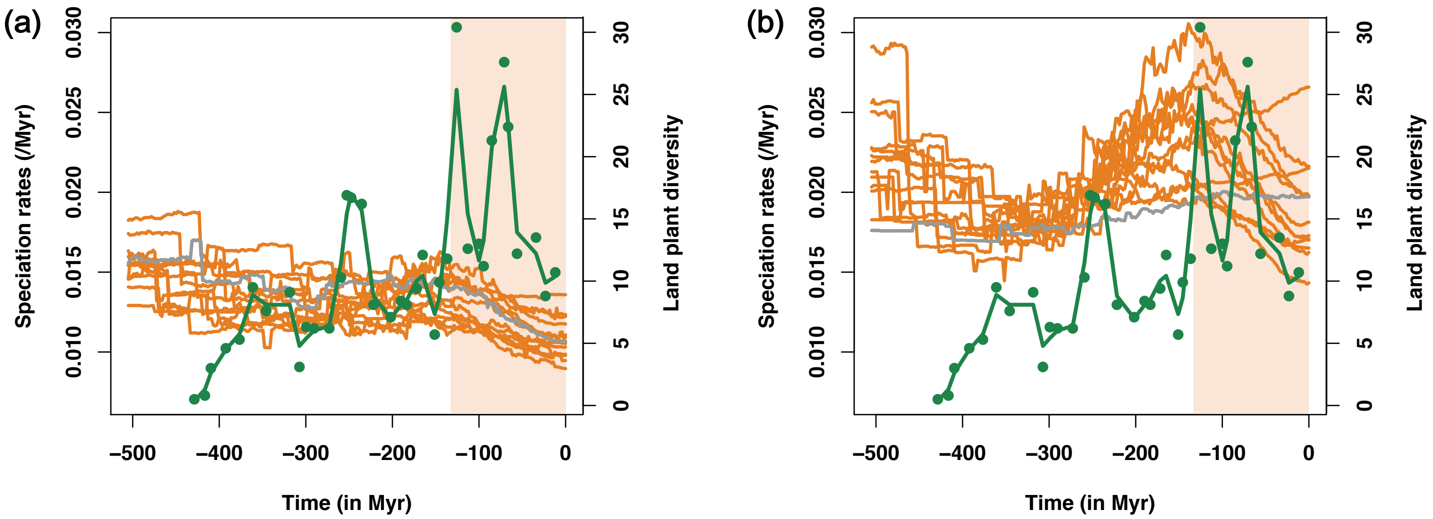

#### Supplementary References:


Alroy, J. (2010). Geographical, environmental and intrinsic biotic controls on Phanerozoic marine diversification. *Palaeontology*, *53*(6), 1211–1235. doi:10.1111/j.1475-4983.2010.01011.x

Bruns, T. D., Corradi, N., Redecker, D., Taylor, J. W., & Öpik, M. (2018). Glomeromycotina: what is a species and why should we care? *New Phytologist*, *220*(4), 963–967. doi:10.1111/nph.14913

Bueno, C. G., & Moora, M. (2019). How do arbuscular mycorrhizal fungi travel? *New Phytologist*, *222*(2), 645–647. doi:10.1111/nph.15722

Buja, A., & Eyuboglu, N. (1992). Remarks on parallel analysis. *Multivariate Behavioral Research*, *27*(4), 509–540. doi:10.1207/s15327906mbr2704_2

Capella-Gutierrez, S., Silla-Martinez, J. M., & Gabaldon, T. (2009). trimAl: a tool for automated alignment trimming in large-scale phylogenetic analyses. *Bioinformatics*, *25*(15), 1972–1973. doi:10.1093/bioinformatics/btp348

Clavel, J., & Morlon, H. (2017). Accelerated body size evolution during cold climatic periods in the Cenozoic. *Proceedings of the National Academy of Sciences of the United States of America*, *114*(16), 4183–4188. doi:10.1073/pnas.1606868114

Cleal, C. J., & Cascales-Miñana, B. (2014). Composition and dynamics of the great Phanerozoic Evolutionary Floras. *Lethaia*, *47*(4), 469–484. doi:10.1111/let.12070

Condamine, F. L., Rolland, J., & Morlon, H. (2019). Assessing the causes of diversification slowdowns: temperature-dependent and diversity-dependent models receive equivalent support. *Ecology Letters*, *22*(11), 1900–1912. doi:10.1111/ele.13382

Davison, J., Moora, M., Öpik, M., Adholeya, A., Ainsaar, L., Bâ, A., … Zobel, M. (2015). Global assessment of arbuscular mycorrhizal fungus diversity reveals very low endemism. *Science*, *349*(6251), 970–973. doi:10.1126/science.aab1161

Dormann, C. F., Gruber, B., & Fründ, J. (2008). Introducing the bipartite package: analysing ecological networks. *R News*, *8*(October), 8–11. doi:10.1159/000265935

Edgar, R. C. (2013). UPARSE: Highly accurate OTU sequences from microbial amplicon reads. *Nature Methods*, *10*(10), 996–998. doi:10.1038/nmeth.2604

Ferretti, L., Raineri, E., & Ramos-Onsins, S. (2012). Neutrality tests for sequences with missing data. *Genetics*, *191*(4), 1397–1401. doi:10.1534/genetics.112.139949

Fruciano, C. (2020). GeometricMorphometricsMix: Miscellaneous functions useful for geometric morphometrics, R package version 1.0-0. R package.

Kalyaanamoorthy, S., Minh, B. Q., Wong, T. K. F., von Haeseler, A., & Jermiin, L. S. (2017). ModelFinder: fast model selection for accurate phylogenetic estimates. *Nature Methods*, *14*(6), 587–589. doi:10.1038/nmeth.4285

Katoh, K., & Standley, D. M. (2013). MAFFT Multiple sequence alignment software version 7: Improvements in performance and usability. *Molecular Biology and Evolution*, *30*(4), 772–780. doi:10.1093/molbev/mst010

Kembel, S. W., Cowan, P. D., Helmus, M. R., Cornwell, W. K., Morlon, H., Ackerly, D. D., … Webb, C. O. (2010). Picante: R tools for integrating phylogenies and ecology. *Bioinformatics*, *26*(11), 1463–1464. doi:10.1093/bioinformatics/btq166

Krüger, M., Krüger, C., Walker, C., Stockinger, H., & Schüßler, A. (2012). Phylogenetic reference data for systematics and phylotaxonomy of arbuscular mycorrhizal fungi from phylum to species level. *New Phytologist*, *193*(4), 970–984. doi:10.1111/j.1469-8137.2011.03962.x

Lekberg, Y., Vasar, M., Bullington, L. S., Sepp, S.-K. K., Antunes, P. M., Bunn, R., … Öpik, M. (2018). More bang for the buck? Can arbuscular mycorrhizal fungal communities be characterized adequately alongside other fungi using general fungal primers? *New Phytologist*, *220*(4), 971–976. doi:10.1111/nph.15035

Lewitus, E., Bittner, L., Malviya, S., Bowler, C., & Morlon, H. (2018). Clade-specific diversification dynamics of marine diatoms since the Jurassic. *Nature Ecology and Evolution*, *2*(11), 1715–1723. doi:10.1038/s41559-018-0691-3

Lewitus, E., & Morlon, H. (2018). Detecting environment-dependent diversification from phylogenies: A simulation study and some empirical illustrations. *Systematic Biology*, *67*(4), 576–593. doi:10.1093/sysbio/syx095

Louca, S., & Pennell, M. W. (2020). Extant timetrees are consistent with a myriad of diversification histories. *Nature, 580*(7804), 502–505. doi:10.1038/s41586-020-2176-1

Lutzoni, F., Nowak, M. D., Alfaro, M. E., Reeb, V., Miadlikowska, J., Krug, M., … Magallón, S. (2018). Contemporaneous radiations of fungi and plants linked to symbiosis. *Nature Communications*, *9*(1), 1–11. doi:10.1038/s41467-018-07849-9

Morlon, H., Lewitus, E., Condamine, F. L., Manceau, M., Clavel, J., & Drury, J. (2016). RPANDA: An R package for macroevolutionary analyses on phylogenetic trees. *Methods in Ecology and Evolution*, *7*(5), 589–597. doi:10.1111/2041-210X.12526

Morlon, H., O'Connor, T. K., Bryant, J. A., Charkoudian, L. K., Docherty, K. M., Jones, E., … Bohannan, B. J. M. (2015). The biogeography of putative microbial antibiotic production. *PLOS ONE*, *10*(6), e0130659. doi:10.1371/journal.pone.0130659

Morlon, H., Robin, S., & Hartig, F. (2022). Studying speciation and extinction dynamics from phylogenies: addressing identifiability issues. *Trends in Ecology & Evolution*, 1–10. doi:10.1016/j.tree.2022.02.004

Muñoz-Pajares, A. J. (2013). SIDIER: Substitution and indel distances to infer evolutionary relationships. *Methods in Ecology and Evolution*, *4*(12), 1195–1200. doi:10.1111/2041-210X.12118

Niklas, K. J. (1988). Patterns of vascular plant diversification in the fossil record: Proof and conjecture. *Annals of the Missouri Botanical Garden*, *75*(1), 35. doi:10.2307/2399465

Niklas, K. J., Tiffney, B. H., & Knoll, A. H. (1983). Patterns in vascular land plant diversification. *Nature*, *303*(5918), 614–616. doi:10.1038/303614a0

Öpik, M., Vanatoa, A., Vanatoa, E., Moora, M., Davison, J., Kalwij, J. M., … Zobel, M. (2010). The online database MaarjAM reveals global and ecosystemic distribution patterns in arbuscular mycorrhizal fungi (Glomeromycota). *New Phytologist*, *188*(1), 223–241. doi:10.1111/j.1469-8137.2010.03334.x

Perez-Lamarque, B., Selosse, M. A., Öpik, M., Morlon, H., & Martos, F. (2020).

Cheating in arbuscular mycorrhizal mutualism: a network and phylogenetic analysis of mycoheterotrophy. *New Phytologist*, *226*(6), 1822–1835. doi:10.1111/nph.16474

Powell, J. R., Monaghan, M. T., Öpik, M., & Rillig, M. C. (2011). Evolutionary criteria outperform operational approaches in producing ecologically relevant fungal species inventories. *Molecular Ecology*, *20*(3), 655–666. doi:10.1111/j.1365-294X.2010.04964.x

Price, M. N., Dehal, P. S., & Arkin, A. P. (2009). Fasttree: Computing large minimum evolution trees with profiles instead of a distance matrix. *Molecular Biology and Evolution*, *26*(7), 1641–1650. doi:10.1093/molbev/msp077

Quince, C., Curtis, T. P., & Sloan, W. T. (2008). The rational exploration of microbial diversity. *ISME Journal*, *2*(10), 997–1006. doi:10.1038/ismej.2008.69

R Core Team. (2020). R: A language and environment for statistical computing. Vienna, Austria: R Foundation for Statistical Computing.

Rabosky, D. L. (2016). Challenges in the estimation of extinction from molecular phylogenies: A response to Beaulieu and O'Meara. *Evolution*, *70*(1), 218–228. doi:10.1111/evo.12820

Rabosky, D. L., & Lovette, I. J. (2008). Density-dependent diversification in North American wood warblers. *Proceedings of the Royal Society B: Biological Sciences*, *275*(1649), 2363–2371. doi:10.1098/rspb.2008.0630

Rambaut, A., Drummond, A. J., Xie, D., Baele, G., & Suchard, M. A. (2018). Posterior summarization in Bayesian phylogenetics using Tracer 1.7. *Systematic Biology*, *67*(5), 901–904. doi:10.1093/sysbio/syy032

Rimington, W. R., Pressel, S., Duckett, J. G., Field, K. J., Read, D. J., & Bidartondo, M. I. (2018). Ancient plants with ancient fungi: liverworts associate with early-diverging arbuscular mycorrhizal fungi. *Proceedings of the Royal Society B: Biological Sciences*, *285*(1888), 20181600. doi:10.1098/rspb.2018.1600

Rognes, T., Flouri, T., Nichols, B., Quince, C., & Mahé, F. (2016). VSEARCH: A versatile open source tool for metagenomics. *PeerJ*, *2016*(10), e2584.

doi:10.7717/peerj.2584

Russel, P. M., Brewer, B. J., Klaere, S., & Bouckaert, R. R. (2019). Model selection and parameter inference in phylogenetics using nested sampling. *Systematic Biology*, *68*(2), 219–233. doi:10.1093/sysbio/syy050

Savary, R., Masclaux, F. G., Wyss, T., Droh, G., Cruz Corella, J., Machado, A. P., … Sanders, I. R. (2018). A population genomics approach shows widespread geographical distribution of cryptic genomic forms of the symbiotic fungus Rhizophagus irregularis. *ISME Journal*, *12*(1), 17–30. doi:10.1038/ismej.2017.153

Sigwart, J. (2009). Coalescent Theory: An Introduction. *Systematic Biology*, *58*(1), 162–165. doi:10.1093/schbul/syp004

Silvestro, D., Cascales-Miñana, B., Bacon, C. D., & Antonelli, A. (2015). Revisiting the origin and diversification of vascular plants through a comprehensive Bayesian analysis of the fossil record. *New Phytologist*, *207*(2), 425–436. doi:10.1111/nph.13247

Strullu-Derrien, C., Selosse, M.-A. A., Kenrick, P., & Martin, F. M. (2018). The origin and evolution of mycorrhizal symbioses: from palaeomycology to phylogenomics. *New Phytologist*, *220*(4), 1012–1030. doi:10.1111/nph.15076

Tajima, F. (1983). Evolutionary relationship of DNA sequences in finite populations. *Genetics*, *105*(2), 437–460.

Tajima, F. (1989). The effect of change in population size on DNA polymorphism. *Genetics*, *123*(3), 597–601.

Varga, T., Krizsán, K., Földi, C., Dima, B., Sánchez-García, M., Sánchez-Ramírez, S., … Nagy, L. G. (2019). Megaphylogeny resolves global patterns of mushroom evolution. *Nature Ecology & Evolution*, *3*(4), 668–678. doi:10.1038/s41559-019-0834-1

Venice, F., Ghignone, S., Salvioli di Fossalunga, A., Amselem, J., Novero, M., Xianan, X., … Bonfante, P. (2020). At the nexus of three kingdoms: the genome of the mycorrhizal fungus Gigaspora margarita provides insights into plant, endobacterial and fungal interactions. *Environmental Microbiology*, *22*(1), 122–141.

doi:10.1111/1462-2920.14827

Watterson, G. A. (1975). On the number of segregating sites in genetical models without recombination. *Theoretical Population Biology*, *7*(2), 256–276. doi:10.1016/0040-5809(75)90020-9

Zanne, A. E., Tank, D. C., Cornwell, W. K., Eastman, J. M., Smith, S. A., FitzJohn, R. G., … Beaulieu, J. M. (2014). Three keys to the radiation of angiosperms into freezing environments. *Nature*, *506*(7486), 89–92. doi:10.1038/nature12872
